# Recurrent evolution of vertebrate transcription factors by transposase capture

**DOI:** 10.1101/2020.05.07.082677

**Authors:** Rachel L. Cosby, Julius Judd, Ruiling Zhang, Alan Zhong, Nathaniel Garry, Ellen J. Pritham, Cedric Feschotte

## Abstract

How genes with novel cellular functions evolve is a central biological question. Exon shuffling is one mechanism to assemble new protein architectures. Here we show that DNA transposons, which are mobile and pervasive in genomes, have provided a recurrent supply of exons and splice sites to assemble protein-coding genes in vertebrates via exon-shuffling. We find that transposase domains have been captured, primarily via alternative splicing, to form new fusion proteins at least 94 times independently over ∼350 million years of tetrapod evolution. Evolution favors fusion of transposase DNA-binding domains to host regulatory domains, especially the Krüppel-associated Box (KRAB), suggesting transposase capture frequently yields new transcriptional repressors. We show that four independently evolved KRAB-transposase fusion proteins repress gene expression in a sequence-specific fashion. Genetic knockout and rescue of the bat-specific *KRABINER* fusion gene in cells demonstrates that it binds its cognate transposons genome-wide and controls a vast network of genes and *cis*-regulatory elements. These results illustrate a powerful mechanism by which a transcription factor and its dispersed binding sites emerge at once from a transposon family.

**One Sentence Summary:** Host-transposase fusion generates novel cellular genes, including deeply conserved and lineage specific transcription factors.

## Main Text

The role of gene duplication has been extensively examined and has contributed to the birth and functional diversification of many genes (*1*). These include key developmental regulators such as *Hox* (*2*) and *Pax* genes (*3*), which have been associated with major biological innovations in animals such as the diversification of body plans and organs such as the eye. Although gene duplicates can evolve diverging developmental functions relative to their parental gene, their domain architecture and biochemical activities tend to remain the same (*2, 3*). Proteins with entirely novel biochemical functions can arise via gene duplication (*4, 5*), but this appears to be rare (*6*). While completely new proteins can occasionally evolve ‘*de novo*’ from previously non-coding sequences (*7, 8*), the most obvious path to form proteins with new functionality is the rearrangement of domains with pre-existing function into new composite architectures. Exon shuffling, the process by which new combinations of exons can be assembled through RNA splicing, has long been viewed as a powerful mechanism to create new protein architectures in eukaryotic evolution (*9*). While the process of exon shuffling is thought to account for the evolution of many new protein architectures (*10*-*12*), the source of new exons and the mechanisms by which they get assimilated have been scarcely characterized. Here we investigate the role of DNA transposons as a source of raw material for the birth of novel proteins via exon shuffling.

DNA transposons encode transposase proteins, which recognize and mobilize DNA through direct sequence-specific interaction with their cognate transposons (*13*). The canonical architecture of transposase proteins consists of a DNA-binding domain and a catalytic nuclease domain. Both domains may be repurposed or ‘domesticated’ for cellular function (*13*). Moreover, the very mobility of DNA transposons may facilitate exon-shuffling by inserting these functional domains into new genomic contexts, where they could then be spliced to host domains to generate novel host-transposase fusion (HTF) genes. Three genes clearly born via transposase capture have been documented previously, including one specific to placental mammals (*GTF2IRD2*; (*14*)) and two specific to primates (*SETMAR* (*15*) and *PGBD3-*CSB (*16*)). A similar scenario has been proposed to explain the origin of the Paired DNA-binding domain of the *Pax* family of transcription factors (TFs), though the deep ancestry of *Pax* genes, which coincides with the emergence of metazoans, has obscured the precise steps by which these factors evolved (*13, 17*). Further, the extent of transposase capture, the mechanisms facilitating it, and the functions of the resulting genes often remain unclear. Here we show that transposase capture has been a widespread mechanism for gene birth of new genes during vertebrate evolution, and that this process is prone to creating lineage-specific transcription factors (TFs).

### Transposase capture is a pervasive mechanism for novel gene formation in tetrapods

To identify host-transposase fusion (HTF) genes, we surveyed all tetrapod gene annotations (NCBI Refseq; Table S1) predicted to encode proteins with at least one domain of transposase origin (Pfam, Table S2) fused in-frame to a host-derived protein sequence (Conserved Domain Architecture Retrieval Tool [CDART] (*18*)). We also required RNA-sequencing evidence supporting all annotated exon/intron junctions (see Methods). To trace the evolutionary origin of each HTF gene, we searched for syntenic orthologs as well as paralogs across all vertebrate genomes available in the NCBI Refseq database (see Methods). This analysis yielded 106 unique HTF genes originating from 94 independent fusion events and 12 subsequent duplication events across the 596 species examined (Fig. 1; Table S3).

**Fig. 1:**
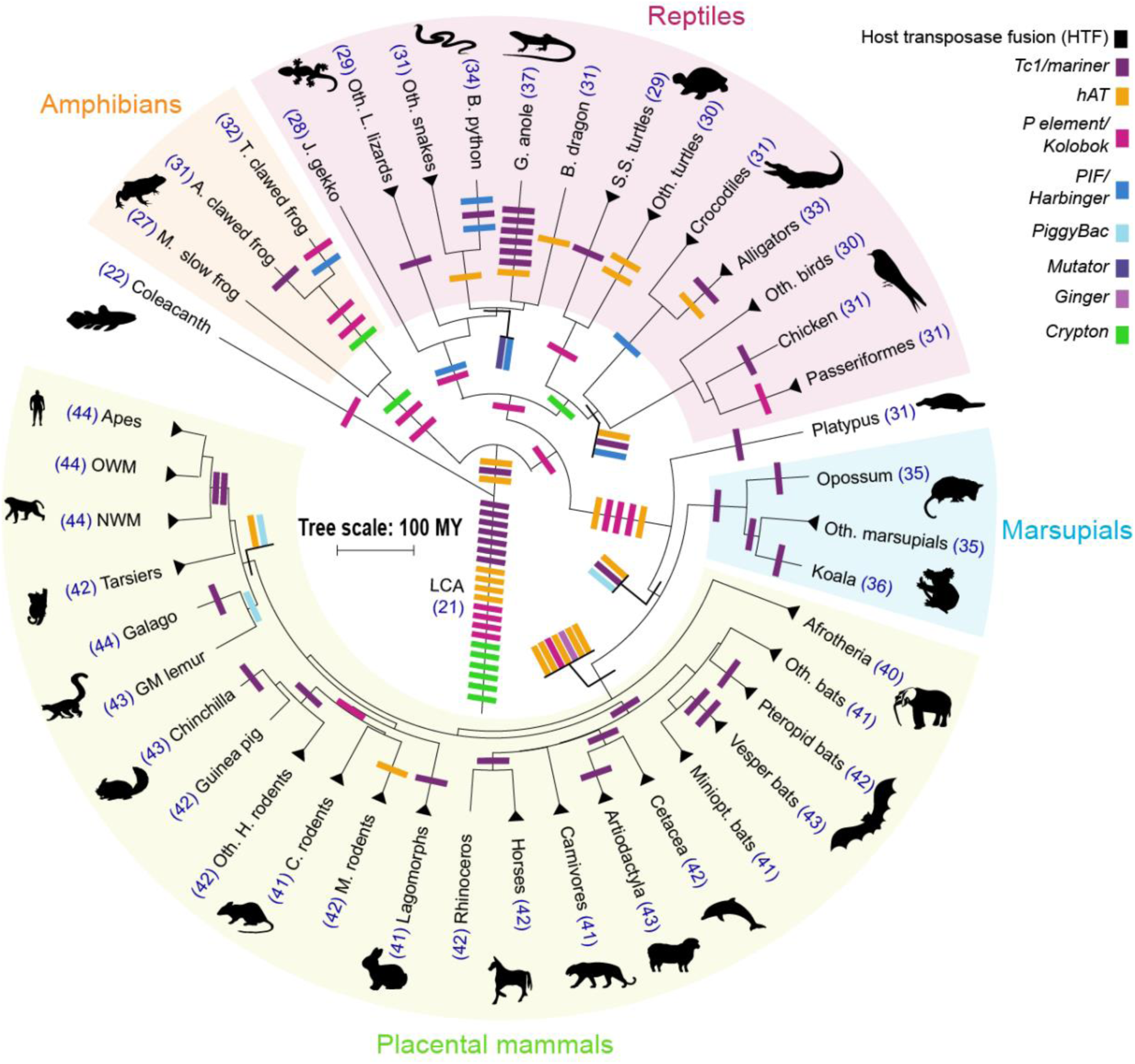
Gene birth by transposase capture is pervasive in tetrapods. Tetrapod phylogenetic tree with boxes representing HTF fusion genes. Color indicates transposase superfamily assimilated. OWM=Old world monkeys; NWM=New world monkeys; GM=Gray mouse; H=Hystricoid; C=Castorid, M=Muroid, Miniopt=Miniopterid, Vesper=Vespertilionid, S.S=soft-shelled; B=bearded dragon; G=Green; B=Burmese python; L=Lacertid; J=Japanese; T=Tropical; A=African; M=Mountain; LCA=last common ancestor; MY=million years.

Placing fusion genes onto the species phylogeny suggest that they have evolved continuously during evolution (Fig. 1). Some fusion events (11.6%) preceded the divergence of tetrapods (>350 million year ago, Mya), while others arose much more recently as they were conserved across relatively small species lineages (<5 species, 26.1%), or found in a single species (21%) (Fig. 1; Table S3). Several species lineages experienced multiple HTFs of recent origins, as seen in the green anole (n=6), the Burmese python (n=3), the tropical clawed frog (n=2), and the vespertilionid bats (n=2), consistent with recent episodes of DNA transposon activity documented in these lineages (Fig. 1; (*19*-*23*)). Mammals generally have more HTF genes (mean = 40.8 ± 3.55) than other clades (Reptiles: mean = 29.3 ± 1.53; Amphibians: mean = 30 ± 2.65), reflecting apparent bursts of HTF evolution in mammalian (5.3% of all events), therian (3.2%) and eutherian (8.4%) ancestors. All known major eukaryotic DNA transposon superfamilies contribute to HTFs (Fig. 1), but Tc1/*mariner* (36.1%), hAT (23.4%), and P element/Kolobok (21.3%) transposases predominate, which mirrors the success of these superfamilies throughout tetrapod evolution (*13, 24*).

To validate that the transposase coding region of each HTF gene has evolved under functional constraint, we performed codon selection analysis on the transposase domain of each HTF shared by two or more species separated by >50 million years (My) of divergence. All tested HTFs (n= 81) display signatures of purifying selection on their transposase domains (*dN/dS <1*, *p <0.05*, LRT, Table S3), supporting their domestication for organismal function.

To further assess the functional capacity of HTF genes, we used publicly available data to examine the RNA expression patterns across 54 tissues for 44 HTFs present in the human genome (GTEx portal). Each of the genes were expressed (transcripts per million, TPM >1) in at least one human tissue, and could be classified into three categories: lowly expressed broadly (TPM >1 in 80% of the tissues, n=23), highly expressed broadly (TPM >10 in 80% of tissues, n=12), and tissue-restricted (TPM >1 in <20% tissues, n=9) (Fig S1). These data suggest that most human HTF genes are broadly expressed and may function in a variety of contexts. Collectively, these data demonstrate that HTF has been a recurrent mechanism for the generation of novel cellular genes in tetrapod evolution, including at least 44 HTFs in humans.

### Transposase capture occurs through alternative splicing

To illuminate the mechanism by which transposase domains are captured to form new chimeric proteins, we examined the gene structure of HTFs. In all cases, the transposase-derived domains are encoded by exons distinct from the host domains, suggesting that transposase capture occurred via splicing events. To further delineate the process, we investigated in detail the birth of *KRABINER,* a recently-evolved HTF we identified in vespertilionid bats. *KRABINER* is predicted to encode a 447-amino acid protein consisting of an N-terminal Krüppel-associated box (KRAB) domain fused to a full-length *Mlmar1 mariner* DNA transposase (Fig. 2A). Using a combination of comparative genomics, PCR, and RT-PCR (see Methods), we inferred that *KRABINER* originated in the common ancestor of the nine vespertilionids examined, but after their divergence from miniopterids, ∼45 Mya, through the following steps: i) *mariner* insertion into the last intron of *ZNF112*, a gene present in all eutherian mammals; ii) alternative splicing to the upstream exons of *ZNF112* using a splice acceptor site pre-existing in the ancestral *Mlmar1 mariner* transposon; and iii) a unique single nucleotide deletion in the transposase coding sequence which generated a single open reading frame coding the chimeric protein (Fig. S2, Fig. 2B). This sequence of events is reminiscent of the process that gave birth to *SETMAR* (*15*) and *PGBD3-CSB* (*25*) in the primate lineage, and suggests that DNA transposons possess features that facilitate their capture via alternative splicing.

**Fig. 2:**
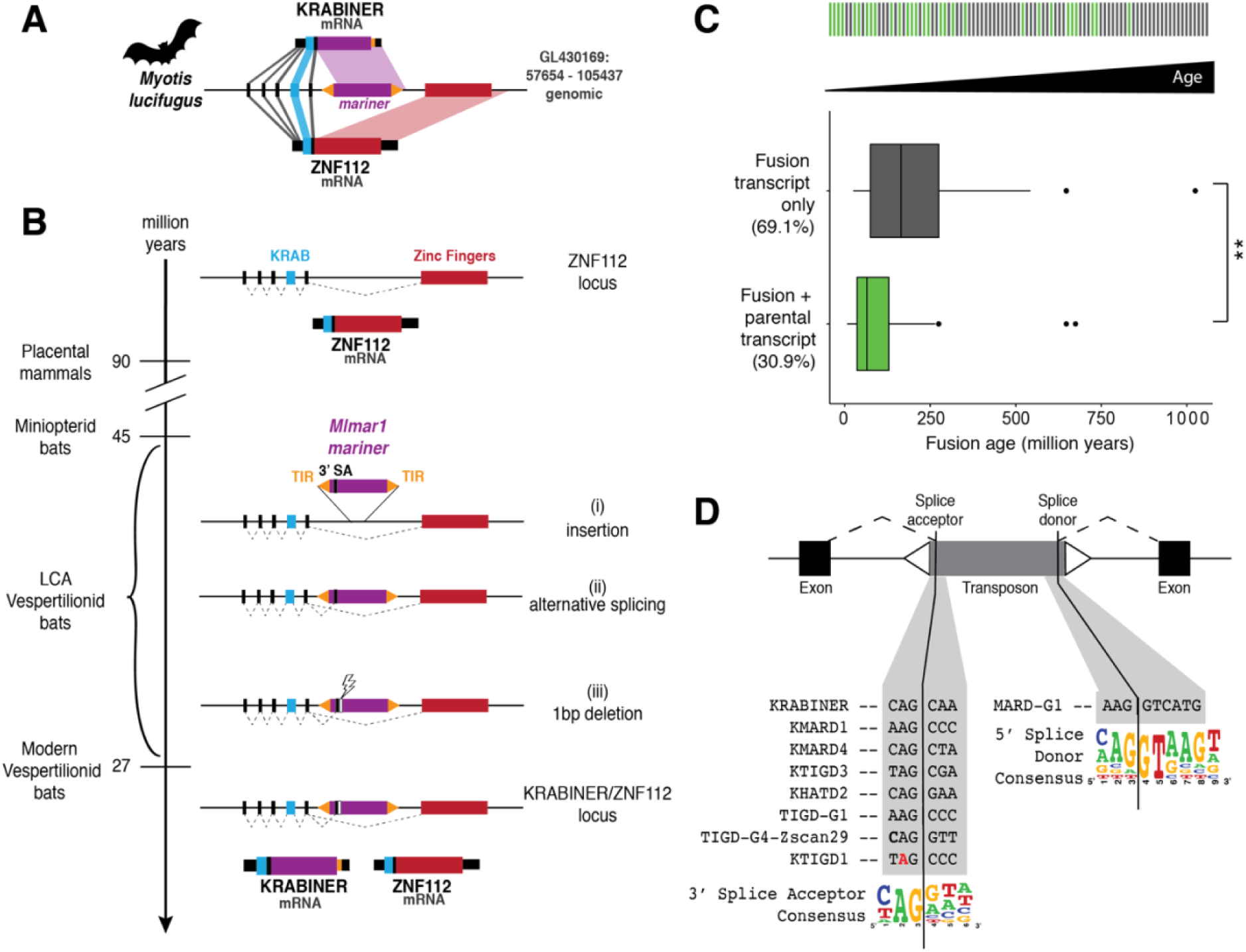
Transposase capture by alternative splicing. A) *ZNF112/KRABINER* locus in vespertilionid bats. B) Steps required for *KRABINER* birth. C) Age of fusion genes with (green) or without (gray) evidence for alternative splicing. Fusion age (bottom) determined by the midpoint of age range for each fusion as described in Table S3; top shows qualitative illustration of host transcript loss over time. D) Summary of transposon splice site usage for 9 HTFs, with canonical mammalian splice sites shown as a sequence logo. Red denotes nucleotides in the splice site that diverge from the transposon consensus sequence. SA=splice acceptor, LCA=last common ancestor. ** *p<0.01* 2-sample Wilcoxon Test.

To assess whether this mechanism is generalizable, we surveyed all HTF gene models for evidence of alternative splicing. For most of the young HTFs (18 out of 33 HTFs <100 My old) we found unequivocal evidence for the co-existence of both fusion and parental gene transcripts, but as HTFs became older only the fusion transcript was generally detected (Fig. 2C; Table S3). These findings suggest that most HTFs are born as alternatively spliced variants of an ancestral gene, but over time the HTF transcript often becomes the primary or sole transcript for that gene. Thus, alternative splicing is a prominent mechanism for the assimilation of transposase domains by the host proteome.

The splice site enabling the capture of *KRABINER*’s transposase was provided by the ancestral *Mlmar1 mariner* transposon (Fig. S2C). To investigate whether this is a more general principle, we selected eight additional HTFs derived from recent transposon families to trace the origin of the splice site used for transposase capture. For each family, we generated a majority-rule consensus sequence, which serves as a proxy for the ancestral transposon (Data S1). In all cases, we found that the splice site was directly derived from the ancestral transposon sequence. For 6 of out 8 HTFs, the splice site sequence was strictly identical to that of the consensus sequence, while in the remaining two the splice site differed from the consensus by a single substitution (Fig. 2D). Importantly, in 8/9 cases examined, the splice site is an acceptor sequence immediately upstream of the transposase ORF. These results suggest that DNA transposons are predisposed for exon shuffling because they carry splice sites positioned to facilitate transposase expression.

### Fusion of transposase DNA-binding domains to KRAB is prevalent

To explore the cellular function of HTFs, we first characterized their protein domain architecture and composition (Fig. 3A, Fig. S3). Amongst transposon-derived domains, DNA-binding domains predominate (76.5%; Fig. S3), though some HTFs also include catalytic or accessory transposase domains (Fig. S3). Among host domains (i.e. not normally found in transposases), we identified 55 distinct conserved domains, most of which (76%) were involved in a single fusion event (Fig. 3A). Several of the host domains are predicted to function in transcriptional and/or chromatin regulation, such as the KRAB, SET, and SCAN domains (*26*-*28*). KRAB was by far the host domain most frequently fused to transposase: we inferred this domain to have been involved in 32 independent fusion events across the phylogeny, accounting for approximately one third of all HTFs (Fig. 3A). KRAB domains are abundant in tetrapod genomes and most commonly found in KRAB-Zinc Finger proteins (KRAB-ZFPs), an exceptionally diverse family of transcription factors (>200 genes in most tetrapod genomes; 487 in humans) (*29*). However, the prevalence of KRAB-transposase fusions is unlikely to be solely explained by the genomic abundance of KRAB-ZFP genes because (i) other equally expansive gene families are not involved in HTF (e.g. olfactory receptors, ∼350 genes in humans; (*30*)) and (ii) we still find two independent KRAB-transposase fusions in bird genomes despite their paucity in KRAB-ZFPs (∼8 genes per genome) (*29*). These observations suggest that the combination of KRAB and transposase is preferentially retained by natural selection.

**Fig. 3:**
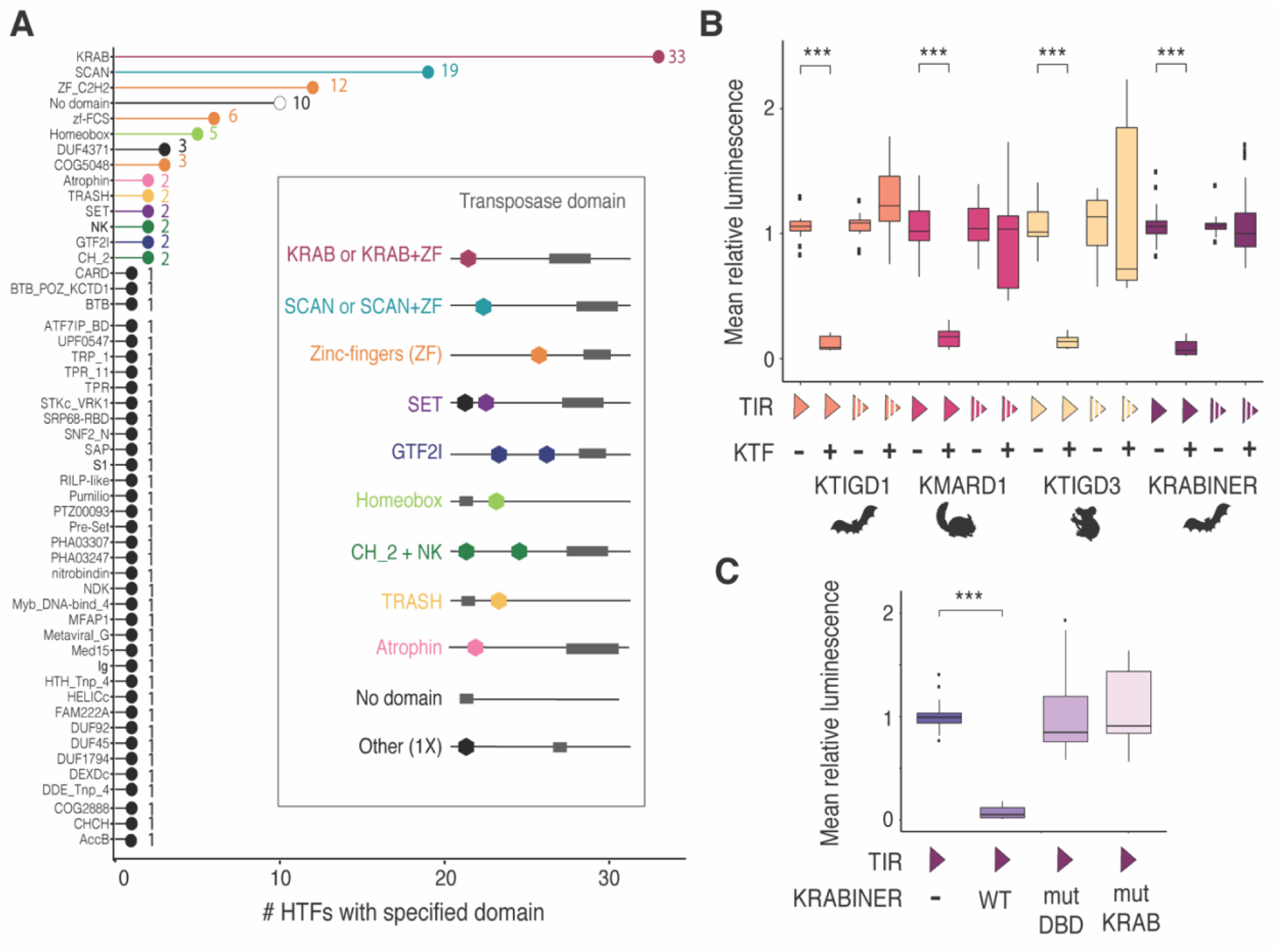
Biochemical activities of host-transposase fusion proteins. A) Diverse host domains are fused to transposases. X-axis specifies the number of HTF genes a given domain is present in; some fusions contain more than one domain. Inset shows representative domain architecture schematic for select host-transposase fusions. B) KRAB-transposase fusions repress gene expression in a sequence-specific manner. C) KRABINER requires both its KRAB and DBD domains to repress gene expression. Y axes in B-C boxplots correspond to mean luminescence relative to the KTF (-) state for each comparison. KTF=KRAB-transposase fusion; TIR=terminal inverted repeat; filled triangle = consensus TIR, interrupted triangle = scrambled TIR; +/- = presence/absence of respectively; *** *adj. p<0.001*; 2-sample Wilcoxon Test, Holm-Bonferroni correction.

### KRAB-transposase fusions act as sequence-specific repressors of gene expression

Given the prevalence of KRAB-transposase fusions and the canonical function of KRAB domains in establishing silent chromatin when tethered to DNA (*26*), we next used these genes as a paradigm to test the hypothesis that transposase fusion creates novel sequence-specific transcriptional regulators. We selected four recently emerged KRAB-transposase fusions for which we had generated consensus sequences of their cognate transposon family, enabling us to identify their terminal inverted repeats (TIRs) which typically contain the transposase binding site (Fig. S4). We cloned the consensus sequence of each TIR or a scrambled version upstream of a firefly luciferase reporter and measured luciferase expression in HEK293T cells in the presence or absence of a vector expressing the cognate HTF protein. Each KRAB-transposase fusion protein strongly repressed luciferase expression in the presence of its cognate intact TIR but not the scrambled sequence (Fig. 3B). These results indicate that KRAB-transposase proteins can repress gene expression in a sequence-specific manner.

To test whether KRAB-transposase repression is dependent on KAP1 (TRIM28), the transcriptional corepressor often recruited by the KRAB domain (*26*), we repeated the reporter assays in HEK293T cells knockout (KO) for *KAP1* (*31*). The results (Fig. S5) show that repression by KMARD1 and KTIGD1 is dependent on KAP1, whereas KRABINER and KTIGD3 are only partially dependent on KAP1. To further dissect the requirement of individual domains, we generated two mutant versions of KRABINER by altering residues predicted to (i) compromise DNA-binding activity (mutDBD) or (ii) the function of the KRAB domain (mutKRAB). To generate the DBD mutant, we leveraged the close similarity of KRABINER’s *mariner* transposase to that of *Mos1* (*22*), a well-characterized transposon from *Drosophila*. Previous studies demonstrated that a single point mutation in the first helix-turn-helix motif of the *Mos1* transposase was sufficient to abolish binding to its TIR (*32*). We mutated the homologous site in *KRABINER’s* DBD, as well as three additional residues shown to directly contact TIR DNA in the *Mos1* transpososome crystal structure (*33*) (Fig. S4B). To generate the KRAB mutant, we introduced several point mutations altering conserved residues previously identified as critical for KRAB-mediated repression (*34*-*37*). While the mutDBD and mutKRAB proteins were expressed at comparable levels as wild-type (WT) KRABINER (Fig. S4C), both lost their ability to repress reporter gene expression (Fig. 3C). Together the results of these reporter assays support the hypothesis that KRAB-transposase fusions yield modular proteins functioning as sequence-specific transcriptional repressors.

### KRABINER regulates transcription in bat cells

To further test whether transposase capture gives birth to transcriptional regulators, we investigated the ability of KRABINER to modulate gene expression in embryonic fibroblasts of the bat *Myotis velifer,* where the gene is endogenously expressed (Fig. S2). We used the CRISPR-Cas9 system to engineer a *KRABINER* KO cell line with a pair of gRNAs designed to precisely delete the *mariner* transposon from the *ZNF112* locus, leaving the rest of the gene intact (Fig. 4A; Fig. S6). We then used a *piggyBac* vector to deliver transgenes at ectopic chromosomal sites into the KO cell line to establish independent clonal lines reintroducing wild-type *KRABINER* (WT, n=4 cell lines), or the predicted DNA-binding mutant (mutDBD, n=3), or the predicted KRAB mutant (mutKRAB, n=3) (Fig. S6). Each transgene was cloned under the control of a tetracycline-inducible promoter and contained a C-terminal *myc* tag to monitor protein expression (Fig. 4A; Fig. S7). The non-induced condition showed leaky expression more closely recapitulating the level of WT *KRABINER* transcription (hereafter termed “rescue”, R), while transgene induction resulted in *KRABINER* over-expression (OE) relative to the parental cell line (Fig. S8A).

**Fig. 4:**
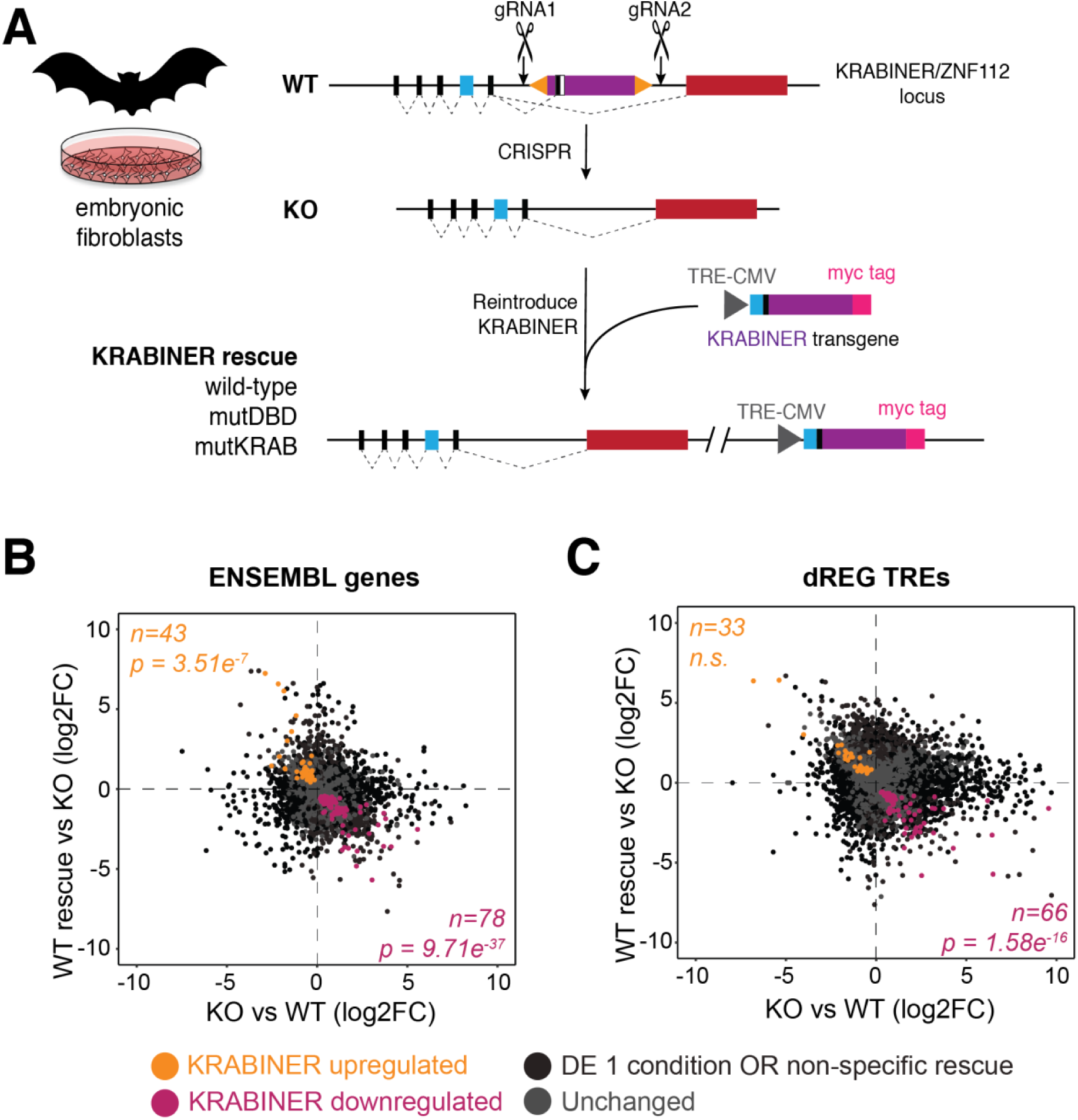
KRABINER regulates transcription of genes and TREs in bat cells. A) Strategy to generate KRABINER KO and rescue lines. TRE=tet responsive element; CMV=cytomegalovirus. B-C) Summary of transcriptional changes of genes and TREs, respectively, upon loss and restoration of KRABINER. KRABINER regulated genes (up or down) change reciprocally between KO vs WT and WT KRABINER rescue vs KO comparisons. *p* values calculated via a right-tailed hypergeometric test. DE 1 condition refers to differential transcription in either the KO vs WT or WT KRABINER vs KO comparison. Non-specific refers to a gene rescued by WT KRABINER and one or both mutDBD and mutKRAB variants. Unchanged refers to genes/TREs with *adj. p*>0.05.

To investigate whether KRABINER modulates transcription, we profiled *KRABINER* KO and WT cells with Precision Run-On followed by sequencing (PRO-Seq), a technique that provides a sensitive measurement of nascent transcription throughout the genome, including genes bodies and transcribed regulatory elements (TREs) such as promoters and enhancers (*38, 39*). We identified 2,644 genes differentially transcribed between WT and KO cells: 1,295 were upregulated in KO (UP), 1,349 were downregulated (DOWN) (DESeq2; *adj. p* <0.05; (*40*)), suggesting KRABINER is capable of regulating genic transcription. To identify transcriptional changes which require both the DNA-binding domain and KRAB activity of *KRABINER,* we also assessed transcriptional changes in the transgenic rescue lines. Of the 2,644 altered genes, 121 genes (43 UP, 78 DOWN) had their transcription level consistently restored in WT transgenic lines but not in any of the mutant transgenic lines (Fig. 4B; Table S4; overlap *p<0.001*, right-tailed hypergeometric test). A similar pattern was observed for TREs (identified using dREG; (*39*)), with 3,472 differentially transcribed TREs following loss of KRABINER, of which 99 were restored exclusively in the WT lines (33 UP, 66 DOWN; Fig. 4C; Table S5; overlap *p<0.001* for downregulated TREs, right-tailed hypergeometric test). A subset of these TREs are associated with restored gene body transcription (18% UP, 12% DOWN), while others are distal (>100 kb) to genes (18% UP, 33% DOWN) or associated with genes bodies that are not differentially transcribed (64% UP, 55% DOWN) (Table S4). While our reporter assays indicate that KRABINER can act as a strong repressor, our loss of function analyses in bat cells suggest that the protein exerts a range of transcriptional modulation on the bat genome.

In addition to transcriptional changes unique to the WT transgenic lines, there were several genes and TREs that were rescued by the WT transgene and either mutDBD (100 genes, 79 TREs) or mutKRAB transgenes (73 genes, 65 TREs; Fig. S9). Thus, while at some loci KRABINER’s regulatory activity appears to require both its DNA-binding and KRAB domains, its transcriptional effects on other loci only requires one of these domains. Such mechanisms may also explain the ability of KRABINER to either activate or repress transcription in a locus-dependent fashion. Taken together, this data suggests that KRABINER contributes to the transcriptional regulation of a subset of genes and cis-regulatory elements in the examined embryonic cell line.

To investigate whether KRABINER regulates a discrete set of genes or a network of related genes, we performed gene ontology enrichment analysis for all genes rescued by the WT transgene (n=121) as well as additional target genes whose promoters were found to be differentially transcribed in the TRE analysis (n=57) (Table S4; ClueGO, see Methods; (*41*)). This analysis revealed a significant enrichment for genes involved in negative regulation of cell migration (GO: 0030336, *adj. p=*2.1e^-6^) and in gastrulation (GO: 0007369, *adj. p=*2.5e^-5^) as the most enriched terms (Fig. S10A; Table S5). Additional significant terms were linked to morphogenetic or developmental pathways, including positive regulation of Wnt signaling pathway (GO: 2000096, a*dj. p=*5.2^e-3^), artery morphogenesis (GO:0048844, *adj. p=*2.5^e-2^), heart valve morphogenesis (GO: 0003179, *adj. p=*8.8e^-3^), and neural crest differentiation (GO: 0014033, *adj. p=*2.2e^-2^) (Fig. S10A; Table S5; 2-sided hypergeometric test, Bonferroni step-down). A similar result was obtained for the downregulated genes alone, suggesting KRABINER’s direct targets may be downregulated (Fig. S10B; Table S5). These results suggest that, in the embryonic cell line examined, KRABINER regulates a set of genes enriched for developmental functions, as may be expected for a canonical transcription factor.

### KRABINER binds to many genomic sites, with a preference for *mariner* TIRs

We next use chromatin immunoprecipitation followed by sequencing (ChIP-seq) to determine if and where KRABINER binds throughout the bat genome. We profiled binding of myc-tagged KRABINER WT protein in the KO cell line background as well as that of mutKRAB and mutDBD mutant proteins 24-hour post induction of transgene expression (*n=3* each) (Fig. S11). With these samples, we called binding peaks using MACS2 (*42*) for each genotype relative to input and filtered out peaks identified in all three genotypes (>50% reciprocal overlap), which are likely to represent spurious or non-specific interactions (see Methods). This analysis identified 1888 WT, 5702 mutKRAB, and 4264 mutDBD peaks respectively. The higher number of peaks obtained with mutKRAB and mutDBD likely reflects the higher expression of those transgenes relative to the WT transgene (Fig. S8). In order to identify genomic sites likely bound directly via KRABINER, we focused on the WT peaks and used the mutKRAB and mutDBD peaks as additional filters or background sets.

Of the 1888 WT KRABINER binding peaks, 56% (n=1070) were private to the WT condition, suggesting that most of KRABINER’s binding requires both functional domains (Fig. 5A). However, there were about twice as many peaks shared between the WT and mutKRAB conditions (n=572, 28%) than the WT and mutDBD (n=291, 15%) (>50% reciprocal overlap; Fig. 5A, Fig. S12), suggesting that most of KRABINER’s binding appears dependent on its DBD. The set of peaks overlapping exclusively between WT and mutDBD conditions likely represent indirect genomic interactions, perhaps mediated through protein-protein interactions via the KRAB domain (*43*) or other region of the protein.

**Fig. 5:**
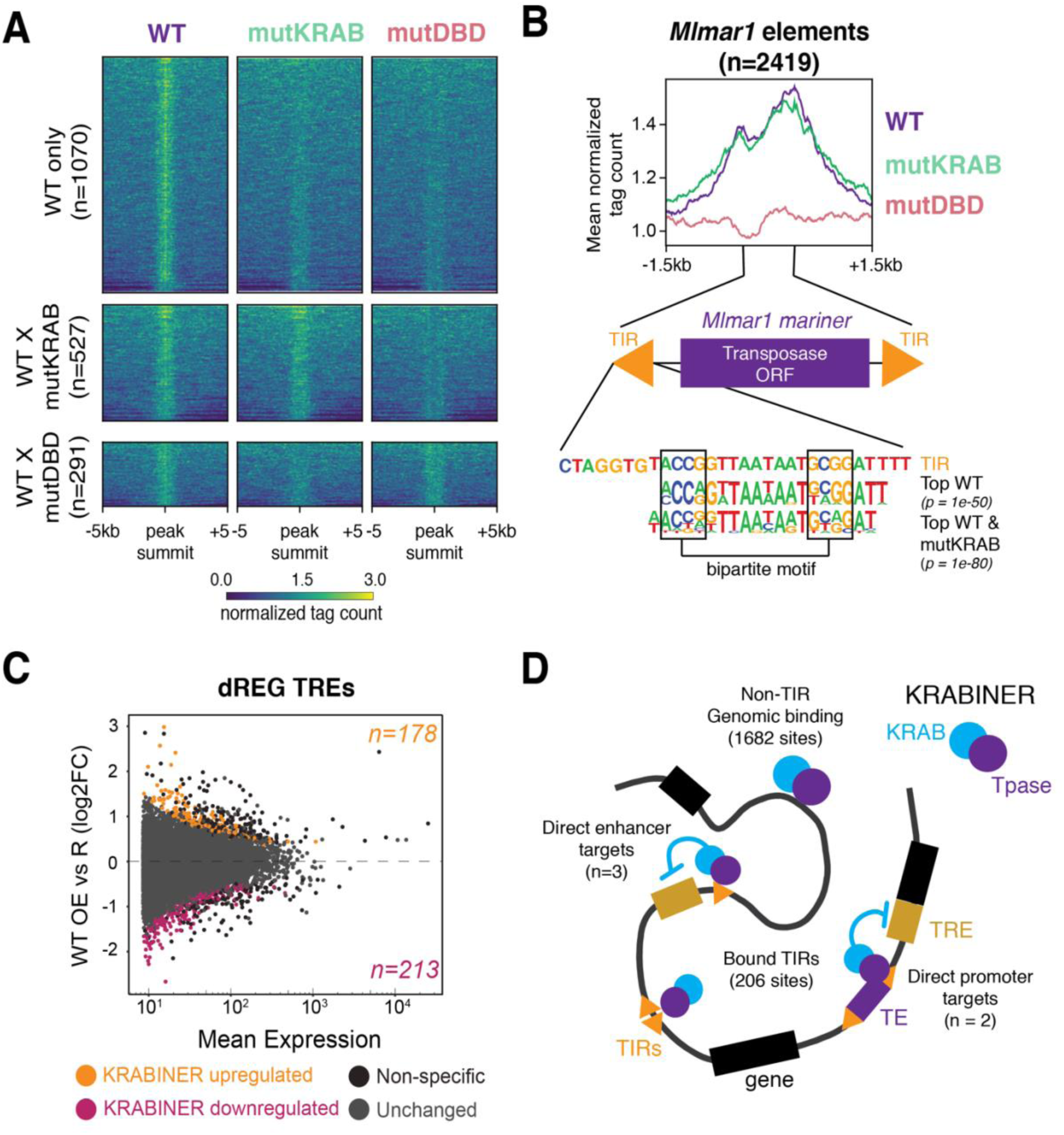
KRABINER binds to *mariner* TIRs in bat cells. A) Heatmaps summarizing merged, library-size and input-normalized ChIP-seq coverage of each KRABINER variant centered on the summit of WT (top), WTXmutKRAB (middle), and WTXmutDBD (bottom) peak sets. B) Metaplot summarizing normalized ChIP-seq coverage of each KRABINER variant over all genomic *Mlmar1* elements (top). The top enriched motif in the WT and WT & mutKRAB peak sets is identical to the predicted bipartite binding motif within the *Mlmar1 mariner* TIR (bottom) (HOMER). C) MA plot summarizing changes in TRE transcription upon over-expression of WT KRABINER. Non-specific (black) refers to changes in TRE transcription that are shared between over-expression of WT KRABINER and one or both mutant KRABINER variants. Unchanged (gray) refers to TREs with *adj. p*>0.05. D) Proposed model for KRABINER’s function as a transcription factor in bats. KRABINER directly binds to *mariner* TIRs within the genome, leads to direct downregulation of a subset of TREs. KRABINER also binds to other genomic regions, and indirectly regulates a number of genes and TREs. Resc. = rescue; OE = overexpression; WT= purple; mutDBD=pink; mutKRAB=green. WT = wild-type, DBD = DNA binding domain, ORF = open reading frame, TIR = terminal inverted repeat.

We then asked if there were any sequence motifs that might explain KRABINER binding to its genomic sites (HOMER; (*44*)). We extracted a 200-bp window centered on the peak summit for all WT peaks and asked which *de-novo* motifs were enriched relative to mutDBD peaks for the WT and WTxmutKRAB peak sets or to the mutKRAB peaks for the WTxmutDBD peak set. For the WT private and WT-mutKRAB peaks, the top enriched motif *(p=*1e^-50^ and *p=*1e^-80^) was identical, and strongly resembled a bipartite region within the *Mlmar1 mariner* TIR sequence that aligns with the binding sites mapped previously for the *Mos1* transposase ((*45*); Fig. S12; Fig. 5B). Of the additional 12 and 17 enriched motifs predicted for the WT and WT-mutKRAB peak sets respectively, all were derived from *Mlmar1* elements (Fig. S12). These data suggested that the WT and mutKRAB proteins bind many *Mlmar1* elements dispersed throughout the genome. Consistent with this, *Mlmar1* elements are strongly enriched within WT (log2 Fold enriched [FE] = 4.43, *p=*2.2^e-16^) and WT-mutKRAB peaks (log2FE = 4.95, *p=*2.2^e-16^), but not the WT-mutDBD peaks (*p*=0.63) (Fig. S12, 2-sided binomial test, 1000 bootstraps, see Methods). In total, the WT KRABINER transgenic protein binds to 206 (8.5%) of all *Mlmar1* elements annotated in the bat genome assembly.

To determine where in the *Mlmar1* element KRABINER binds, we plotted the input-normalized ChIP-seq reads for each genotype over all *Mlmar1* transposons in the genome. We found that a substantial fraction of these transposons was bound by the WT and mutKRAB proteins, but not the mutDBD protein, consistent with the peak-based approach (Fig. 5B; Fig. S12). The ChIP-seq read coverage peaks within the TIR regions, and especially the 3’ TIR (Fig. 5B; Fig. S12), which is consistent with the binding activity of other *mariner* transposons (*32, 33, 46*). Collectively, our ChIP-seq data demonstrates that KRABINER is capable of binding numerous genomic sites, with a preference for *Mlmar1* TIRs.

### KRABINER binding is associated with downregulation of nearby TREs

We next asked if KRABINER genomic binding was associated with transcriptional change. To do this, we induced expression of the KRABINER transgenes and performed PRO-seq 24-hours post induction (OE), conditions matching our ChIP-seq experiments. We then identified TREs differentially transcribed between the OE vs rescue conditions, which are of the same exact genotype and in principle differ only in the level of KRABINER expression (Figs. S7-8). Because TREs represent discrete transcriptional units such as promoters and enhancers, we reasoned that KRABINER binding to or near these regions would more likely impact transcription than binding within a gene body. We identified 391 TREs (178 UP, 213 DOWN; Fig. 5C) differentially expressed upon overexpression of the WT KRABINER protein but neither of the mutant proteins (mutDBD or mutKRAB). Additionally, several TREs were differentially expressed in the same direction upon OE of WT protein and either mutDBD or mutKRAB (Fig S13), consistent with the hypothesis that a subset of KRABINER’s transcriptional changes require only one of its functional domains.

To determine if KRABINER binding is associated with differential TRE transcription, we first examined whether WT KRABINER ChIP-seq peaks were located near (<1 kb) differentially-expressed TREs. While the total number of KRABINER peaks located nearby differentially expressed TREs was small (n=6), it was a significant enrichment over the random expectation (log2FE = 2.58, empirical *p=0,* 10000 bootstraps, see Methods) (Fig. S14). Interestingly, all were downregulated TREs, consistent with the results of our reporter assays suggesting that tethering KRABINER to DNA induces local transcription repression. Furthermore, five out of six differentially expressed TREs were located within ∼1-kb of at least one *Mlmar1* element, and the TIRs of these transposons were located near the summit of the KRABINER peaks. Finally, several of the TREs connected with KRABINER binding to nearby *Mlmar1* elements were also associated with changes in adjacent gene expression.

For example, a differentially expressed TRE is located in the promoter region of the bat ortholog of the DNA damage recognition and repair factor gene *XPA*, which is downregulated (log2FC = −0.69, *adj. p* = 0.0079) upon WT KRABINER OE and located immediately adjacent to a *mariner* TIR bound by KRABINER (Fig. S15). A similar pattern is seen for an intergenic enhancer element, located between the bat homolog of the family with sequence similarity 174 member B (*FAM174B*) and chromodomain helicase DNA binding protein (*CHD2*) genes (Fig. S16). This region contains three distinct TREs, two of which are downregulated upon WT KRABINER OE (log2FC = −1.1, *adj. p=*0.04 and log2FC = −1.09, *adj. p=*0.006 respectively), and this change is associated with KRABINER binding to the *Mlmar1* TIRs immediately upstream of these TREs (Fig. S16). Other regulated TREs include the promoter of the bat homolog of the small nuclear ribonucleoprotein polypeptide A’ (*SNRPA1*) gene (log2FC = −0.84, *adj. p*=0.02), which is located downstream of four bound *Mlmar1* elements (Fig. S17), and a distal enhancer (log2FC = −1.24, *adj. p*=0.01) upstream of the nucleoporin 50 (NUP50) gene (Fig. S18). Importantly, each of the KRABINER-downregulated TREs are near two or more bound TIR sequences, suggesting KRABINER’s effect on TREs may be strengthened by additional binding sites.

## Discussion

While gene birth via duplication has been extensively documented, how novel protein architectures and biological functions are born has remained poorly characterized. In this study, we validate the long-standing prediction that exon-shuffling is a major evolutionary force generating genetic novelty (*9*) and provide compelling evidence that DNA transposons are potent sources of raw material for this process. DNA transposons contribute not only DNA-binding and nuclease domains but also, in many cases, the splice sites that facilitate their capture into host-transposase fusion genes. By characterizing the evolution of HTF genes, we derived first principles of how host-transposase fusion occurs. We found that HTF genes are born in 3 general steps: i) insertion of the DNA transposon into an existing host gene, ii) fusion via alternative splicing to the host gene using splice sites provided by the transposon, and iii) subsequent selection for maintenance of the fusion gene, sometimes at the expense of the parental transcript. Additionally, we found that in some cases, such as the *Pax* genes, the HTF repertoire can be further expanded via gene duplication. These principles illuminate how transposon-mediated exon-shuffling occurs and provide a foundation for the identification of host-transposase fusion genes in other lineages.

Our work elucidates the mechanism by which several proteins with crucial developmental functions originated in evolution. Perhaps the most evocative example of this is *Pax6,* which controls eye development and patterning across animals (*47*). The emergence of *Pax* genes is widely viewed as a key step in the early evolution of eyes, a spectacular innovation of metazoans (*47*). Mutations in another HTF gene expressed predominantly in the brain, *POGZ*, have been repeatedly associated with autism and intellectual disability in humans (*48, 49*). While these are example of HTFs with relatively deep evolutionary origins and likely serving broadly conserved functions, our analysis indicates that the process of gene birth via transposase capture has been a continuous source of lineage-specific innovation. For instance, *KRABINER* is a gene specific to vespertilionid bats which appears to regulate, at least in cell culture, a subset of genes involved in developmental processes. Further experiments are needed to determine whether these findings apply *in vivo* and whether KRABINER regulates a process unique to the biology of these bats. Overall, our study shows that host-transposase fusion is not an anecdotal phenomenon, but rather a pervasive phenomenon fueling biological novelty in vertebrates, with a propensity for spawning new transcription factors.

Our study also provides clues as to why transposase capture appears predisposed to generate novel transcriptional regulators. One key lies in the fact that transposons not only encode DNA-binding proteins, but their mobilization and amplification throughout the genome provides an inherent mechanism to disperse a large pool of cognate binding sites. Thus, it is easy to envision how the capture of a transposase DNA-binding domain could provide a facile path for the emergence of a new cis-regulatory network (*50*). Historically this model has been difficult to test, however, because previously recognized transposase-derived transcription factors (such as *Pax*) and their network of regulated genes evolved hundreds of millions of years ago, which would have obscured the transposon origin of their genomic binding sites. Recent studies of two young HTF genes, *SETMAR* (40-58 my old; (*15, 51*)) and *PGBD3-CSB* (>40 my old; (*25, 52, 53*)) have begun to provide evidence in support of the network model. Our study of *KRABINER,* another young HTF, further bolsters and generalizes this model. We provide strong evidence that KRABINER acts as a canonical transcription factor, and that at least some of its transcriptional regulatory activity in bat cells is directed by KRABINER binding to its cognate *mariner* TIRs. However, only a minority of genes regulated by KRABINER appear to be direct targets, which suggests that KRABINER is integrated within a complex transcriptional network (Fig. 5D). Collectively, our results are consistent with the notion that transposase capture facilitates the wiring of new cis-regulatory circuits.

Although we focused on the tetrapod lineage in this study, we propose that the principles and implications of transposase capture revealed herein extend beyond vertebrates. Transposases are ancient and possibly the most abundant and ubiquitous genes in nature (*54*) and a variety of host domains that regulate transcription exist in all branches of the tree of life. It is easy to envision how these sequences have provided the raw material for the assemblage of endless combination of transposase-host fusion proteins throughout evolution.

## Acknowledgements

We acknowledge David Ray, Woody Wright, Nels Elde, John Lis, Helen Rowe, Joanna Wysocka, and Todd Macfarlan for providing reagents. We also thank members of the Feschotte lab for helpful discussion.

## Author Contributions

RC, EP, and CF conceived of and designed the study. RC and RZ identified and analyzed host-transposase fusion genes. RC and AZ performed luciferase reporter assays. RC generated KRABINER KO cell lines and RC and NG generated the KRABINER rescue cell lines. JJ generated and processed PRO-Seq and ChIP-seq data. RC analyzed all data and generated figures. RC wrote the paper and CF reviewed and edited the paper. All authors reviewed the paper.

## Data and Materials Availability

All processed sequencing data is deposited in the Gene Expression Omnibus (GSE148789) and raw data is deposited in the SRA archive (PRO-Seq: SRP256595; ChIP-Seq: SRP256596). Supplemental sequence and motif data is available at (https://github.com/rcosby13/Cosby2020_supp_data), and code for processing the PRO-Seq data is available at (https://github.com/JAJ256/PROseq_alignment.sh<u>; commit</u> 55a08db). All cell lines and reagents are available upon request.

## Materials and Methods

### Cell lines and culture methods

The following cell lines were used in this study: *Myotis velifer* embryonic fibroblasts, *Myotis lucifugus* embryonic fibroblasts (both a gift from David Ray, Texas Tech University), *Eptesicus fuscus* immortalized skin fibroblasts (a gift from Woody Wright, UT Southwestern Medical Center; (*55*)), HEK293T cells (a gift from Nels Elde, University of Utah), HEK293T-Rex cells (ThermoFisher; a gift from John Lis, Cornell University), HEK293T-WT cells and HEK293T-KAP1-KO cells (both a gift from Helen Rowe, University College London; (*31*)). All human cell lines were cultured in high glucose DMEM supplemented with 10% FBS, 1% penicillin/streptomycin, and 1% sodium pyruvate. All bat cell lines were cultured in high glucose DMEM supplemented with 20% FBS, 1% penicillin/streptomycin, and 1% sodium pyruvate. All cells were grown at 37°C and 5% CO2 and passaged as needed upon reaching 80% confluency. All cell culture experiments were performed in sterile conditions in a biosafety hood.

### Identifying and characterizing transposase fusion genes

To identify transposase fusion genes, we first extracted all eukaryotic transposase-derived Pfam domains (Table S2). These domains were determined to be transposase derived in previous studies (*56*-*67*), via a combination of sequence similarity and phylogenetic analysis. We then searched all NCBI Refseq tetrapod gene annotations (Table S1; Conserved Domain Architecture Tool [CDART]; (*18, 68*)) for gene models that met the following criteria: 1) contained a transposase domain, 2) had two or more exons (to exclude standalone transposases), and 3) RNA-seq/EST support for all annotated introns. Gene models that met these criteria and two vespertilionid bat genes identified in *de-novo* RNA-seq data (Data S2-S3) were considered to be host transposase fusion genes (HTF). Each transposase fusion gene was further characterized to determine its domain structure (Conserved Domain Search, default parameters; (*69*)), originating gene (NCBI) and transposon (Repbase v20170127; (*70*)), where possible, and timing of gene birth. Due to inconsistencies in annotation and genome quality, transposase-fusion gene age was determined using a combination of homology-based searches (BLASTn; NCBI nr/nt and NCBI Refseq_genomes databases; (*68*)) of closely related species and synteny. Specifically, a transposase-fusion gene was considered to be orthologous if it: 1) had a hit containing both the transposase domain and the host domain in the same transcript (nr/nt) or in the same orientation on the same contig (Refseq_genomes) and 2) was located in a syntenic region of the genome, determined by the identity of flanking genes. We assigned genes a taxonomic span based on the most parsimonious evolutionary relationship between all species possessing the gene. Each fusion was also assigned a corresponding age range in millions of years based on estimates of taxa divergence (Timetree; (*71*)). The insertion timing of the originating transposon for each gene was determined by using the TE sequence and 200bp of flanking genomic DNA as a query against genomes of closely related species (BLASTn; Refseq_genomes; (*68*)), requiring a hit with 100% coverage to be considered conserved. Tissue-specific expression data (Transcripts Per Million, TPM) for human transposase-fusion genes were obtained from the GTEx Portal on 03/01/2020 and log10(TPM) values were plotted using the ggplot2 (*72*) package in R (*73*).

### Selection Analysis

For each HTF conserved in at least two species with a divergence > 50 million years (to allow for enough statistical power), we collected and aligned transposase ORF sequences for each species (Kalign, default parameters; (*74*)). Alignments were manually curated and stop codons were removed. These alignments were then used to determine functional constraint of each transposase using the Phylogenetic Analysis by Maximum Likelihood (PAML) package to estimate *dN/dS* (codeml; CodonFreq = 2, model = 0, Nsites = 0, fix_omega = 0, omega = 0.4) (*75*). Significance of *dN/dS* values were determined via comparison to a model assuming neutral evolution (fix_omega = 1, omega = 1; Likelihood-ratio-test [LRT], *p<0.05*, chi-sq. distribution).

### Transposase consensus sequence generation

To generate consensus sequences for the cognate transposons for nine recently-evolved KRAB-transposase fusions (*KTIGD1*, *KMARD1*, *KTIGD3*, *KRABINER, KMARD4, KHATD2, TIGD-G1, TIGD-G4-Zscan29,* and *MARD-G1*), we first determined the boundaries of the transposon by taking the transposon sequence plus increasing amounts of flanking genomic sequence (+/- 200bp intervals) and querying the appropriate genome (*Pteropus alecto* [ASM32557v1], *Chinchilla lanigera* [ChiLan1.0], *Phascolarctos cinereus* [phaCin_unsw_v4.1], *Myotis lucifugus* [Myoluc2.0], *Pelodiscus sinensis* [PelSin1.0], *Anolis carolinensis* [AnoCar2.0], *Mus musculus* [GRCm38.p6], *Bubalus bubalis* [UOA_WB_1], *Fukomys damaranensis* [DMR_v1.0], or *Oryctolagus cuniculus* [OryCun2.0] respectively). We then extracted the sequence of one full-length copy and used it as a query to collect at least ten additional full-length copies of each transposon. These transposon sequences were then aligned (Kalign; (*74*)) and manually curated to correct CpG sites. We then used the curated alignments to generate a majority rule consensus for each element (Data S1) and annotated each consensus for the presence of transposase ORFs and terminal-inverted-repeat (TIR) sequences.

### Determining HTF gene birth mechanism

To determine the birth mechanism for each HTF, we first partitioned the genes into two classes: genes born via splicing or genes born via duplication (either via ancestral whole genome duplications or segmental duplications), the latter of which occur after the gene is born via splicing. We then asked if those genes born by splicing showed evidence of alternative splicing. To do this, we surveyed each gene model and if the fusion transcript (transposase and host sequence) co-occurred with the original host transcript we considered it to have originated via alternative splicing. For a subset of spliced HTFs for which we could construct consensus sequences of its cognate transposon (see above), we also determined whether the splice site was present in the transposon and, if so, whether the sequence was present in the ancestral consensus sequence.

### Tracing the birth and evolution of KRABINER

To determine which bat species have the *Mlmar1 mariner* insertion at the *ZNF112* locus, two complementary approaches were taken: (i) identifying via BLASTn the presence/absence of the insertion in genomic data, where available (*Myotis lucifugus* [Myoluc2.0], *Myotis brandtii* [ASM41265v1; (*76*)]*, Myotis davidii* [ASM32734v1; (*77*)]*, Eptesicus fuscus* [EptFus1.0]*, and Miniopterus natalensis* [Mnat.v1]; (*78*)), and/or (ii) PCR amplification of *Mlmar1*-KRAB in DNA extracted from a variety of bat species (*Myotis occultus, Myotis velifer, Myotis ridelyi, Myotis sp., Kerivoula papillosa, Eptesicus furinalis,* and *Miniopterus magnater*; gift from Robert Baker, Texas Tech University). For the first method (BLAST), bat genomes were queried with the *mariner* insertion +/- 200bp flanking nucleotides, and the insertion was considered to be present at the *ZNF112* locus if the hit covered 100% of the query sequence, including flanking DNA. For the PCR approach, primers were designed to target the unique genomic DNA flanking the *mariner* insertion (Fig. S2A, Table S6; NEB Q5-HF polymerase #M0492). Amplicons were run on a 1% agarose gel to determine presence/absence of the insertion, and amplicons were excised, gel extracted (Zymo Gel DNA Extraction Kit #D4007), and subcloned (ThermoFisher Zero Blunt TOPO PCR cloning kit #K280020). The subcloned inserts were then Sanger sequenced to validate the presence of the insertion (Data S2).

To test for *KRABINER* expression, RT-PCR was performed. RNA was extracted from three bat cell lines, *M. velifer* embryonic fibroblasts, *M. lucifugus* embryonic fibroblasts, and *E. fuscus* immortalized skin fibroblasts (Qiagen RNEasy Mini Kit #74104) and converted to cDNA (Maxima First Strand cDNA synthesis kit with dsDNAse #K1672). To verify correct splicing of the KRAB domain (exon 4) to the *mariner* transposase (exon 6), primers were designed to span the junctions of exons 4-5 (FWD) and 5-6 (REV) (Fig. S2B, Table S6). Additionally, primers were designed in exons 1 and 6 to amplify the full length *KRABINER* transcript (Fig. S2, Table S6). cDNA was amplified from all cell lines using the specified primers and amplicons were gel-extracted, subcloned, and Sanger-sequenced for validation as described above (Data S2).

### KRABINER mutant sequence design

To generate a KRABINER DNA-binding domain mutant, the transposase region of KRABINER was aligned to the sequence of the closely related *Drosophila Mos1* transposon to identify the DNA binding domain. Residues homologous to those known to be critical for *Mos1* binding to its TIRs, either via gel-shift experiments (*32*) or crystal structure (*33*), were mutated (R150A, Q202A, S206A, R208A). To generate a KRABINER KRAB domain mutant, the KRAB sequence was aligned to the KRAB_Abox Pfam consensus domain (PF01352), and several conserved residues that had previously been shown to be critical for KRAB function were mutated (D12A, V13A, E20A, E21A, L32A, Y33A, R34A) (*34*-*37*).

### Vector construction

To generate expression vectors, the ORFs of all KRAB-transposase fusions except for KRABINER were synthesized as gBlocks (IDT) with 15bp of homology on the 5’/3’ end to facilitate In-Fusion cloning (Clontech, #638920) into either the pcDNA3.1+ (Addgene #V790-20; BamHI/NotI sites) for over-expression or pcDNA4/TO/myc-his-B (Thermofisher #V103020, gift from John Schimenti; NotI/XbaI sites) for inducible expression. The pcDNA3.1+-ORFs are N-terminal FLAG-tagged and the pcDNA4/TO/myc-his-B-ORFs are C-terminal myc tagged. Wild-type KRABINER expression vectors were generated in a similar manner, except that the KRABINER sequence was amplified from cDNA extracted from *Myotis velifer* embryonic fibroblasts (Qiagen RNEasy Mini Kit #74104; Maxima First Strand cDNA synthesis kit with dsDNAse #K1672) using primers unique to the KRABINER transcript (Table S6). Mutant KRABINER sequences (DNA-binding and KRAB mutants) were synthesized as gBlocks (IDT) and cloned into the vectors as described for the other KTFs.

Firefly luciferase reporter vectors were generated using the pGL3pro vector (Promega #E1751; a gift from Todd Macfarlan). For each tested KTF, the TIR sequence of the appropriate reconstructed transposon consensus plus 15bp homology on the 5’/3’ end to the pGL3pro vector (BamHI/NotI sites) was synthesized as complementary oligos, which were then mixed and annealed to generate dsTIR fragments for In-Fusion cloning. A sequence scrambled version of each TIR (Fig. S4) was also synthesized and cloned into the pGL3pro vector to test for sequence specificity.

To generate the KRABINER rescue vectors, we digested the PB-TRE-KRABdCas9 (a gift from Joanna Wysocka; (*79*)) with XhoI and MluI to remove the KRAB-dCas9 insert and replaced it (via In-Fusion cloning) with either wild-type (PB-TRE-WT), DNA binding mutant (PB-TRE-mutDBD), or KRAB mutant forms (PB-TRE-mutKRAB) of KRABINER, all PCR amplified from the expression vectors described above (Table S6).

All vectors were validated by Sanger sequencing.

### Luciferase Assays

Luciferase assays were performed in one of two variants: overexpression of KRAB-transposase fusions in HEK293T cells (pcDNA3.1+ vectors) or doxycycline inducible expression of KRAB-transposase fusions in HEK293T-Rex cells (pcDNA4/TO/myc-His-B vectors). In all cases, cells were first seeded at 500,000 cells per well in a 12 well plate (day 1) and allowed to grow. On day 2, cells were transfected with three plasmids (1. KRAB-transposase fusion expression vector or empty vector, 2. firefly luciferase vector [consensus or scrambled TIR, Fig. S4], and 3. pRL-SV40 [Promega #E2231; a gift from Todd Macfarlan], each at 333 ng/*μ*L for a total 1*μ*g DNA) via Lipofectamine 2000 (Fisher #11668030). 2mL of cell culture media was added on day 3, and cells were either treated (pcDNA4/TO/myc-His-B) or not (pcDNA3.1+) with 1*μ*g/mL doxycycline. On day 4, the cells were lysed and each sample split into 5 wells of a 96-well plate (n=5 technical replicates). Luminescence readings for both firefly and renilla were measured via plate reader (Varioskan LUX; Promega DualGlo #E2920). Firefly and renilla values were first blank-subtracted (untransfected cell lysate) and resulting firefly luminescence was normalized to blank-subtracted renilla luminescence. The values for each replicate were then normalized to the average renilla-normalized firefly luminescence of the empty vector condition for each experiment. Each vector combination included a minimum of three independent experiments, and significant difference in mean luminescence relative to the empty vector control was determined via pairwise Wilcoxon Test with Bonferroni multiple testing corrections (sig. if *adj p<0.05*). Luciferase assays were also repeated in HEK293T cells that were either wild-type or KAP1-KO (*80*) (Fig. S4) to test for dependency of the effect on KAP1. All statistical tests were performed in R (*73*). Data boxplots were produced using ggplot2 (*72*).

### Generating and validating KRABINER KO cells

KRABINER KO cell lines were generated using the CRISPR-Cas9 system as previously described (*81*). In brief, a pair of gRNAs (gRNA 1 (GL430169:87458-87477) - CATTTAGTTTCAGCCTCTCATGG, gRNA 2 (GL430169:89264-89286) - TAATACGTAAGCTGCTGTGTGGG) were designed to the unique genomic DNA sequence (Myoluc2.0) flanking the *mariner* insertion, in order to generate a complete deletion of the *mariner* element (Fig. S6A). gRNAs were also designed to be unique to that location, allowing no genic off targets with up to 3 mismatches (BreakingCas/CRISPOR/CasOFFinder; (*82*-*84*)). These gRNAs were synthesized as oligos (IDT), annealed, and cloned into the PX459 vector (Addgene #62988). We then seeded *M. velifer* embryonic fibroblasts into 6 well plates and transfected with the resulting Cas9-gRNA vector pair (5 *μ*g DNA each, for a total of 10 *μ*g DNA) via electroporation (ThermoFisher Neon Transfection System #MPK10025; 1600V, 20ms, 1 pulse). Cells were allowed to recover for one day, and then were treated with 1.5 *μ*g/mL puromycin for 1 week to overcome low transformation efficiency (< 50%). Cells were then seeded at low density (∼60 cells) in 98-well plate format for clonal expansion. Plates were checked 1 week after seeding to eliminate wells seeded with more than one cell. Following clonal expansion, clones were genotyped using PCR primers outside of the gRNA targeted region (Table S6) to identify clones homozygous for the *mariner* deletion (DNA QuickExtract Lucigen #QE09050). Clones passing initial genotyping were then further expanded, and RNA and DNA were extracted (Qiagen RNEasy Mini Kit #74104/Qiagen DNA Blood and Tissue Kit #69504). Clones were again genotyped to verify absence of the *mariner* insertion (Fig. S6A), and amplicons were gel-extracted and sequence verified as described above to identify deletion alleles (Fig. S6C). RNA was extracted, converted to cDNA, and absence of KRABINER transcription was verified using the protocol described above (Fig. S6B; Table S6). We selected a homozygous clone that met the above criteria for use in this study (hereafter referred to as KO).

### Generating and validating *KRABINER* rescue cells

The KO cell line was transfected with 7 µg of one of the following PiggyBac vectors: PB-TRE-WT, PB-TRE-mutDBD, or PB-TRE-mutKRAB as well as 3 µg of the PiggyBac transposase expression vector (gift from Joanna Wysocka; (*79*)) via electroporation as described above. Cells were allowed to recover for 24 hours and then treated with 1.5 µg/mL puromycin for 1 week to select for construct integration. Cells were then clonally expanded and genotyped as described above for KO cells (Table S6). We tested for genomic integration of the KRABINER transgene by designing primers to the CMV promoter (FWD) and the ubiquitin C promoter (REV), which are not present elsewhere in the genome (Fig. S7B; Table S6). PCR products were gel extracted and sequenced as described above to verify presence of correct transgene. We validated expression of the KRABINER transgene at the RNA level via both RTPCR (for WT; reverse primer anchored in the Myc-His tag; Fig. S7C) and Precision-Run-on-Sequencing (PROSeq) (all rescue variants; Fig. S8A). We also validated protein expression and inducibility of the KRABINER rescue transgenes via western blot (mouse anti-myc antibody ThermoFisher #MA1-21316, 1:1000 dilution in 5% milk/TBST; rabbit anti-β-actin loading control Cell Signaling #8457, diluted 1:1000 in 5% BSA/TBST; anti-rabbit HRP conjugated secondary antibody Cell Signaling #7074S; anti-mouse HRP conjugated secondary antibody Cell Signaling #7076S). Both secondaries were diluted 1:5000 in 5% milk/TBST.

### KRABINER rescue transgene immunofluorescence assays

To assess KRABINER protein localization in bat cells, we performed immunofluorescence assays. In brief, WT clonal lines (P4C11, P4B3, P4D7, and P3E12) as well as a mixed population of mutDBD cells were seeded at a density of 200,000 cells per well of a 4-well chambered coverslip (Ibidi #80426) and treated with 1 *μ*g/mL doxycycline (or not) for 24 hours. Cells were then fixed and permeabilized with 300uL of 100% methanol per well for 3 minutes at −20°C. Following fixation, cells were washed 3X w/ sterile PBS for 10 minutes each, and then blocked with 2% BSA/0.01% saponin in PBS for 1 hour at RT with rocking. Cells were washed again as described above, and then incubated overnight with rocking in a humidity chamber at 4°C with anti-myc primary antibody (ThermoFisher #MA1-21316) diluted 1:1000 in blocking solution. On day 2, cells were washed and treated with anti-mouse secondary antibody (ThermoFisher # A28175; 2 *μ*g/mL in blocking solution) for 2 hours at RT, then washed again. Slides were then mounted and DAPI counterstained (Fluoromont DAPI; Southern Biotech #0100-20) and imaged at 100X with an oil immersion objective (EVOS FL).

### Sample preparation for PRO-seq and ChIP-seq

For PRO-seq, all samples (Table S7), were seeded into 15cm plates and processed as described below. Rescue transgenic lines were also either treated with 1 *μ*g/mL doxycycline (induced, OE) or not (non-induced, rescue) 24 hours prior to PRO-seq. For ChIP-seq, WT and mutDBD transgenic cell lines (Table S7) were seeded into 15cm plates, grown to 80% confluency, and treated with 1 *μ*g/mL doxycycline 24 hours prior to the experiment. Cell viability was assessed via Trypan blue staining (Countess II, ThermoFisher #C10228), with >90% viability required for both PRO-seq and ChIP-seq experiments.

### PRO-seq library preparation

PRO-seq libraries were prepared as previously described (*38, 85*). In brief, cells were grown to approximately 80% confluency and were chilled on ice, washed twice with ice cold PBS, incubated in PBS with 1 mM EDTA for 5 minutes, and harvested by scraping. Cells were permeabilized by incubating on ice for 5 minutes in permeabilization buffer (10 mM Tris-Cl pH 7.5, 10 mM KCl, 250 mM Sucrose, 5 mM MgCl_2_, 1 mM EGTA, 0.5 mM DTT, 0.05% Tween-20, 4 U/mL SUPERase-In [Thermo Fisher], 1X EDTA Free Pierce Protease Inhibitors [Thermo Fisher], and 0.1% NP-40). Permeabilization was verified using Trypan blue staining followed by visual inspection. Cells were washed once with permeabilization buffer and flash frozen in LN_2_ in storage buffer (50 mM Tris-Cl pH 8.0, 40% v/v Glycerol, 5 mM MgCl_2_ 0.1 mM EDTA, and 0.5 mM DTT). Nuclear Run-On was performed for 5 minutes at 37°C by mixing equal volumes of permeabilized cells in storage buffer with 2X Run-On buffer (10 mM Tris-Cl pH 8.0, 5 mM MgCl_2_, 1 mM DTT, 300 mM KCl, 40 µM ATP, 40 µM GTP, 40 µM Biotin-11-CTP [Perkin Elmer], 40 µM Biotin-11-UTP [Perkin Elmer], 2 U/mL SUPERase-In [Thermo Fisher], and 1% Sarkosyl). RNA was isolated by TRIzol [Thermo Fisher] extraction and was fragmented by base hydrolysis in 0.2 N NaOH on ice for 5 min. Following neutralization with pH 6.8 Tris-Cl, RNA was passed through an RNase free Bio-Gel P-30 column [Bio-Rad]. The 3’ and 5’ adapters used were identical to those in (*85*) except 6 random nucleotides were added at the ligation junction as a unique molecular identifier (UMI) to facilitate computational removal of PCR duplicates. The 3’ adapter was ligated to the total RNA pool (containing both nascent and non-nascent RNAs) for 1 h at 25 °C using 30 U T4 RNA ligase 1 [NEB] in 10% PEG8000. Nascent RNA was then captured using Dynabeads MyOne C1 Streptavidin beads [Thermo Fisher] and washed once with high salt wash buffer and once with low salt wash buffer as described in (*85*). RNA was 5’ phosphorylated while attached to the beads using T4 PNK [NEB] and then 5’ uncapped using RppH [NEB] while attached to the beads. RNA was then eluted from the beads using TRIzol, the 5’ adapter was ligated using 10 U T4 RNA Ligase 1 for 1 h at 25 °C. RNA was again isolated using Dynabeads MyOne C1 Streptavidin beads, and reverse transcription was performed on the beads using Maxima H-RT [Thermo Fisher]. cDNA was eluted from the beads by heating, and final libraries were amplified for 11 PCR cycles using Q5 High Fidelity DNA Polymerase [NEB]. Libraries were sequenced on an Illumina NextSeq 500 platform using paired end 37 by 37-bp chemistry. Reads for all samples are accessible at GSE148789.

### PRO-seq alignment and processing

The pipeline used to process PRO-seq data is available at https://github.com/JAJ256/PROseq_alignment.sh. Briefly, UMIs were extracted and adapter sequences were trimmed using fastp (*86*). Ribosomal reads were removed by mapping to one copy of the human rDNA repeat using bowtie2 (--fast-local; (*87*)) and retaining unmapped reads. Reads were then mapped to the Myoluc2.0 assembly, which is the closest organism (< 10 million years diverged from *Myotis velifer*) with a publicly available genome, with the sequence of the wild-type KRABINER transgene spiked in as an additional contig, using bowtie2 --sensitive-local. PCR duplicates were removed using UMI-tools (*88*) with the “directional” method (avg. 0.02% duplication rate), and bigWig score tracks for visualization and downstream analysis were generated using deepTools bamCoverage (*89*), with high accessibility regions (contigs AAPE02059006, AAPE02072785, GL429886, GL429923, GL429927, GL429979, GL430411, GL430644, GL431357) blacklisted. Gene Body regions were defined as transcription start site (TSS) plus 500bp to transcription end site (TES) and TSS minus 500bp TES for + and – strand Myoluc2 ENSEMBL gene annotations respectively (GCA_000147115.1). The PRO-seq data was also used to call transcriptional regulatory elements (TREs) for each sample (dREG; (*39*)) and then merged (bedtools merge; (*90*)) to generate a comprehensive TRE set. Read counts in gene bodies and TREs were calculated by generating single nucleotide resolution bedGraph files using a custom script and bedtools map (*90*). TRE annotations and sample bigWig files are available at GSE148789.

### Differential transcription analysis of genes and TREs

Proseq read counts from each sample were combined into a single counts matrix. Differential expression was performed for gene bodies and TREs separately using DESeq2 (*40*). Samples were separated into condition and treatment by both genes and TREs (Principal Components Analysis, rlog normalized counts; Fig. S8B-C). Three comparisons were performed: KO vs WT, WT/mutDBD/mutKRAB Rescue vs KO, and WT/mutDBD/mutKRAB OE vs Rescue. The OE vs Rescue comparison was performed individually for each rescue, controlling for clonal identity. We considered a gene or TRE to be regulated by KRABINER if it exhibited significant (*adj. p<0.05*) reciprocal changes in the KO vs WT and the WT/mutDBD/mutKRAB R vs KO comparison or if it was differentially transcribed in the OE vs R comparison (*adj. p<0.05*). To identify differentially transcribed genes and TREs unique to the WT rescue lines or shared between the WT rescue lines and either the mutDBD or mutKRAB mutant lines, we used the UpsetR package (*91*). Genic transcription across replicates was highly correlated (average Spearman R = 0.97, *p*<0.001; Fig. S8). Where possible, KRABINER regulated TREs were associated with genes as follows: promoter (TSS +/-500bp), upstream (TSS -150kb), downstream (TES +150kb), intergenic (between two genes), or distal (no gene on contig OR >1Mb from nearest gene). Raw expression counts for genes and TREs as well as DESeq2 outputs for all comparisons are available at GSE148789.

### Gene Ontology Enrichment Analysis of KRABINER regulated genes and TREs

To determine if KRABINER regulated genes are enriched for certain pathways or terms, we first generated a list of KRABINER regulated genes, which included genes that exhibited significant (*adj. p<0.05*) reciprocal changes in the KO vs WT and WT rescue vs KO comparisons (n=121) and genes whose promoters were found to be differentially transcribed in the TRE analysis (manual inspection, n=57) (Table S4). We then performed gene ontology (GO) enrichment analysis using the ClueGO package (*41*) in Cytoscape (*92*) for either all genes (n=158) or only downregulated genes (n=113). We specifically assessed the following human gene ontologies: GO Biological Process, GO Cellular Component, GO Molecular Function, GO Immune System Process, KEGG, REACTOME Pathways, REACTOME Reactions, and WikiPathways (all accessed 2/16/2020) using the GO Term Fusion option. Significant terms (*adj. p*<0.05; 2-sided hypergeometric test, Bonferroni step-down) were plotted as a Cytoscape network.

### ChIP-seq sample preparation and sequencing

For each KRABINER expression construct, 2 × 10^7^ cells were pelleted at 500xg for 5 min at room temperature and washed once with 1 mL of PBS + 1X Pierce Protease Inhibitors [Thermo Fisher]. Cells were then fixed with 1% formaldehyde in PBS + inhibitors for 10 minutes at room temperature, with rocking. The fixation reaction was quenched with 0.125M glycine solution for 5 minutes at room temperature, with rocking. Pellets were washed twice with ice cold PBS + inhibitors and stored at −80C until use. Frozen fixed cell pellets were then resuspended in 2 mL lysis buffer (1 % SDS, 10 mM EDTA, 10 mM Tris-Cl pH 8.0, and 1X Pierce Protease Inhibitors [Thermo Fisher]) and sonicated to average fragment size of ∼500-800 bp using a Diagenode Bioruptor. Sonicated chromatin was diluted in 1X TBST+PI (25 mM Tris-Cl pH 8.0, 0.15 M NaCl, 0.05% Tween-20, and 1X Pierce Protease Inhibitors [Thermo Fisher]). For input libraries, 3.33% of total sonicated chromatin from each was pooled across genotypes of the same KRABINER variant for total input material of ∼10% respective to each individual IP. For each IP, 100 µL Pierce c-Myc magnetic beads (Thermo Fisher) were washed twice in ice cold TBST+PI and then added to sonicated lysate and incubated for 16 h at 4 °C. Beads were washed three times with low salt wash buffer (0.1% SDS, 1% Triton-X100, 2 mM EDTA, 150 mM NaCl, 20 mM Tris-Cl pH 8.0), twice with high salt wash buffer (0.1% SDS, 1% Triton-X100, 2 mM EDTA, 500 mM NaCl, 20 mM Tris-Cl pH 8.0), once with lithium chloride wash buffer (0.1% SDS, 1% Triton-X100, 2 mM EDTA, 500 mM LiCl, 20 mM Tris pH 8.0), and three times with final wash buffer (2 mM EDTA, 10% glycerol, 20 mM Tris-Cl pH 8.0). Beads were the resuspended in TE buffer (10 mM Tris-Cl pH 8.0, 10 mM EDTA). Crosslinks were reversed at 65 °C for 16 h using 0.5% SDS and 0.4 µg/µl Proteinase K (Ambion) in a thermomixer. DNA was extracted twice with phenol:chloroform, once with chloroform, precipitated in ethanol, and quantified using the Qubit dsDNS-HS assay. Library preparation was done using the NEBNext® Ultra™ II DNA Library Prep Kit for Illumina® (New England Biolabs, Ipswich, MA) according to manufacturer instructions with 4 cycles of PCR amplification. Final library products were isolated using SPRI beads and sequenced on the HiSeq 4000 platform in PE150 mode by Novogene Corporation Inc (Sacramento CA). Reads for all samples are available at GSE148789.

### ChIP-seq data processing and analysis

ChIP-seq reads were trimmed for adapter sequences and error-corrected by overlap analysis using fastp (*86*). Reads were aligned to the Myoluc2.0 assembly as for PROSeq above using bowtie2 (*87*) with parameters --sensitive-local -I 0 -X 1500 and filtered for uniquely mapping (MAPQ > 1) concordantly aligned reads using samtools (*93*). Replicates were well correlated genome-wide (average Spearman R = 0.7, 10kb bins, Fig. S11, deeptools plotCorrelation (*89*)), and were thus merged to generate a single library size normalized bigWig coverage track for each transgene (deeptools bamCoverage (*89*)), with high accessibility regions (contigs AAPE02059006, AAPE02072785, GL429886, GL429923, GL429927, GL429979, GL430411, GL430644, GL431357) blacklisted. Merged bigWig coverage tracks for each transgene were then normalized to matched input samples (deeptools bigwigCompare –operation ratio (*89*)). Replicates were kept as separate BAM files for peak calling.

DNA binding peaks for each KRABINER variant (WT, mutDBD, and mutKRAB) were called relative to matched input samples using MACS2 ((*42*); -g 1771694414, otherwise default parameters). Peaks called in all three genotypes, which are likely to be spurious, were removed (bedtools intersect, >50% reciprocal overlap; (*90*)). A similar intersection was performed to identify peaks unique to the WT transgene or shared between the WT transgene and either the mutDBD (WT-mutDBD peaks) or the mutKRAB (WT-mutKRAB peaks) transgene. Deeptools was used to generate heatmaps and metaplots of ChIP-seq signal coverage (library and input normalized bigWig files, see above) over the different WT peak sets (computeMatrix -- reference-point TSS –before/afterRegionStartLength 5000 and plotHeatmap; (*89*)).

To annotate the peaks, we first identified sequence motifs enriched in each peak set (WT unique, WTXmutDBD, and WTXmutKRAB peaks) relative to a background set (all mutDBD peaks for the WT unique or WTXmutKRAB peak sets or all mutKRAB peaks for the WTXmutDBD peaks) (HOMER, findMotifsGenome.pl –len 20; (*44*); Data S4-6). We then determined which TEs were enriched for KRABINER binding, as previously described (*94*). In brief, the significance of transposon overlaps with genomic intervals (ie: KRABINER peak sets) is determined by shuffling TE locations while maintaining distance to TSS. Myoluc2 genomic TE locations were determined using RepeatMasker (v.4.5.0; http://www.repeatmasker.org). with a custom bat TE library. A TE family was considered to be enriched if it overlapped more than expected by chance, with a binomial *p value*<0.05 (1000 shuffles). Enrichment plots were made using the ggplot2 package (*72*). All raw and normalized bigWig files and peak files are available at GSE148789.

### Overlap Enrichment Calculations

To determine if KRABINER binding peaks are enriched in or near differentially transcribed genes/TREs, we developed a pseudo-random shuffling method to calculate empirical *p* values. First, we calculate the distance between the peaks and the nearest TSS and discard those peaks that exceed the 90th percentile of the distribution (*d*) in order to reduce bias introduced by the non-chromosomal Myoluc2 assembly. We then generate a matrix of all ENSEMBL TSS coordinates, discarding those located within *d* bp of the end of a contig, which is necessary to avoid placing a shuffled feature beyond the end of a contig. For each overlap test, we then randomly sample an equal number of ENSEMBL TSS and place a feature (average peak size: 500bp) either upstream or downstream (chosen randomly) of the TSS, maintaining distance to TSS for the original feature. We then determine the overlaps between this random set of features within 10kb of a differentially transcribed TRE (GenomicRanges countOverlaps; (*95*)), and repeat the shuffle and overlap process 10,000 times to generate a distribution of expected overlaps. We then compare this distribution to the observed overlaps for the true peaks with differentially transcribed TREs. The empirical p value is determined by the following equation: *p* = ((#expected overlaps ≥ #observed overlaps) + 1)/10001. *p* values are considered significant at *p <0.05*.

---

**Fig. S1:**
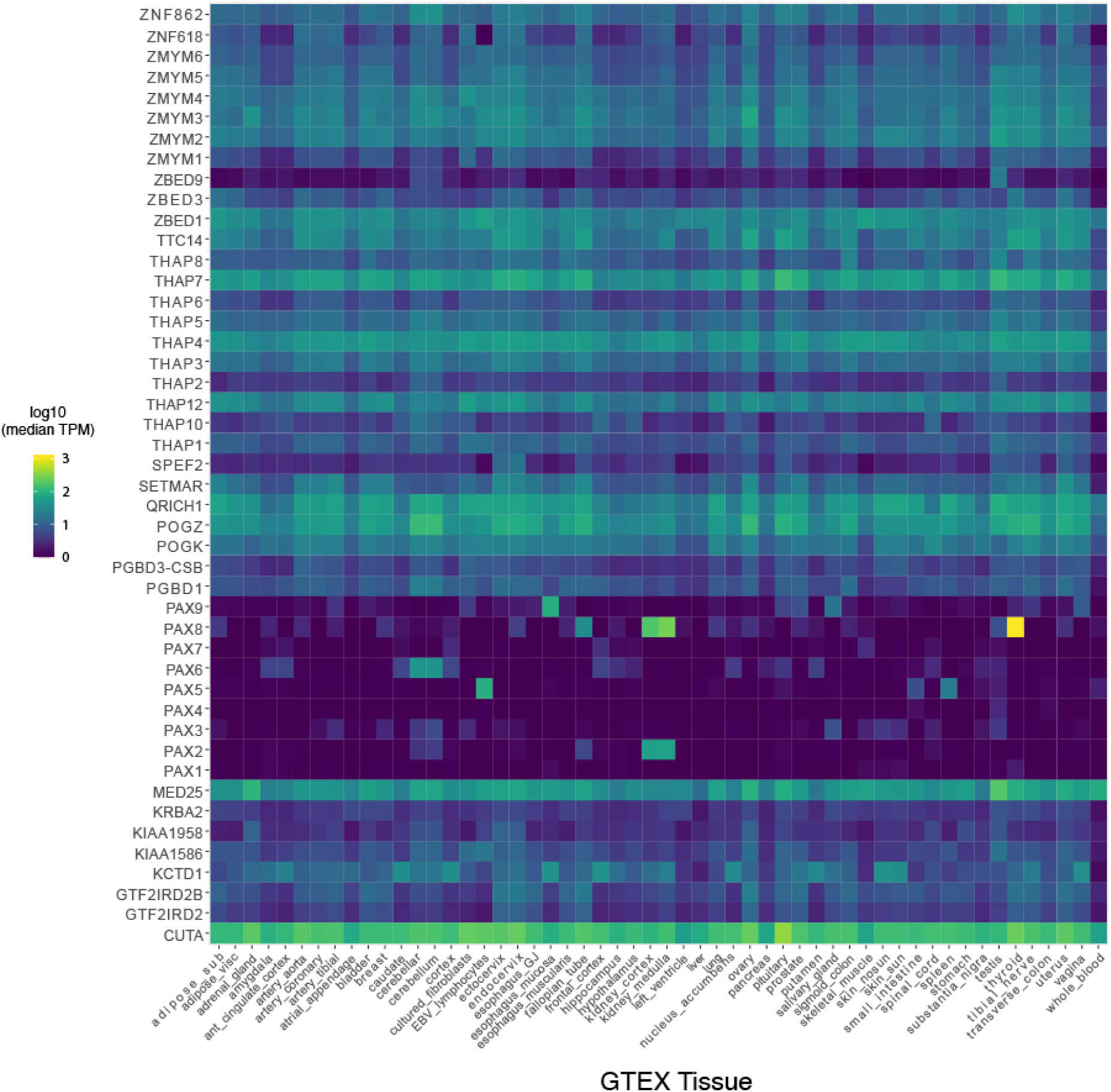
Human HTF genes show variable tissue-specific gene expression and functional constraint. Heatmap plotting tissue specific expression of human HTF genes across 54 human tissues (GTEx, accessed 03/01/2020). Expression is plotted as log10 transformed transcripts per million (TPM).

**Fig. S2:**
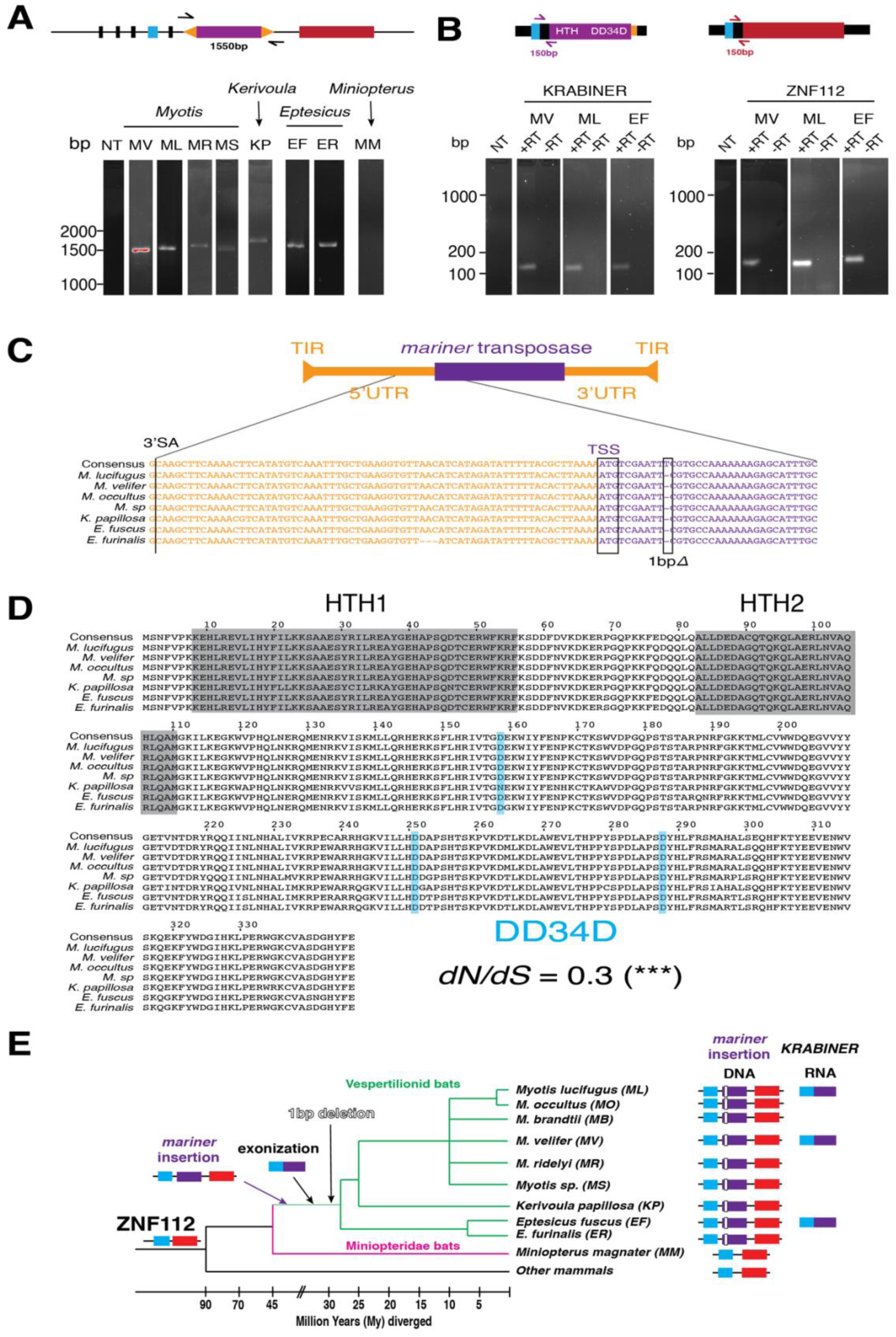
KRABINER evolved in the vespertilionid bat ancestor. A) PCR genotyping of the *Mlmar1 mariner* insertion at the ZNF112 locus in 9 vespertilionid bats and one outgroup (MM). B) RTPCR validation of KRABINER (left) and ZNF112 (right) expression in bat cell lines. C) Partial alignment of the ZNF112 *Mlmar1 mariner* insertion across 7 vespertilionid bat species compared to the consensus sequence, highlighting the unique 1bp deletion at this locus. D) Amino acid alignment of KRABINER’s *mariner* transposase sequence, including DNA binding domains (HTH) and catalytic domain (DD34D), across 7 vespertilionid bat species. E) Summary of the events required for KRABINER evolution.

**Fig. S3:**
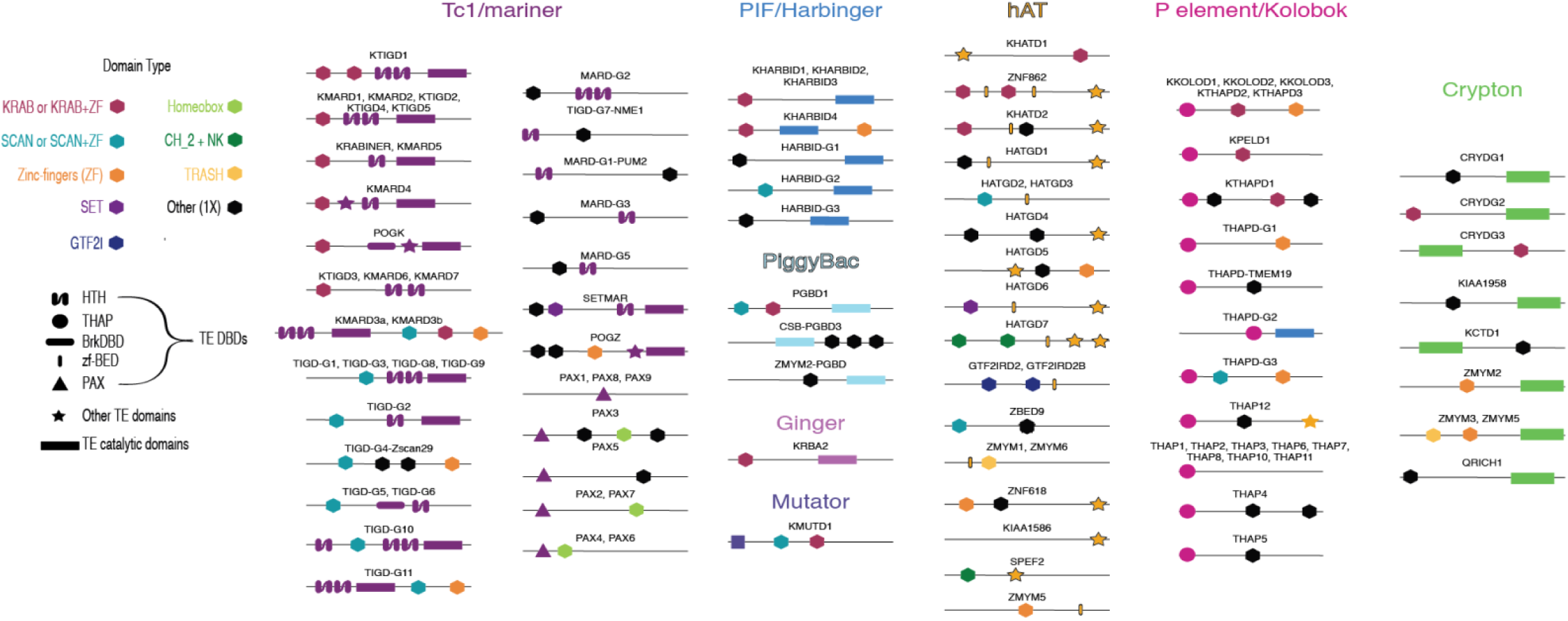
HTF domain structure is varied. Hexagons represent host domains, colored by identity. TE domains are colored by the superfamily that provided them. Some gene architectures are found in multiple fusions.

**Fig. S4:**
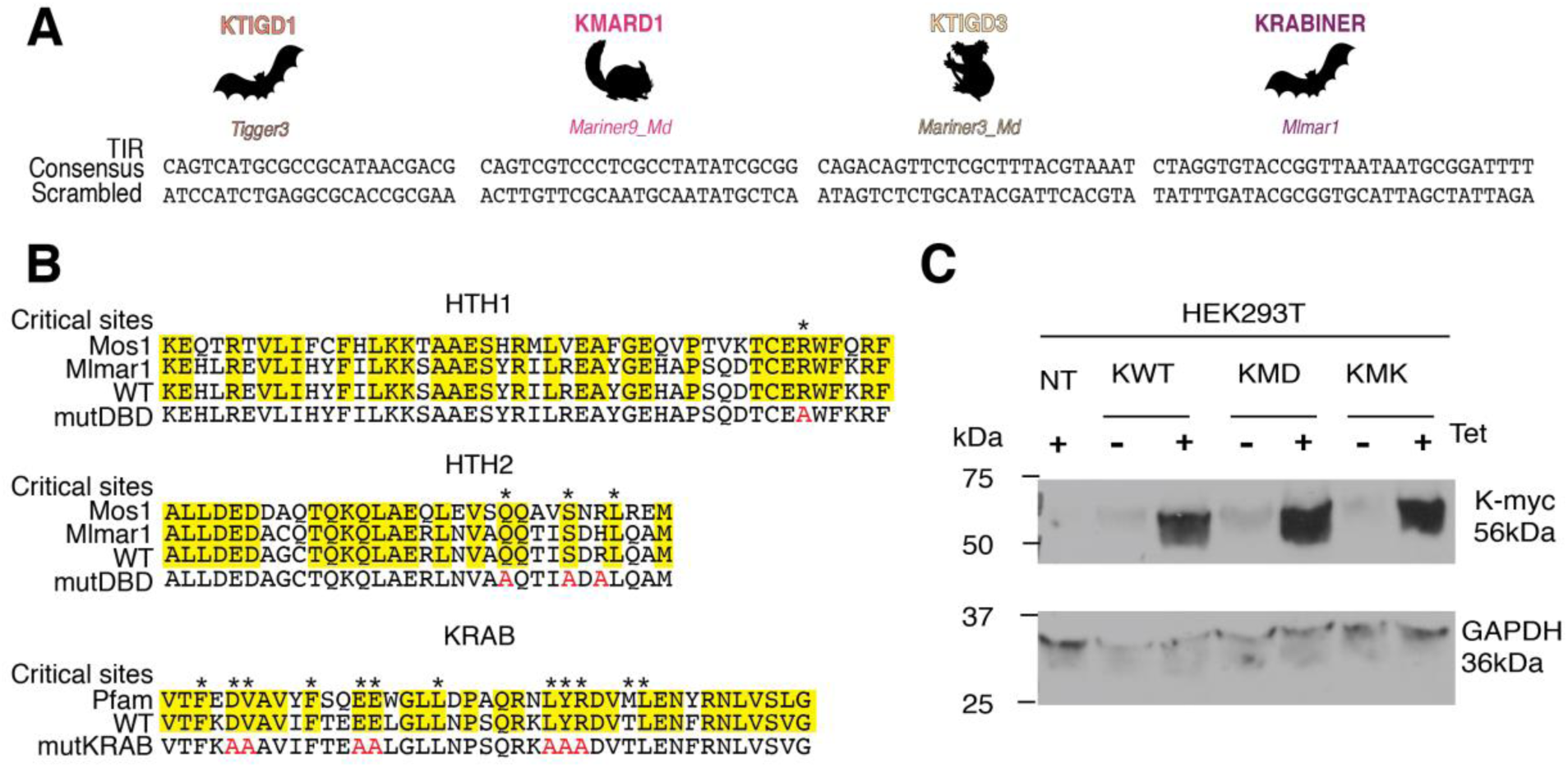
Sequences of TIRs and mutants used in KRAB-transposase luciferase assays. A) Consensus and scrambled TIR sequences used for KRAB-transposase fusion assays. B) Alignments of KRABINER’s DNA binding domains to the closely related *Mos1* transposase DNA binding domains (top) and alignment of KRABINER’s KRAB domain to the Pfam consensus (bottom). Sites shown to be critical domain function are marked with an *; residues mutated in the MUTDBD and MUTKRAB mutant constructs are highlighted in red. HTH=helix-turn-helix C) Western blot showing protein expression of myc-tagged KRABINER or GAPDH in HEK293T cells transiently transfected with the specified KRABINER variant or non-transfected (NT) control. Tet=tetracycline induction.

**Fig. S5:**
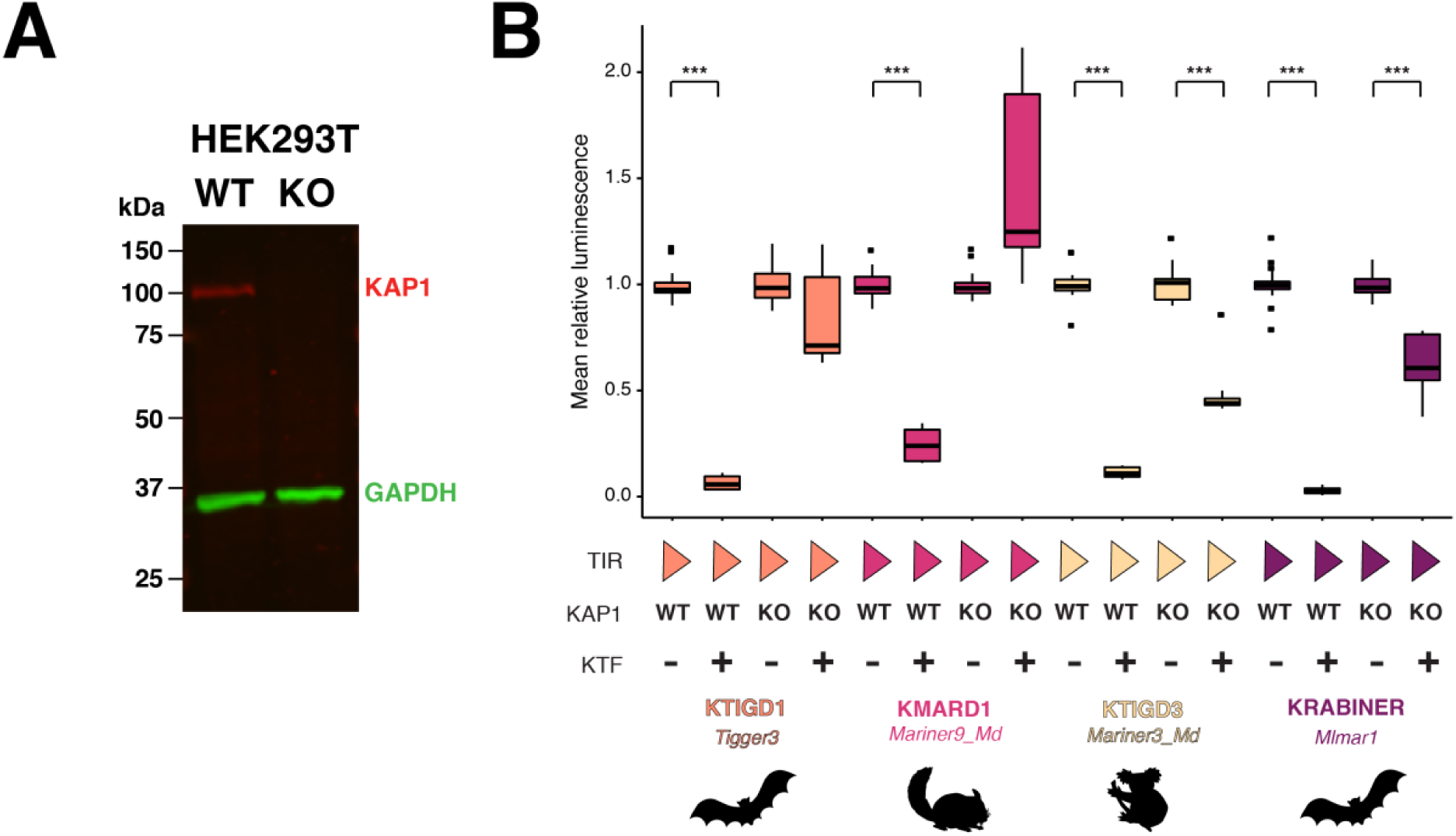
KTIGD3 and KRABINER regulate gene expression in a KAP1-independent manner. A: Western blot validation of KAP1 KO cell lines. B: Boxplot summarizing luciferase assay in HEK293T cells, WT or KAP1 KO, for all four tested KRAB-transposase fusions. TIR=Terminal inverted repeat; Triangle=consensus TIR. *** *adj. p <* 0.001; 2 sample Wilcoxon test, Bonferroni correction. *N=15* per condition.

**Fig. S6:**
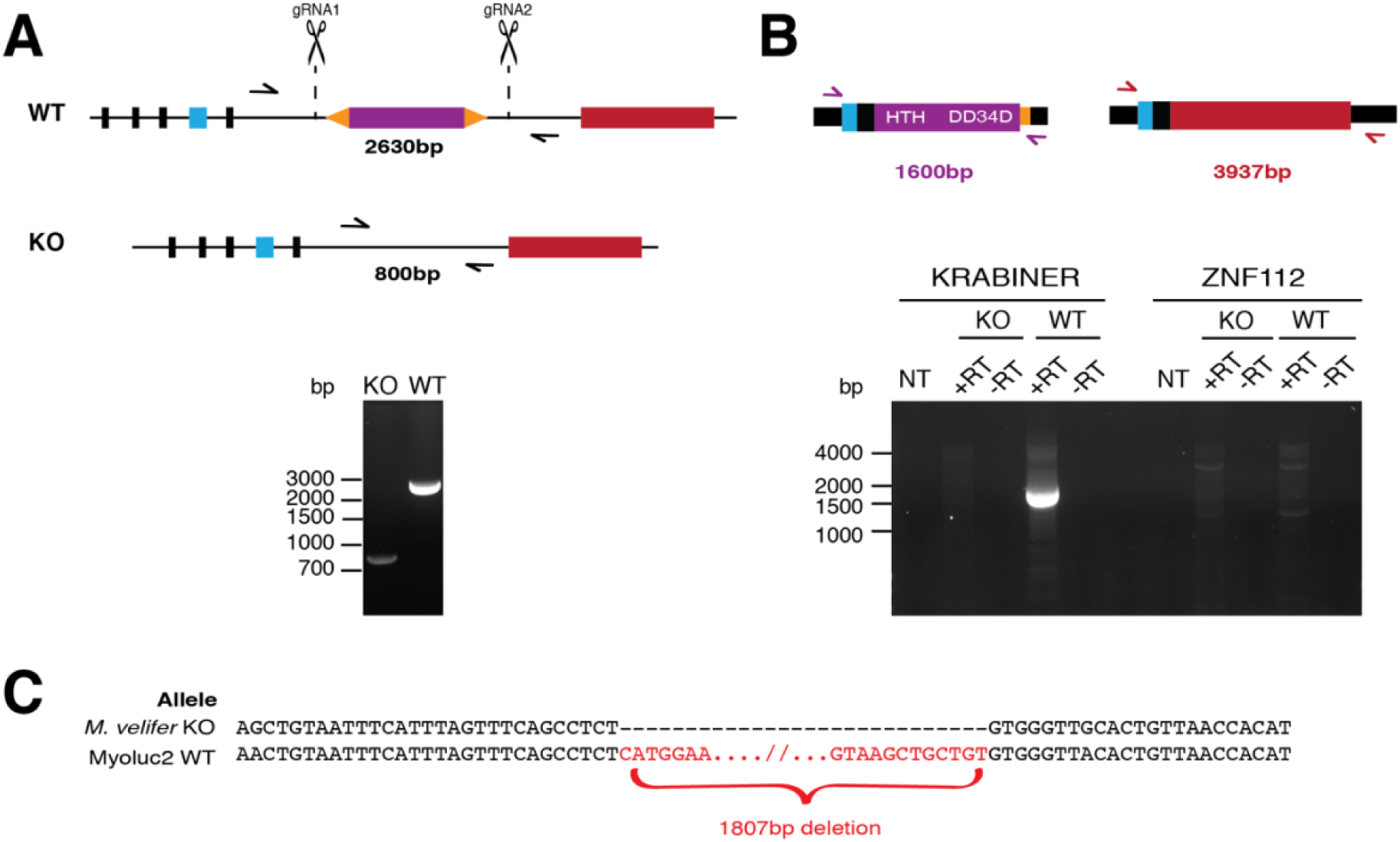
KRABINER KO cell line validation. A) PCR genotyping of KO clone, with estimated product sizes for the WT allele (top) and KO allele (bottom). B) RT-PCR assaying expression of full-length KRABINER and ZNF112 mRNAs in WT and KO cells. C) The KO clone is homozygous for a single 1807bp deletion (top) relative to the Myoluc2.0 reference allele (bottom).

**Fig. S7:**
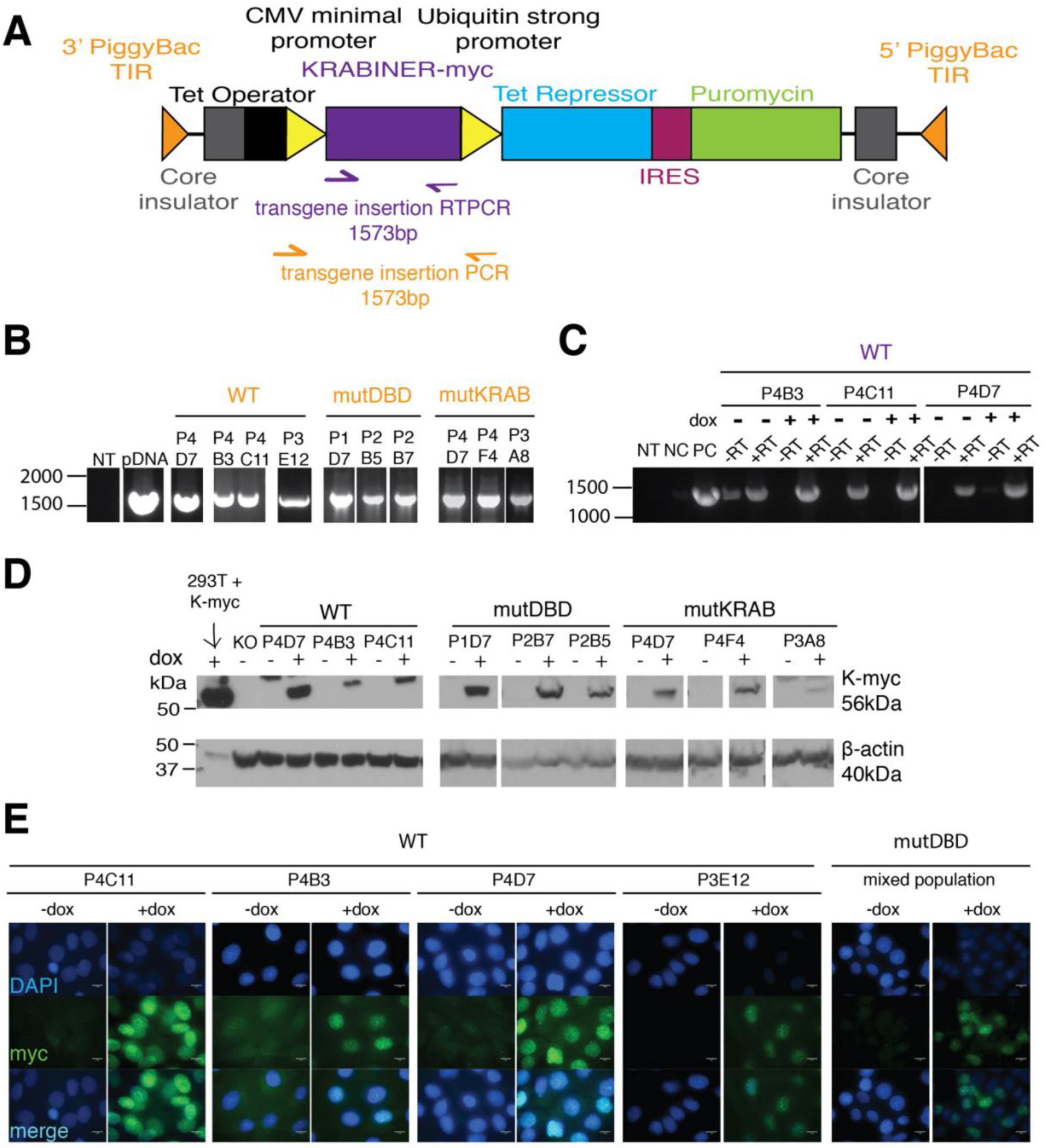
KRABINER rescue cell line validation. A) Schematic diagram of the rescue transgene, with expected product sizes for PCR and RTPCR primers. B) PCR validation of transgene insertion in rescue lines. NT=no template; pDNA=PiggyBac transgene plasmid. C) RTPCR validation of transgene expression in WT rescue cell lines. Negative control (NC)=KO cDNA; positive control (PC)=PiggyBac transgene plasmid. D) Western blot validating transgene protein expression across rescue lines, with beta-actin as a loading control. E) Localization of wild-type KRABINER in clonal rescue lines and DNA-binding mutant KRABINER in mixed populations following doxycycline induction. 100X oil immersion.

**Fig. S8:**
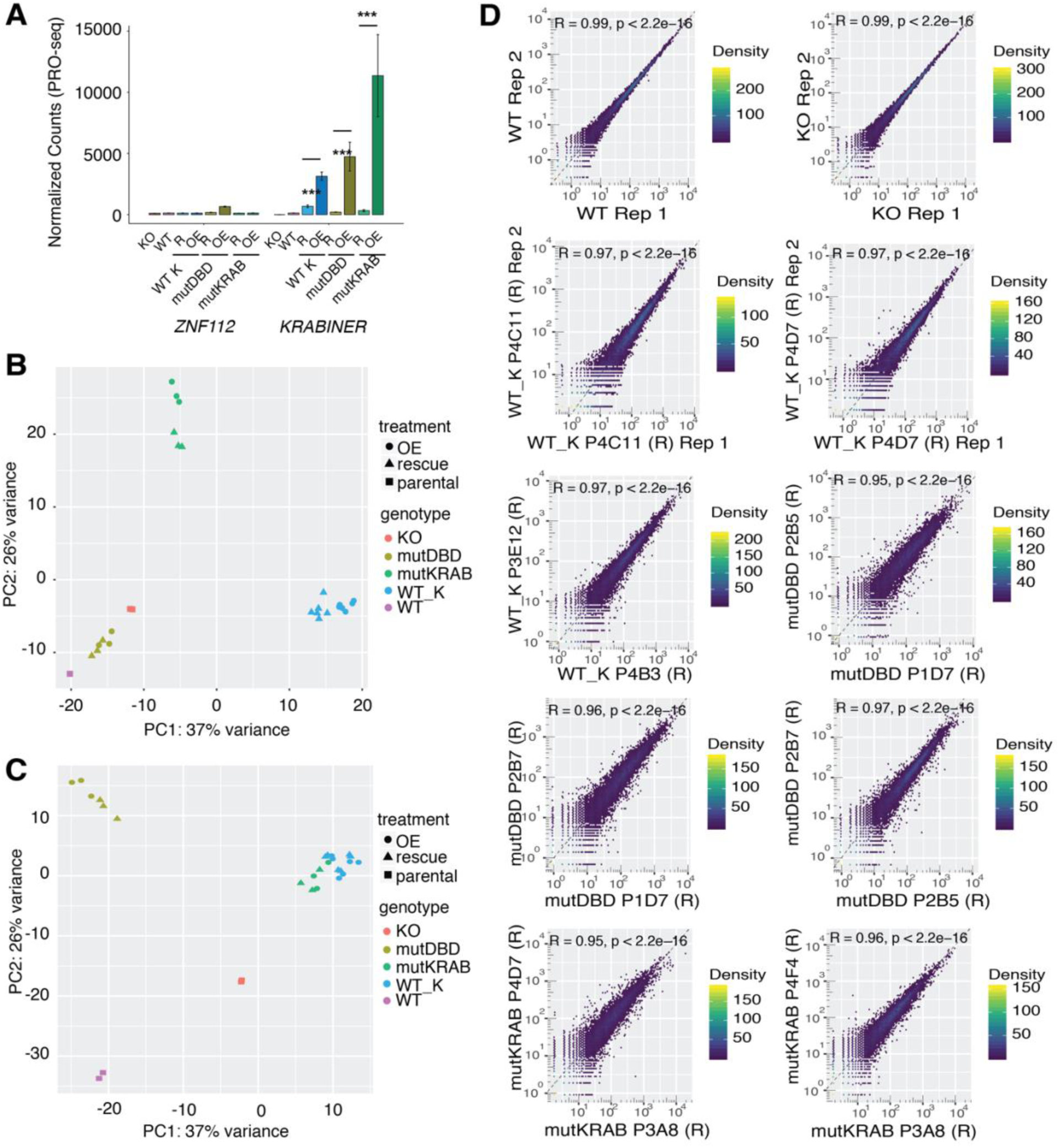
PRO-Seq QC metrics for WT, KO, and rescue cell lines. A) Non-induced KRABINER rescue lines (R) express wild-type levels of KRABINER, while induction results in over-expression (OE). ZNF112 is unchanged across all conditions. DESeq2 library normalized PRO-seq counts; *** *adj. p<0.001* One-way ANOVA with Tukey HSD correction. B-C) PCA plots generated from rlog normalized counts for genic (B) and TRE (C) transcription respectively. D) Representative pairwise replicate correlation scatterplots based on library-size normalized PRO-seq read counts mapped to gene bodies for rescue (R) samples. Density refers to the number of genes within a given hex bin. Spearman *R* is comparable across OE samples.

**Fig. S9:**
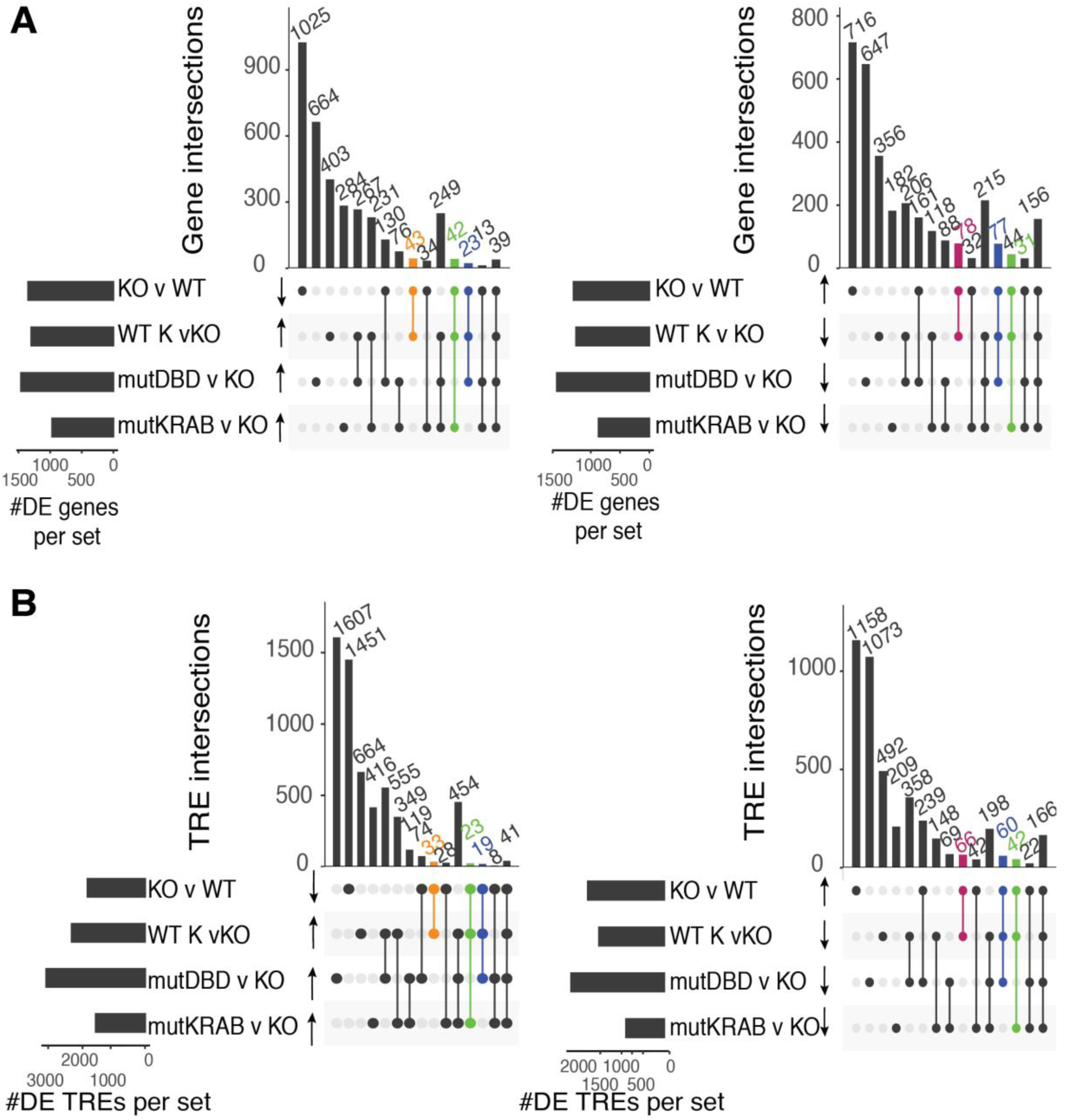
Some KRABINER mediated transcriptional changes require only one of its functional domains. A-B) Upset plots summarizing overlaps between different differentially transcribed gene (A) and TRE (B) sets for the rescue comparison. Left = KRABINER upregulated genes/TREs (orange); right = KRABINER downregulated genes/TRES. Green=genes/TREs rescued by both WT and mutKRAB transgenes; blue = genes/TREs rescued by both WT and mutDBD transgenes.

**Fig. S10:**
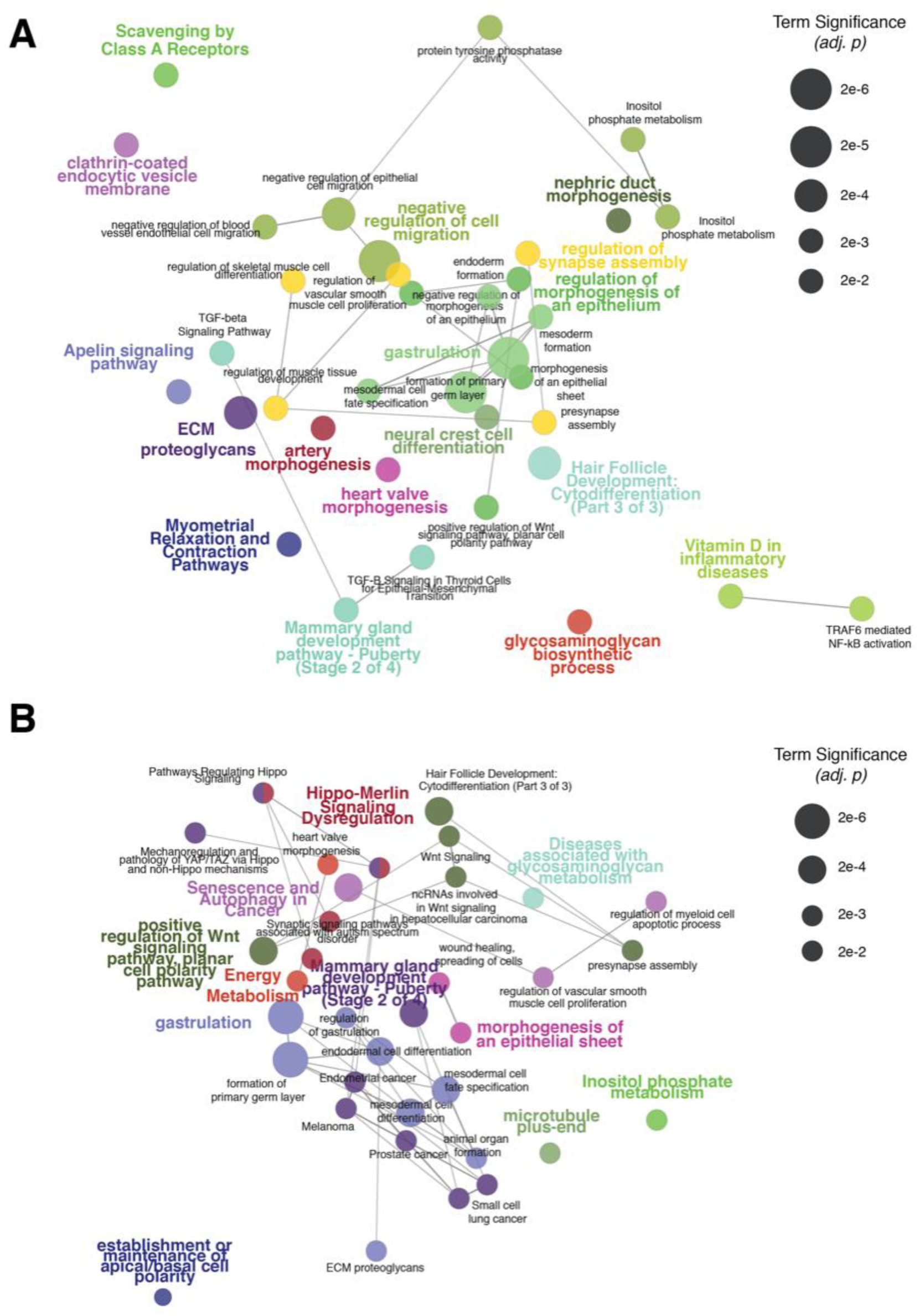
KRABINER regulates a non-random set of genes. Gene ontology enrichment analysis networks for A) all KRABINER regulated genes (n=178) or B) KRABINER downregulated genes only (n=113). Node size for each term corresponds to the significance of each enriched term (*adj. p < 0.05*, Bonferroni step-down). Related terms are clustered into subnetworks, with the most significant term for each subnetwork emphasized.

**Fig. S11:**
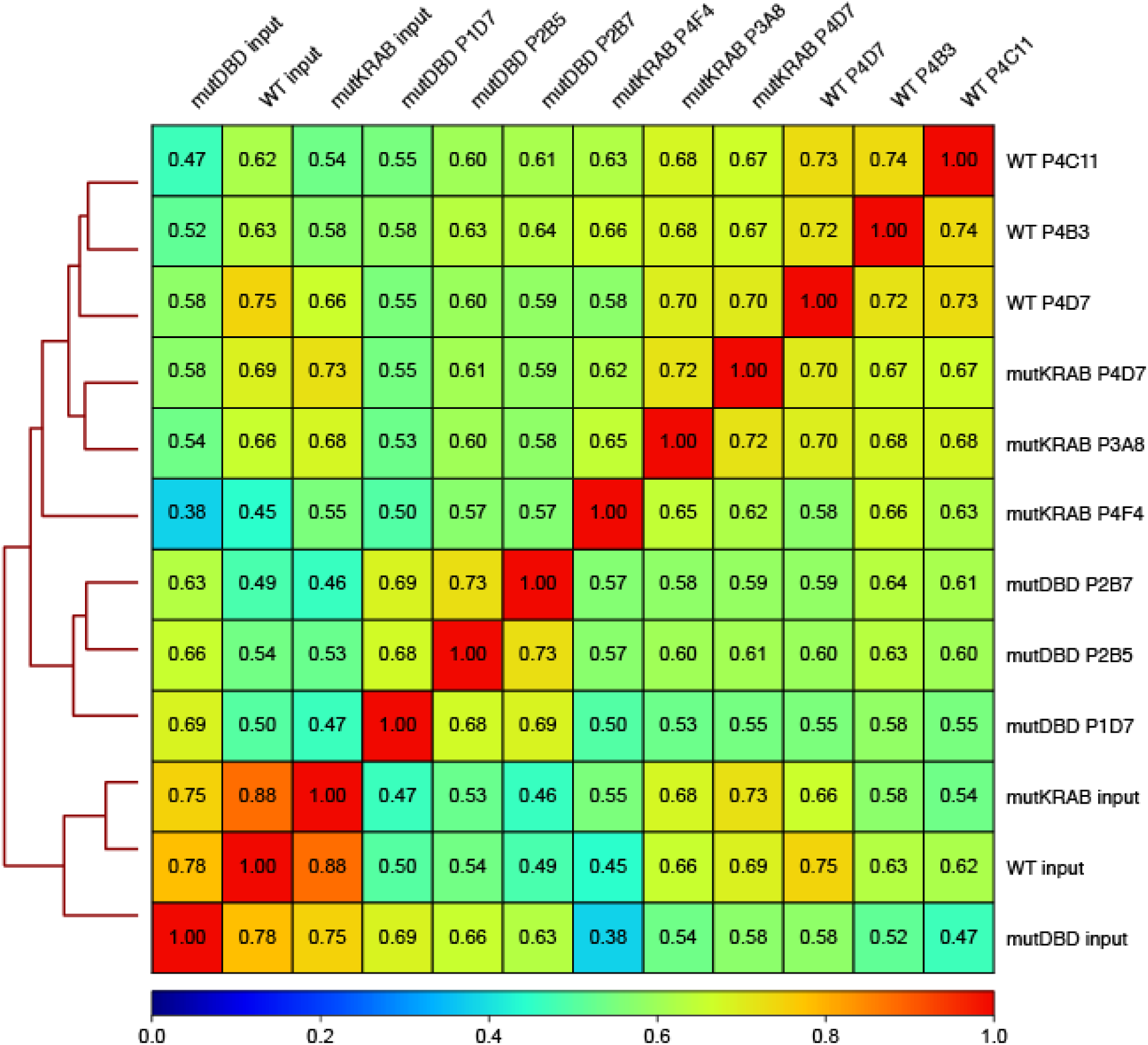
ChIP-seq replicates are correlated. Heatmap summarizing pairwise Spearman correlation between ChIP-seq samples (library size normalized reads, genome-wide 10kb bins).

**Fig. S12:**
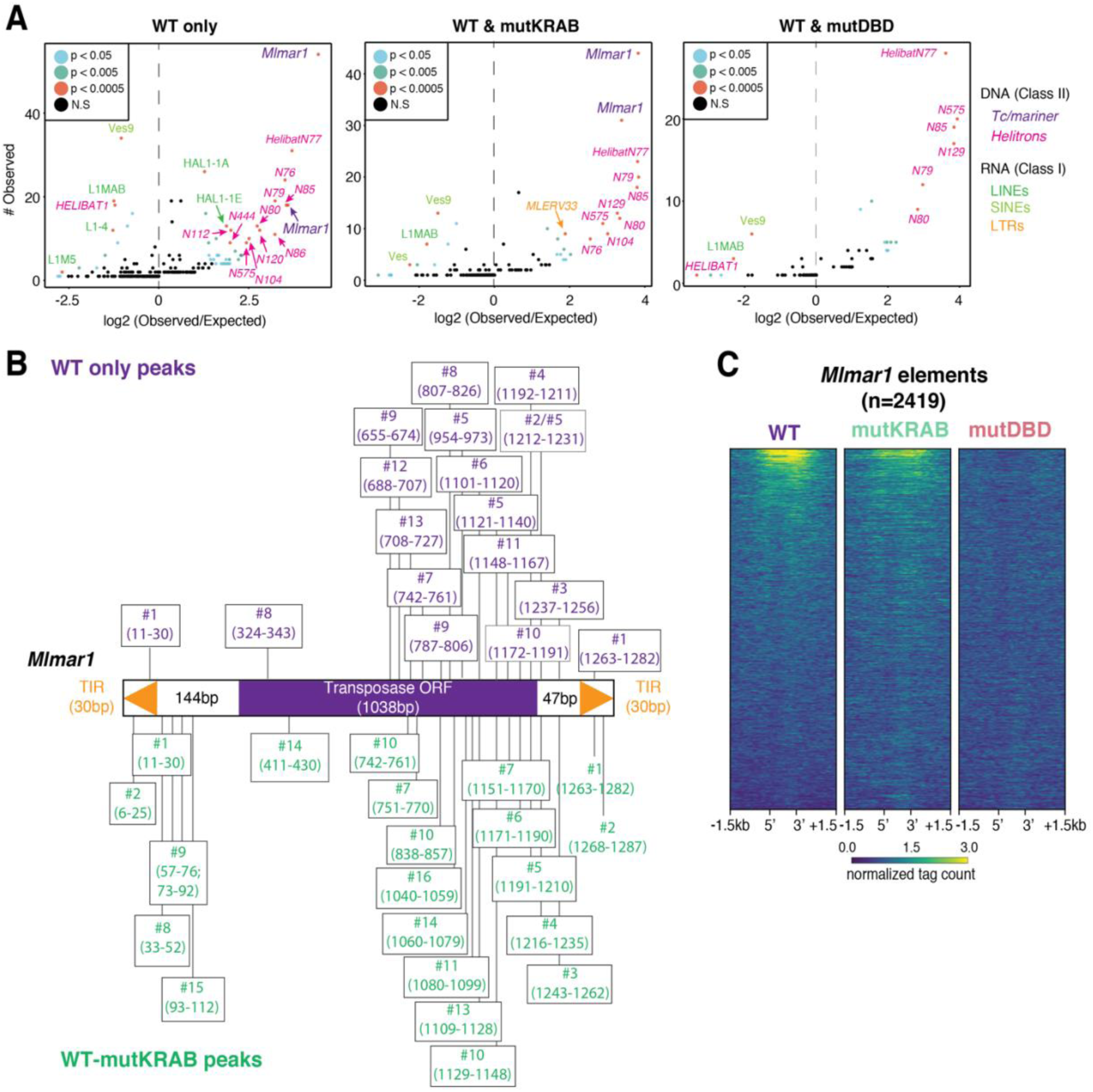
KRABINER binds *Mlmar1 mariner* elements in bat cells. A) Enrichment of transposon families in WT only, WTXmutKRAB, and WTXmutDBD peak sets. Observed = number of overlaps between a TE family and a given peak set. Expected = # of expected overlaps between a TE family and a given peak set after shuffling TE locations 1000 times. *p* values determined using the binomial distribution, n=1000 shuffles. B) Enriched motifs (Data S4-S5) identified in either WT only peaks (purple, top) or WT-mutKRAB peaks (green, bottom) mapped to the *Mlmar1 mariner* consensus sequence. C) Heatmaps summarizing merged, library-size and input-normalized ChIP-seq coverage of each KRABINER variant centered on all *Mlmar1 mariner* elements in the Myoluc2 genome. WT=wild-type, DBD = DNA binding domain, TIR = terminal inverted repeat, ORF = open reading frame, LTR = long terminal repeat, LINE = long interspersed nuclear element, SINE = short interspersed nuclear element.

**Fig. S13:**
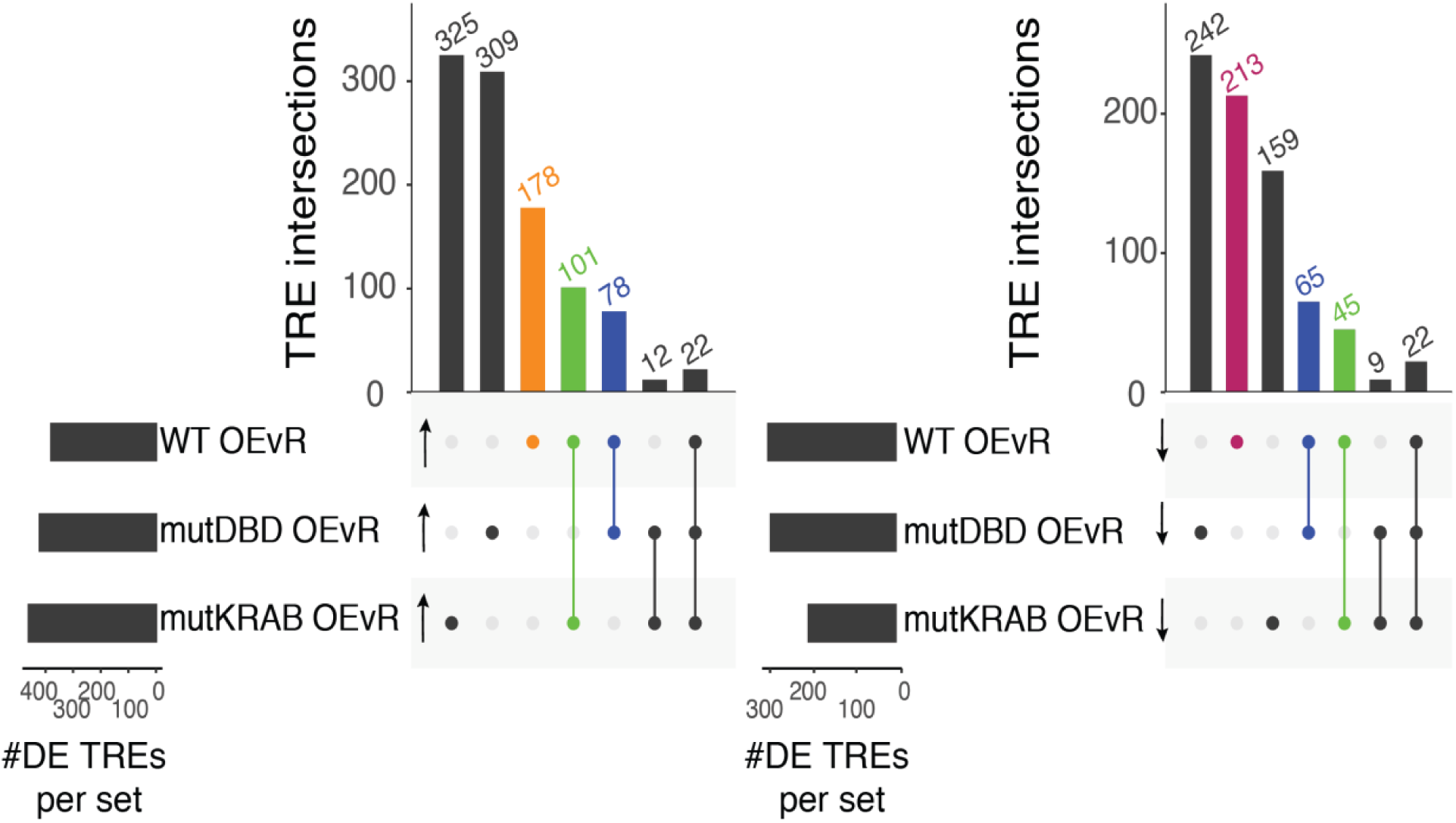
KRABINER over-expression results in changes in TRE transcription in a domain-dependent manner. Upset plots summarizing overlaps between different differentially transcribed TRE sets for the OE vs rescue comparison. Left = KRABINER upregulated TREs (orange); right = KRABINER downregulated TRES. Green=TREs rescued by both WT and mutKRAB transgenes; blue = TREs rescued by both WT and mutDBD transgenes.

**Fig. S14:**
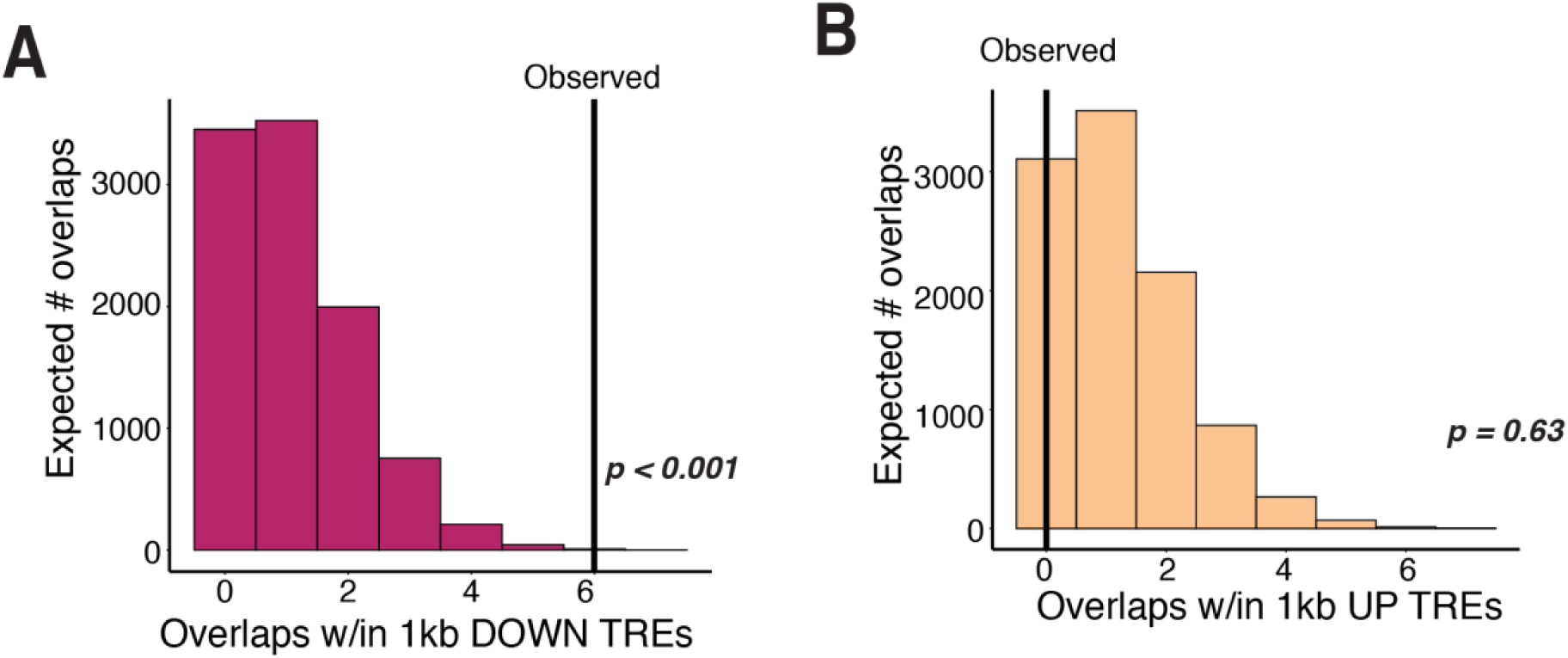
KRABINER binding is associated with transcriptional downregulation of some TREs. Histogram plots summarizing the expected distribution of the number of overlaps of WT KRABINER peaks (all sets, n=1888) within 1kb of either downregulated (A) or upregulated (B) TREs based on shuffling peak locations in the genome while maintaining distance to TSS (10,000 shuffles; see Methods). Lines represent observed number of overlaps for each set; empirical *p* determined via shuffling analysis.

**Fig. S15:**
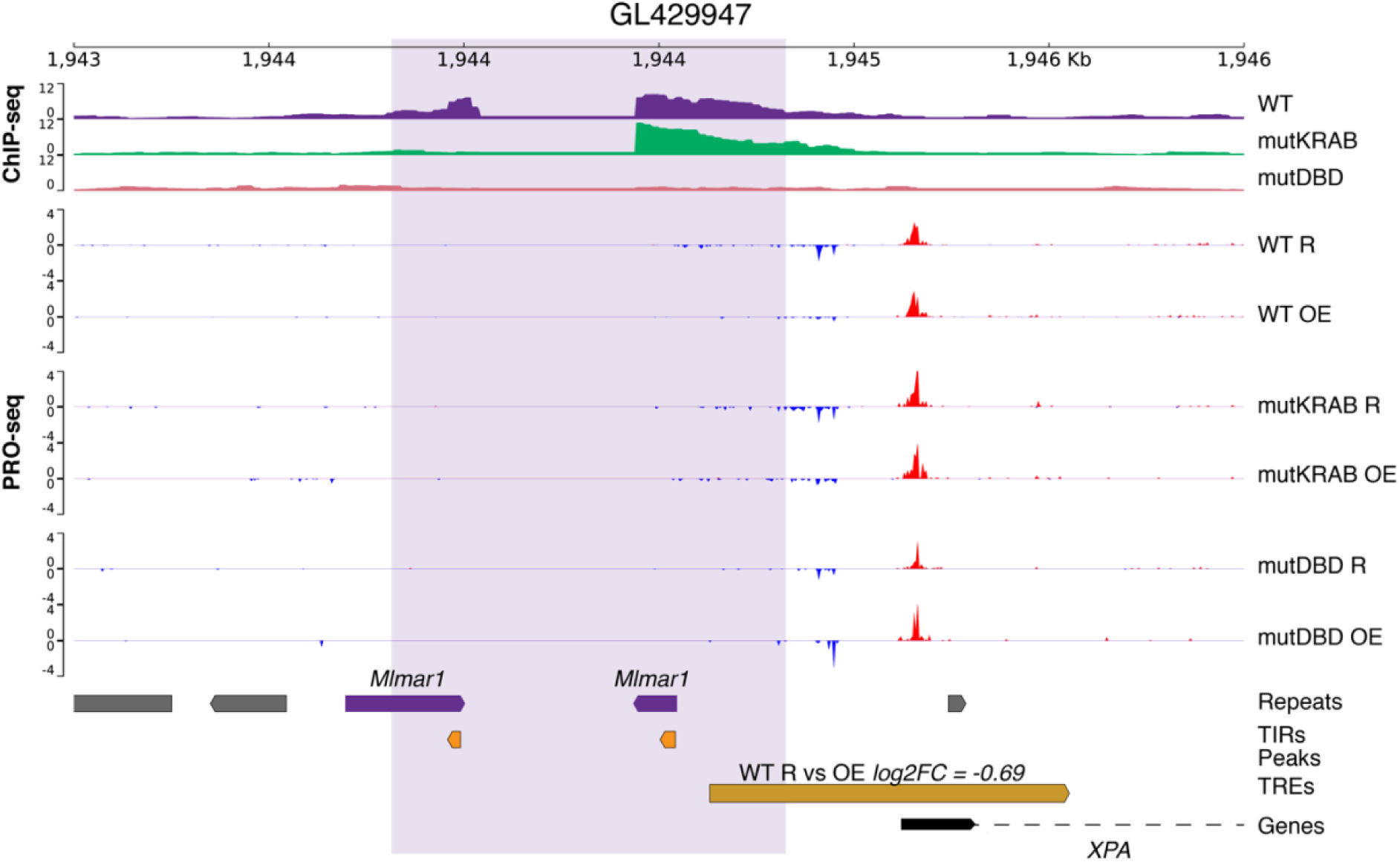
KRABINER regulation of the *XPA* locus. Genome snapshot of the promoter of the *XPA* gene. WT and mutKRAB KRABINER bind to *Mlmar1* TIRs upstream of the promoter, which is downregulated upon overexpression fo WT KRABINER. No KRABINER binding peak is called due to assembly gaps. Pro-Seq signal represents merged replicates (WT *n=*4; mutDBD *n=3;* mutKRAB *n=3*); ChIP-seq signal represents merged replicates (WT *n=3,* mutDBD *n=3,* mutKRAB *n=*3). Log2FC is determined by DESeq2 (OEvsR comparison) and is significant at *adj. p<.05*. Shaded purple box represents the KRABINER-bound region. R = rescue; OE = overexpression; WT= purple; mutDBD=pink; mutKRAB=green.

**Fig S16:**
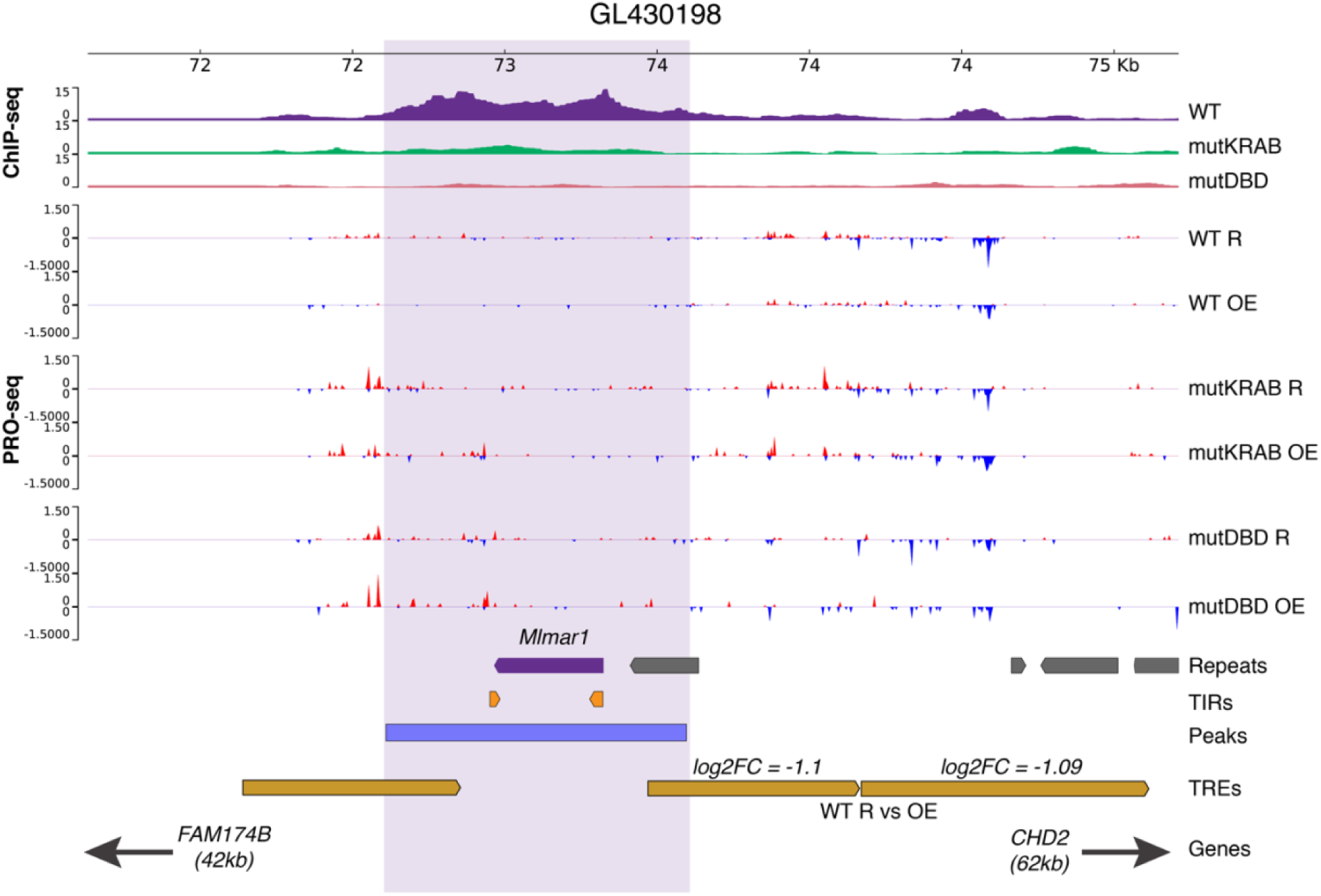
KRABINER regulation of enhancers near *FAM174B* and *CHD2* genes. WT KRABINER binds to two *Mlmar1* TIRs proximal to two enhancers that are downregulated upon overexpression of WT KRABINER. Pro-Seq signal represents merged replicates (WT *n=*4; mutDBD *n=3;* mutKRAB *n=3*); ChIP-seq signal represents merged replicates (WT *n=3,* mutDBD *n=3,* mutKRAB *n=*3). Log2FC is determined by DESeq2 (OEvsR comparison) and is significant at *adj. p<.05*. Shaded purple box represents the KRABINER-bound region. R = rescue; OE = overexpression; WT= purple; mutDBD=pink; mutKRAB=green.

**Fig S17:**
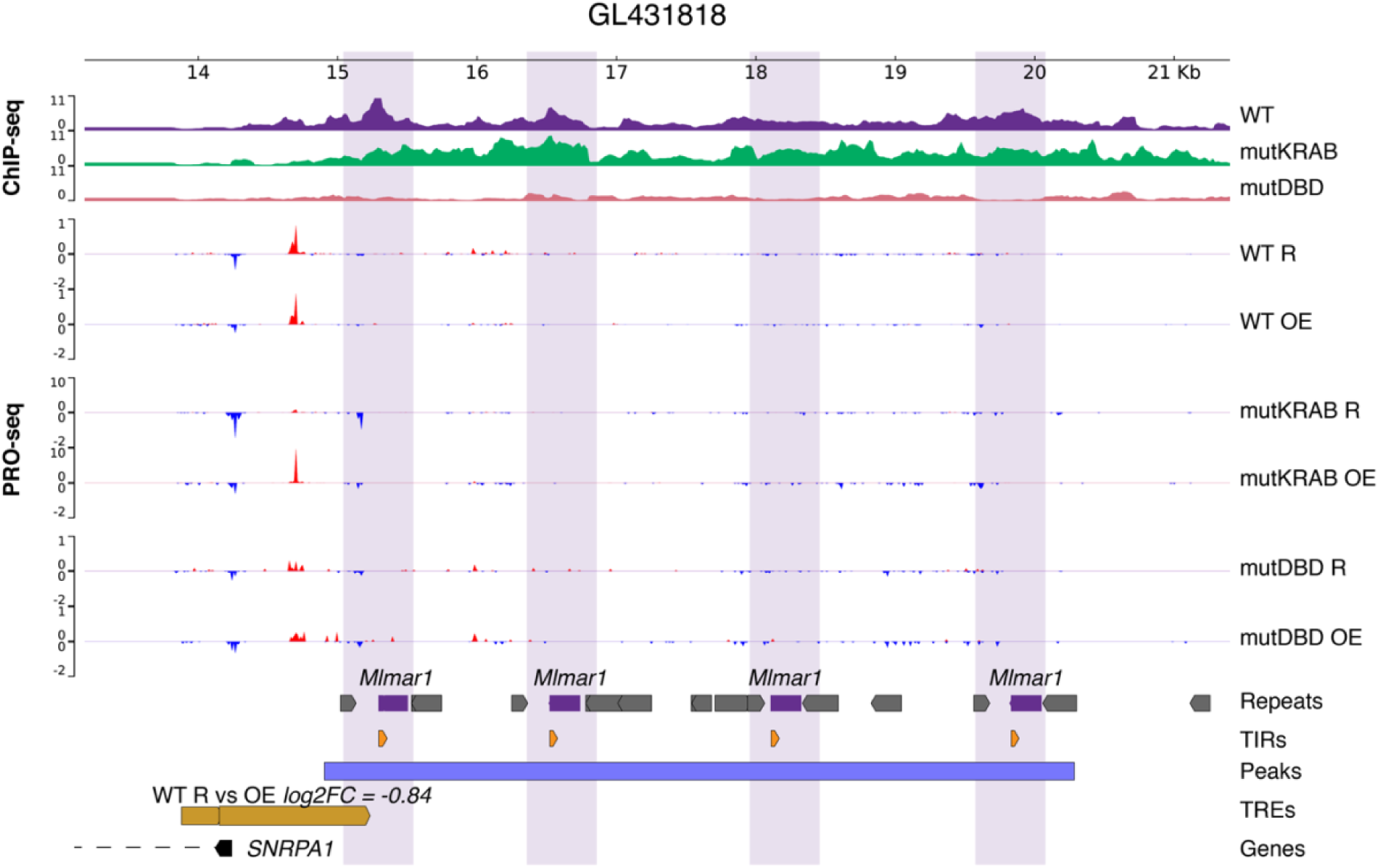
KRABINER regulation of the *SNRPA1* locus. Genome snapshot of the promoter of the *SNRPA1* gene. WT and mutKRAB KRABINER bind to four *Mlmar1* TIRs upstream of the promoter, which is downregulated upon overexpression of WT KRABINER. Pro-Seq signal represents merged replicates (WT *n=*4; mutDBD *n=3;* mutKRAB *n=3*); ChIP-seq signal represents merged replicates (WT *n=3,* mutDBD *n=3,* mutKRAB *n=*3). Log2FC is determined by DESeq2 (OEvsR comparison) and is significant at *adj. p<.05*. Shaded purple box represents the KRABINER-bound region. R = rescue; OE = overexpression; WT= purple; mutDBD=pink; mutKRAB=green.

**Fig S18:**
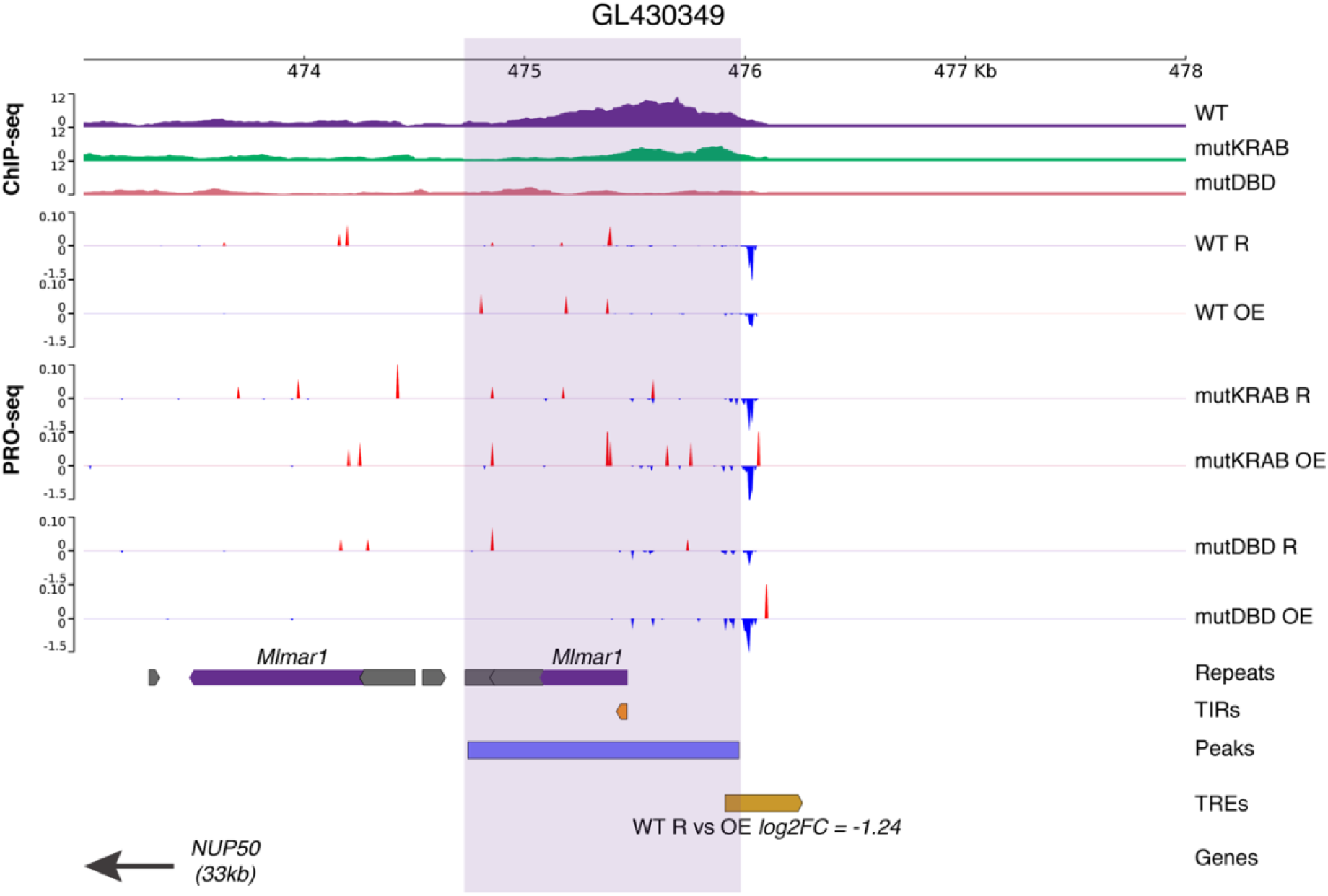
KRABINER regulation of an enhancer upstream of *NUP50* locus. WT KRABINER binds to an *Mlmar1* TIR proximal to an enhancer that is downregulated upon overexpression of WT KRABINER. Pro-Seq signal represents merged replicates (WT *n=*4; mutDBD *n=3;* mutKRAB *n=3*); ChIP-seq signal represents merged replicates (WT *n=3,* mutDBD *n=3,* mutKRAB *n=*3). Log2FC is determined by DESeq2 (OEvsR comparison) and is significant at *adj. p<.05*. Shaded purple box represents the KRABINER-bound region. R = rescue; OE = overexpression; WT= purple; mutDBD=pink; mutKRAB=green.

**Table S1.** NCBI Refseq genomes queried in HTF search

| **Taxa name** | **Taxa ID** | **# genomes** |
| --- | --- | --- |
| Tetrapoda | 32523 | 596 |
| Mammalia | 40674 | 373 |
| Eutheria | 9347 | 366 |
| Metatheria | 9263 | 6 |
| Monotremata | 9255 | 1 |
| Sauropsida | 8457 | 213 |
| Aves | 8782 | 160 |
| Testidunes | 8459 | 20 |
| Crocodylia | 1294634 | 4 |
| Squamata | 8509 | 28 |
| Amphibia | 8292 | 10 |

**Table S2.**
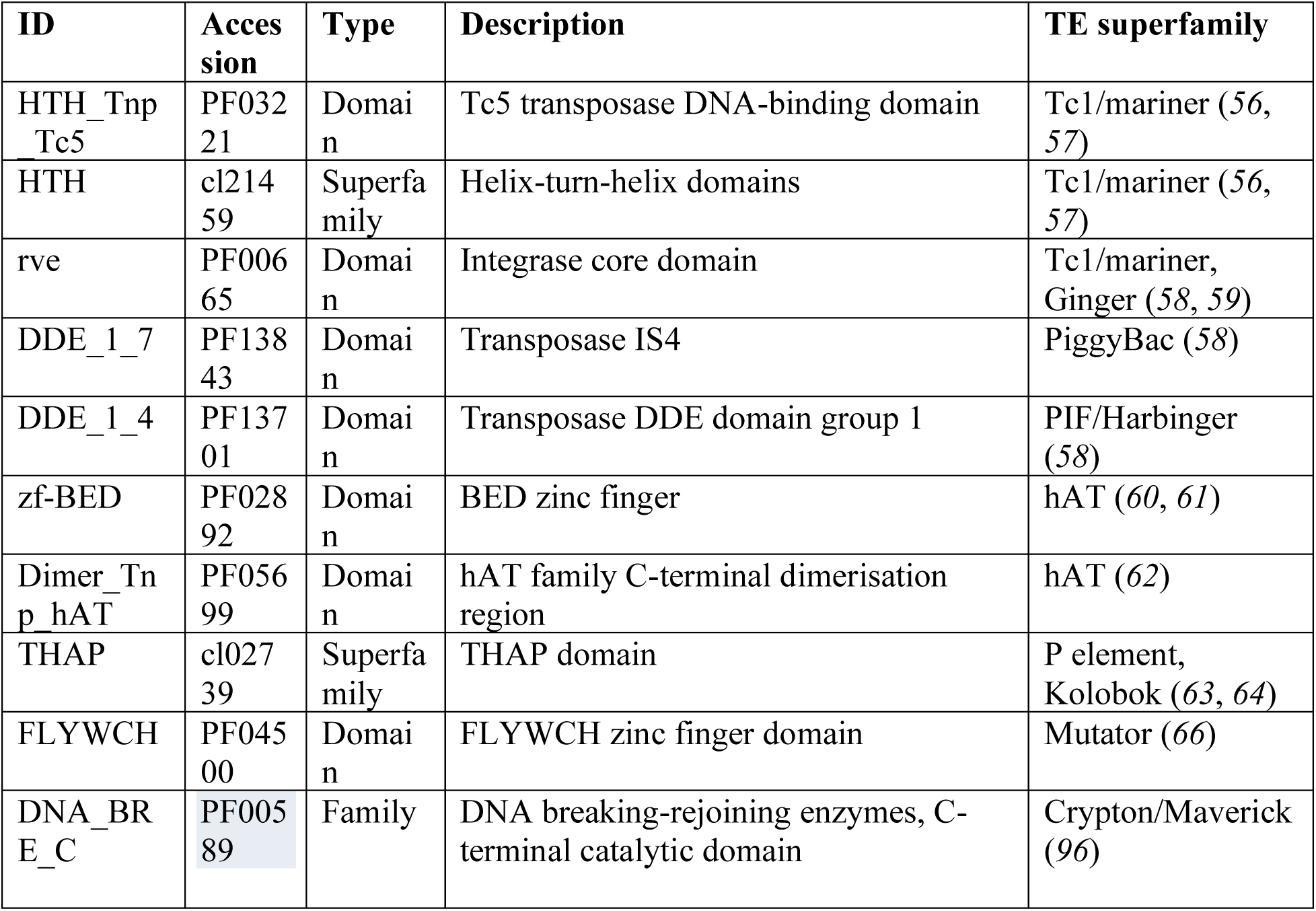
Transposase PFAM domains used as query for HTF search

**Table S3. Summary of identified host-transposase fusion genes**

See attached Excel file

**Table S4: Genes and TREs differentially transcribed by KRABINER**

See attached Excel file

**Table S5: Gene Ontology Enrichment Analysis**

See attached Excel file

**Table S6.**
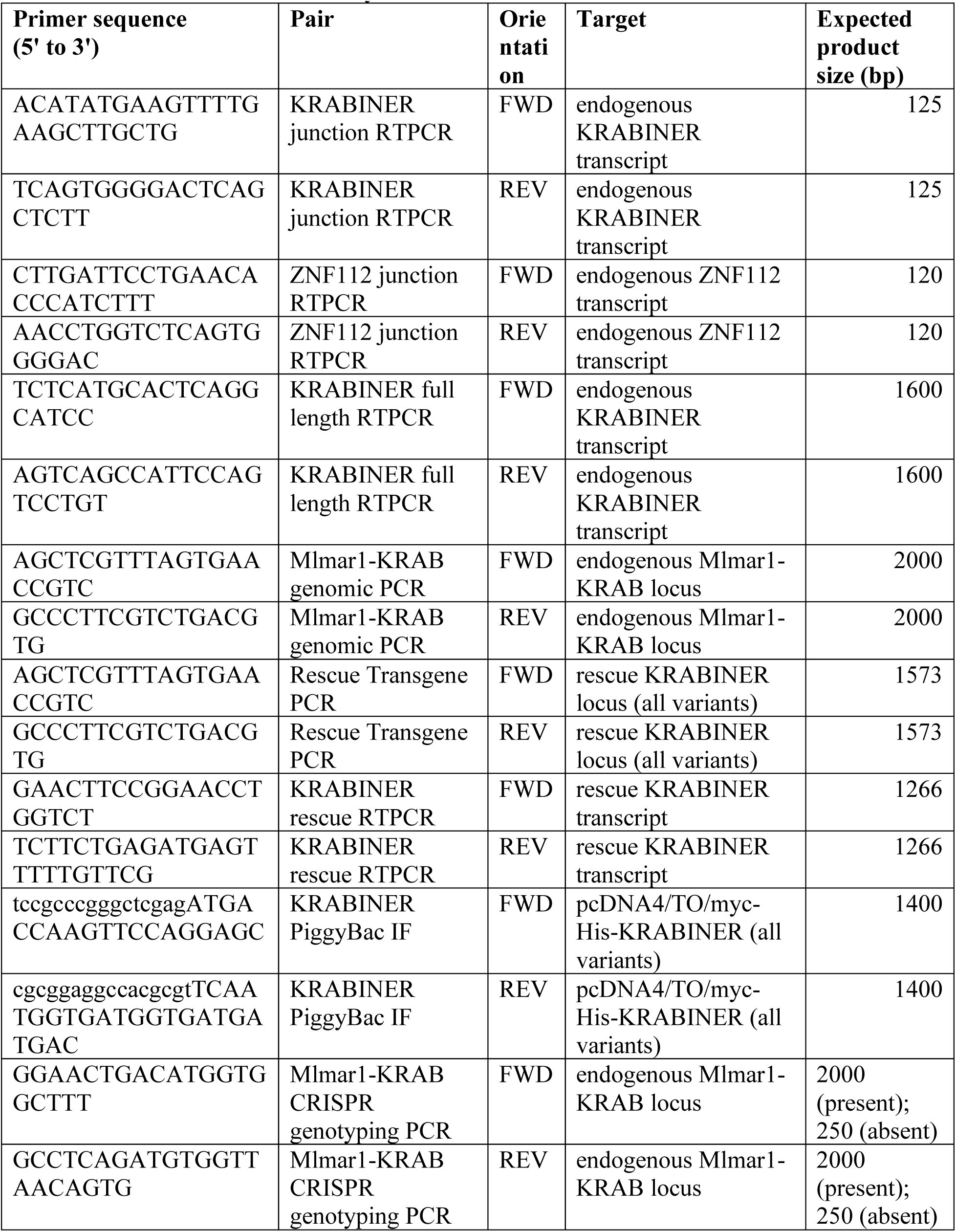
Primers used in this study

**Table S7. Sequencing and alignment statistics**

See attached excel file

**Data S1. Transposon consensus sequences**

Annotated consensus sequences of cognate transposons for nine host-transposase fusion genes.

**Data S2. KRABINER evolutionary history sequences**

DNA sequences of the orthologous *Mlmar1 mariner* insertion responsible for KRABINER in seven vespertilionid bat species, and KRABINER CDS sequences from three bat species.

**Data S3. KTIGD5 coding sequence.**

Sequences of the KTIGD5 CDS for three vespertilionid bat species.

**Data S4. Homer *de-novo* motifs enriched in WT KRABINER only peaks.**

Motif enrichment analysis (HOMER; (*44*)) of WT only KRABINER peaks (n=1070) relative to mutDBD KRABINER peaks (n=4264). Motifs were considered enriched at *p<1e-10*.

**Data S5. Homer *de-novo* motifs enriched in WTxmutKRAB KRABINER peaks.**

Motif enrichment analysis (HOMER; (*44*)) of WTxmutKRAB KRABINER peaks (n=572) relative to mutDBD KRABINER peaks (n=4264). Motifs were considered enriched at *p<1e-10*.

**Data S6. Homer *de-novo* motifs enriched in WTxmutDBD KRABINER peaks.**

Motif enrichment analysis (HOMER; (*44*)) of WTxmutDBD KRABINER peaks (n=291) relative to mutKRAB KRABINER peaks (n=5702). Motifs were considered enriched at *p<1e-1*.

