## Supplemental Data 6 for "Recurrent evolution of vertebrate transcription factors by transposase capture": DataS6_WT-mutDBD_KRABINER_peaks_homerResults.html

./KWT-KMD-only-peaks-KMKbg-20mer/ - Homer de novo Motif Results


### Homer *de novo* Motif Results (./KWT-KMD-only-peaks-KMKbg-20mer/)

Known Motif Enrichment Results  
Gene Ontology Enrichment Results  
If Homer is having trouble matching a motif to a known motif, try copy/pasting the matrix file into
STAMP  
More information on motif finding results: HOMER
| Description of Results
| Tips
  
Total target sequences = 289  
Total background sequences = 7267  
\* - possible false positive  

|  |  |  |  |  |  |  |  |  |
| --- | --- | --- | --- | --- | --- | --- | --- | --- |
| Rank | Motif | P-value | log P-pvalue | % of Targets | % of Background | STD(Bg STD) | Best Match/Details | Motif File |
| 1 | C T G A A G C T T G A C C G T A A C G T G T C A T G A C C G T A C G A T A C G T C G A T T C G A G T C A A T G C A G T C C G A T A G C T C G T A C T G A G T A C | 1e-12 | -2.960e+01 | 3.11% | 0.07% | 32.3bp (35.3bp) | AT3G25990(Trihelix)/colamp-AT3G25990-DAP-Seq(GSE60143)/Homer(0.576) More Information \| Similar Motifs Found | motif file (matrix) |
| 2 \* | C T G A A C G T C T A G T C G A G A C T G A C T T C A G A C T G C A G T T A G C C T G A A G T C G A C T G C T A A C G T A T C G C G T A A T G C C G T A A G T C | 1e-8 | -1.950e+01 | 4.84% | 0.60% | 57.2bp (66.6bp) | HAP4/HAP4\_YPD/1-GZF3,11-HAP4(Harbison)/Yeast(0.591) More Information \| Similar Motifs Found | motif file (matrix) |
| 3 \* | C G T A C A T G A T C G C T G A G A T C C A G T A T G C A G T C A G C T A T G C C G T A T A G C A G T C C A G T A T G C G T A C T A G C A C G T A G C T A G T C | 1e-7 | -1.809e+01 | 3.11% | 0.22% | 46.2bp (58.2bp) | PB0097.1\_Zfp281\_1/Jaspar(0.574) More Information \| Similar Motifs Found | motif file (matrix) |
| 4 \* | A C T G C G T A A G T C C G T A A C T G A C T G A C T G C G T A C G T A A C T G A C T G A G T C C G T A A G T C A G T C A G T C A G T C C G T A C G T A A G T C | 1e-7 | -1.612e+01 | 1.38% | 0.00% | 38.8bp (0.0bp) | PCBP2(KH)/Homo\_sapiens-RNCMPT00044-PBM/HughesRNA(0.558) More Information \| Similar Motifs Found | motif file (matrix) |
| 5 \* | T A C G T C G A A G T C C G A T A C G T G C T A G T A C G C T A G C A T C A G T T G A C G T C A C G A T C A G T C T A G T C G A G C T A C G T A C A T G A C G T | 1e-6 | -1.453e+01 | 2.08% | 0.10% | 28.9bp (27.7bp) | PB0028.1\_Hbp1\_1/Jaspar(0.588) More Information \| Similar Motifs Found | motif file (matrix) |
| 6 \* | A T G C C G T A A G T C G T A C C G T A A G T C A G T C C G T A C T A G A C T G A C T G A C T G C T A G G T A C C T G A A C T G G T A C G C A T G T A C A G T C | 1e-5 | -1.359e+01 | 2.77% | 0.27% | 54.3bp (56.1bp) | BORIS(Zf)/K562-CTCFL-ChIP-Seq(GSE32465)/Homer(0.705) More Information \| Similar Motifs Found | motif file (matrix) |
| 7 \* | A C T G A C G T A C T G A C G T A C T G A C G T A C T G C G T A A G T C C T G A A C T G A C T G C T A G A C T G C T A G A T C G A C T G A C T G A T C G A C T G | 1e-5 | -1.199e+01 | 10.03% | 3.99% | 53.8bp (76.8bp) | SeqBias: polyC-repeat(0.649) More Information \| Similar Motifs Found | motif file (matrix) |
| 8 \* | A C T G A T C G T A G C T G C A G C T A C G T A C G T A C T A G T A C G A T G C G T A C G A C T A C T G C T A G A T C G A T G C T G C A C T A G C A T G C T A G | 1e-5 | -1.161e+01 | 3.11% | 0.46% | 52.5bp (62.4bp) | Zfx/MA0146.2/Jaspar(0.684) More Information \| Similar Motifs Found | motif file (matrix) |
| 9 \* | A C G T A C G T A C T G A C T G A C G T A C G T A C T G A G T C A C G T A C T G A G T C A G T C A G T C A G T C A G T C A C G T A C T G A C G T A G T C C G T A | 1e-4 | -1.138e+01 | 3.46% | 0.60% | 58.8bp (75.5bp) | INSM1/MA0155.1/Jaspar(0.637) More Information \| Similar Motifs Found | motif file (matrix) |
