## Supplemental Data 5 for "Recurrent evolution of vertebrate transcription factors by transposase capture": DataS5_WT-mutKRAB_KRABINER_peaks_homerResults.html

./KWT-KMK-only-peaks-KMDbg-20mer/ - Homer de novo Motif Results


### Homer *de novo* Motif Results (./KWT-KMK-only-peaks-KMDbg-20mer/)

Known Motif Enrichment Results  
Gene Ontology Enrichment Results  
If Homer is having trouble matching a motif to a known motif, try copy/pasting the matrix file into
STAMP  
More information on motif finding results: HOMER
| Description of Results
| Tips
  
Total target sequences = 526  
Total background sequences = 6043  
\* - possible false positive  

|  |  |  |  |  |  |  |  |  |
| --- | --- | --- | --- | --- | --- | --- | --- | --- |
| Rank | Motif | P-value | log P-pvalue | % of Targets | % of Background | STD(Bg STD) | Best Match/Details | Motif File |
| 1 | C G T A C G A T A T G C A G C T C T A G G T A C C G T A A G C T A C G T C T G A A G C T A C G T C T G A G T C A G T A C A G T C T C A G C A T G C G A T C G A T | 1e-80 | -1.847e+02 | 6.84% | 0.00% | 50.0bp (0.0bp) | POU4F3/MA0791.1/Jaspar(0.591) More Information \| Similar Motifs Found | motif file (matrix) |
| 2 | A C T G G T C A C G T A A C G T A G C T G A T C G T C A A C G T C G T A C T G A A G T C G A T C A C G T T A C G G C A T T G C A G T A C C T G A A T G C A T G C | 1e-57 | -1.332e+02 | 5.89% | 0.05% | 63.4bp (38.8bp) | HRB27C(RRM)/Drosophila\_melanogaster-RNCMPT00028-PBM/HughesRNA(0.538) More Information \| Similar Motifs Found | motif file (matrix) |
| 3 | A C G T A C G T A C T G A C G T G T C A C G T A A C G T G T A C T G C A C G T A T G C A A C T G C T G A C G T A C G T A C G T A A G T C C G T A A T G C C T A G | 1e-42 | -9.701e+01 | 3.99% | 0.02% | 54.3bp (27.6bp) | PB0083.1\_Tcf7\_1/Jaspar(0.614) More Information \| Similar Motifs Found | motif file (matrix) |
| 4 | A G T C C G T A C G T A A C T G C G T A A C T G C G T A C G T A C G T A A G T C C G T A A C G T A G T C C G T A C G T A C G T A C G T A A C T G A C G T A C T G | 1e-32 | -7.525e+01 | 3.23% | 0.00% | 57.5bp (0.0bp) | Tcf7/MA0769.1/Jaspar(0.613) More Information \| Similar Motifs Found | motif file (matrix) |
| 5 | A C G T A C G T G T C A C G T A C G T A C G T A T C A G C G T A G T C A C G T A A G C T C T G A A C G T A G T C A G C T C G T A A C G T A G T C C G A T G T C A | 1e-32 | -7.525e+01 | 3.23% | 0.02% | 56.9bp (0.0bp) | RVE1/MA1184.1/Jaspar(0.617) More Information \| Similar Motifs Found | motif file (matrix) |
| 6 | A G C T G C T A A C T G T A G C A C G T G T C A A T G C C G T A A G T C C G T A G A C T A C G T G A C T A G T C A G T C G A T C G T C A G A T C G T C A C A G T | 1e-30 | -6.995e+01 | 3.04% | 0.03% | 52.8bp (18.0bp) | RLR1/RLR1\_YPD/[](Harbison)/Yeast(0.515) More Information \| Similar Motifs Found | motif file (matrix) |
| 7 | G A T C C A G T C A G T A C G T A G T C G C T A C A T G C G A T G A C T C G T A C T G A C G A T G A C T A G C T A T C G A G C T C T A G C A T G C G T A C G A T | 1e-25 | -5.954e+01 | 2.66% | 0.02% | 45.7bp (0.0bp) | Foxh1(Forkhead)/hESC-FOXH1-ChIP-Seq(GSE29422)/Homer(0.568) More Information \| Similar Motifs Found | motif file (matrix) |
| 8 | C A G T C A T G G A C T G C T A G T C A A G T C G C T A G T A C G A T C C G T A A C T G T C A G C G A T G T C A C A G T C A G T A C T G C T G A T C A G C G T A | 1e-23 | -5.444e+01 | 2.47% | 0.02% | 61.8bp (0.0bp) | AGL25(MADS)/colamp-AGL25-DAP-Seq(GSE60143)/Homer(0.608) More Information \| Similar Motifs Found | motif file (matrix) |
| 9 | C G T A A C G T A C G T A C G T A C T G C G T A A C G T C G T A A C G T T A C G A C G T C G T A A C G T A C T G A G T C C G A T C G T A A C G T A C G T A C G T | 1e-21 | -4.992e+01 | 2.66% | 0.04% | 52.0bp (21.1bp) | PB0105.1\_Arid3a\_2/Jaspar(0.568) More Information \| Similar Motifs Found | motif file (matrix) |
| 10 | A T C G A C T G A C G T A G C T A G T C G C T A C G T A C G T A A C G T C G A T A T C G C G T A C G A T T G A C C G T A A C G T A C G T A C G T A T C G A C G T | 1e-21 | -4.992e+01 | 2.66% | 0.05% | 39.9bp (25.7bp) | br-Z1/dmmpmm(Pollard)/fly(0.519) More Information \| Similar Motifs Found | motif file (matrix) |
| 11 | A C G T C T A G C G T A C T A G C T A G T C A G T C G A C T G A T C A G G T C A G T C A A T C G A G C T C A T G T A C G T C G A G T C A G T C A G T C A G C A T | 1e-19 | -4.448e+01 | 2.09% | 0.01% | 41.8bp (0.0bp) | SpiB(ETS)/OCILY3-SPIB-ChIP-Seq(GSE56857)/Homer(0.562) More Information \| Similar Motifs Found | motif file (matrix) |
| 12 | A G T C G C T A A T C G A C T G A G T C A T G C G T A C C G T A T C A G T A C G A T G C T A G C A T G C G A C T A T C G T A G C C T A G A T G C A C T G A T C G | 1e-19 | -4.448e+01 | 2.09% | 0.03% | 40.5bp (68.3bp) | Zfx/MA0146.2/Jaspar(0.606) More Information \| Similar Motifs Found | motif file (matrix) |
| 13 | A C G T C T A G A G C T A G C T A C G T A C G T A C G T C G A T A C T G A T G C G T C A G T C A C G A T C G T A A G T C A G T C C G T A A C G T C T A G A G T C | 1e-17 | -3.962e+01 | 1.90% | 0.01% | 46.7bp (0.0bp) | Pp\_0229(RRM)/Physcomitrella\_patens-RNCMPT00229-PBM/HughesRNA(0.617) More Information \| Similar Motifs Found | motif file (matrix) |
| 14 | C G A T A C G T A G C T A C T G A C T G G T C A T C G A T A G C A C G T T C G A A T G C G T C A A T G C A G C T G C A T T G C A C T A G T C A G T G C A G C T A | 1e-16 | -3.693e+01 | 2.09% | 0.05% | 62.2bp (48.0bp) | ems/dmmpmm(Bergman)/fly(0.502) More Information \| Similar Motifs Found | motif file (matrix) |
| 15 | G A C T C T A G G A C T G C A T A C T G C G A T C G A T G A C T A C G T A C T G G C A T T G A C C G T A T C G A G T A C C G T A C T G A C A T G T A G C G C T A | 1e-14 | -3.277e+01 | 1.90% | 0.05% | 31.4bp (43.8bp) | Tbx20(T-box)/Heart-Tbx20-ChIP-Seq(GSE29636)/Homer(0.583) More Information \| Similar Motifs Found | motif file (matrix) |
| 16 | G T C A C T A G T A C G A C T G G A T C C T G A A C G T A C T G A C G T A C T G A G T C A G T C G T A C A C G T A G C T T A C G C T G A A G C T A G T C C A T G | 1e-14 | -3.277e+01 | 1.90% | 0.05% | 64.7bp (48.1bp) | PB0130.1\_Gm397\_2/Jaspar(0.598) More Information \| Similar Motifs Found | motif file (matrix) |
| 17 | A C G T A T C G C T G A A G C T C A T G A C G T A G T C G T C A A T G C C T G A C A T G C G A T C T A G C T A G T A C G A G T C G T A C C T G A A T C G T A G C | 1e-13 | -3.022e+01 | 1.52% | 0.02% | 46.6bp (0.0bp) | FEA4(bZIP)/Corn-FEA4-ChIP-Seq(GSE61954)/Homer(0.548) More Information \| Similar Motifs Found | motif file (matrix) |
| 18 | G T C A A G C T A C T G A T C G C G A T C G A T A T C G G T A C G C T A C T G A C G T A C G A T T C G A A C G T G A T C C G T A C G T A G T C A C G T A G T C A | 1e-12 | -2.871e+01 | 1.71% | 0.04% | 52.6bp (65.3bp) | exd/dmmpmm(Noyes\_hd)/fly(0.585) More Information \| Similar Motifs Found | motif file (matrix) |
| 19 \* | T G C A T G C A A C G T A G C T A G T C G A T C A G T C G C A T G A T C A G T C A T C G A C T G A C G T C G T A G T C A G T A C C G T A A T G C G T C A C A G T | 1e-9 | -2.129e+01 | 1.14% | 0.02% | 53.3bp (30.9bp) | PB0204.1\_Zfp740\_2/Jaspar(0.511) More Information \| Similar Motifs Found | motif file (matrix) |
| 20 \* | A G T C G T C A A G T C G T A C C A T G C T A G T A C G A G T C G T C A T C G A T G C A G T A C G C T A T A G C T C A G T A G C C T G A C A G T A T C G A C G T | 1e-7 | -1.721e+01 | 1.14% | 0.04% | 46.4bp (38.1bp) | Pax8(Paired,Homeobox)/Thyroid-Pax8-ChIP-Seq(GSE26938)/Homer(0.593) More Information \| Similar Motifs Found | motif file (matrix) |
| 21 \* | C G T A A G T C C G T A A C G T A C G T A C G T A C G T A C G T C G T A C G T A A C T G A G T C C G A T A C G T C G T A C G T A C G T A C G T A C G T A A C G T | 1e-7 | -1.706e+01 | 0.95% | 0.02% | 56.1bp (0.6bp) | hb/dmmpmm(Papatsenko)/fly(0.572) More Information \| Similar Motifs Found | motif file (matrix) |
| 22 \* | A C T G A C T G A C T G T A C G T C A G A T C G A C T G A C T G A C T G A C T G A C T G A C T G A C T G A C T G A C T G A C T G A C T G C A T G A C T G A C T G | 1e-4 | -1.072e+01 | 2.66% | 0.69% | 41.9bp (48.3bp) | PB0097.1\_Zfp281\_1/Jaspar(0.833) More Information \| Similar Motifs Found | motif file (matrix) |
| 23 \* | C G A T A C T G C T A G C G T A C T G A C G A T C T A G C T A G C T G A C G T A C A G T C T A G T C A G C G T A C G T A C G A T C T A G T C A G C T G A C T G A | 1e-4 | -1.030e+01 | 0.76% | 0.04% | 46.3bp (16.2bp) | CEBP:CEBP(bZIP)/MEF-Chop-ChIP-Seq(GSE35681)/Homer(0.572) More Information \| Similar Motifs Found | motif file (matrix) |
| 24 \* | A G T C A G C T A C G T G C A T A C G T A C G T A G C T A G T C A G T C A C G T A G C T A G T C A G T C A C G T A G C T A G T C A G T C A C G T A C G T A G T C | 1e-3 | -9.169e+00 | 0.57% | 0.02% | 28.6bp (12.5bp) | EWSR1-FLI1/MA0149.1/Jaspar(0.790) More Information \| Similar Motifs Found | motif file (matrix) |
| 25 \* | C T A G T C G A A C T G C G T A A C T G C T G A A C T G C G T A A C T G C G T A A C T G C G T A A C T G C G T A C A T G C G T A C A T G C G T A C T A G C G T A | 1e-1 | -2.750e+00 | 0.76% | 0.29% | 42.8bp (49.4bp) | BPC6/MA1402.1/Jaspar(0.965) More Information \| Similar Motifs Found | motif file (matrix) |
