## Supplemental Data 4 for "Recurrent evolution of vertebrate transcription factors by transposase capture": DataS4_WT_only_KRABINER_peaks_homerResults.html

./KWT-only-peaks-KMDbg-20mer/ - Homer de novo Motif Results


### Homer *de novo* Motif Results (./KWT-only-peaks-KMDbg-20mer/)

Known Motif Enrichment Results  
Gene Ontology Enrichment Results  
If Homer is having trouble matching a motif to a known motif, try copy/pasting the matrix file into
STAMP  
More information on motif finding results: HOMER
| Description of Results
| Tips
  
Total target sequences = 1063  
Total background sequences = 6000  
\* - possible false positive  

|  |  |  |  |  |  |  |  |  |
| --- | --- | --- | --- | --- | --- | --- | --- | --- |
| Rank | Motif | P-value | log P-pvalue | % of Targets | % of Background | STD(Bg STD) | Best Match/Details | Motif File |
| 1 | G T C A A G T C A G T C C T G A A C T G C G A T A C G T C G T A C G T A C G A T C G T A C T G A A C G T A C T G G T A C C T A G C T A G C G T A A C G T C G A T | 1e-50 | -1.169e+02 | 2.63% | 0.00% | 47.1bp (0.0bp) | PB0042.1\_Mafk\_1/Jaspar(0.549) More Information \| Similar Motifs Found | motif file (matrix) |
| 2 | C G T A C G T A C G T A C T A G T G A C C G T A A T G C C G A T G A C T C A G T A C G T C A T G T C G A G C T A C T A G G A C T C A G T A C G T A G T C G C A T | 1e-48 | -1.119e+02 | 2.54% | 0.03% | 55.9bp (7.9bp) | AT2G20110/MA1380.1/Jaspar(0.629) More Information \| Similar Motifs Found | motif file (matrix) |
| 3 | C G T A C G T A A G C T A C G T C G T A A G C T A G T C C T G A A C G T C A T G A C G T A G C T A C G T A G C T G A T C C G A T A G C T A C G T A C T G C T G A | 1e-39 | -9.188e+01 | 2.16% | 0.02% | 52.5bp (0.0bp) | SeqBias: polyA-repeat(0.635) More Information \| Similar Motifs Found | motif file (matrix) |
| 4 | T G A C T A C G C G T A C A G T A T C G A C T G C G T A T A G C G T C A A G C T A C G T C G T A A T G C A C G T G A C T C G A T A C T G C T G A T C A G A C T G | 1e-31 | -7.260e+01 | 1.79% | 0.03% | 52.6bp (42.2bp) | FMR1(KH)/Drosophila\_melanogaster-RNCMPT00015-PBM/HughesRNA(0.602) More Information \| Similar Motifs Found | motif file (matrix) |
| 5 | C A T G T C A G T G A C T G C A T G C A C T G A A T C G A T G C T G A C A G C T C A G T T G A C G C T A T A C G T G A C T A G C T G C A T A C G T G C A T A G C | 1e-24 | -5.559e+01 | 1.69% | 0.05% | 59.3bp (51.1bp) | Smad3(MAD)/NPC-Smad3-ChIP-Seq(GSE36673)/Homer(0.542) More Information \| Similar Motifs Found | motif file (matrix) |
| 6 | C T A G C G T A C G T A C T G A C T G A G T A C G T C A G T C A A G C T A C T G T G A C C G A T T A G C A C G T C T A G A G C T C A T G A G C T A C G T G A C T | 1e-23 | -5.416e+01 | 1.41% | 0.02% | 51.8bp (0.0bp) | REF6/MA1415.1/Jaspar(0.574) More Information \| Similar Motifs Found | motif file (matrix) |
| 7 | A T C G C G A T C T A G C T G A T C G A T G C A G A T C T G A C C G T A C G A T A C G T C G T A T C G A G C A T C G T A T A G C A C G T T A C G G T C A A G C T | 1e-22 | -5.205e+01 | 1.79% | 0.05% | 57.0bp (62.0bp) | Barx1(Homeobox)/Stomach-Barx1.3xFlag-ChIP-Seq(GSE69483)/Homer(0.591) More Information \| Similar Motifs Found | motif file (matrix) |
| 8 | C G A T A G T C A C G T G A C T C A G T C T G A A T C G T A C G G A C T A G C T T A G C G C A T T G A C T G C A C T G A C G T A C A G T G A T C G C T A C A T G | 1e-21 | -4.970e+01 | 1.32% | 0.02% | 60.4bp (57.6bp) | HSFB3(HSF)/colamp-HSFB3-DAP-Seq(GSE60143)/Homer(0.599) More Information \| Similar Motifs Found | motif file (matrix) |
| 9 | C G A T A C T G A C T G C T A G C T A G T A C G C T G A T C G A G C T A C G A T T A C G A C G T A C T G C A G T T C G A T A C G G T C A C G A T C T G A T A C G | 1e-19 | -4.532e+01 | 1.22% | 0.03% | 40.5bp (7.4bp) | PF10\_0068(RRM)/Plasmodium\_falciparum-RNCMPT00199-PBM/HughesRNA(0.584) More Information \| Similar Motifs Found | motif file (matrix) |
| 10 | C T A G A C T G C G A T C T G A G T C A G T A C A G T C C G A T A G T C C T G A C T G A C G A T C G A T G C A T G C A T A G C T A G C T A C G T A C T G T C G A | 1e-19 | -4.491e+01 | 1.60% | 0.06% | 50.1bp (29.4bp) | pan/dmmpmm(Pollard)/fly(0.531) More Information \| Similar Motifs Found | motif file (matrix) |
| 11 | T A C G T A C G C G T A C G A T C G T A T C G A A C G T A G T C A G T C C G T A A C T G A C T G C G A T A G T C C T G A C G T A A T G C A G T C C G T A A C T G | 1e-17 | -4.101e+01 | 1.13% | 0.02% | 48.6bp (0.0bp) | WIP5(C2H2)/colamp-WIP5-DAP-Seq(GSE60143)/Homer(0.672) More Information \| Similar Motifs Found | motif file (matrix) |
| 12 | G T A C C G T A C G T A A G T C T C G A A G C T C G A T C T A G T G C A A G T C A G C T C A T G A G T C G T C A C T G A C T A G A T C G G A T C T A G C T C G A | 1e-14 | -3.362e+01 | 1.60% | 0.11% | 41.4bp (39.9bp) | Nr5a2/MA0505.1/Jaspar(0.671) More Information \| Similar Motifs Found | motif file (matrix) |
| 13 | C A T G G C T A C A G T C T A G T A G C C T G A C A G T C A G T C A G T G T C A G T A C C G T A G C A T C G T A C G A T C G T A C G A T C T G A A C G T A G C T | 1e-13 | -3.078e+01 | 1.32% | 0.08% | 47.1bp (46.6bp) | AP1/MA0940.1/Jaspar(0.584) More Information \| Similar Motifs Found | motif file (matrix) |
| 14 \* | C G T A A C T G T C G A C G T A G T C A A C G T A C G T A C T G C G T A G T C A C G T A C G T A G T A C T A G C A C G T C G T A T A C G C T A G C T G A C T A G | 1e-10 | -2.465e+01 | 0.75% | 0.00% | 53.7bp (0.0bp) | At2g41835(C2H2)/col-At2g41835-DAP-Seq(GSE60143)/Homer(0.611) More Information \| Similar Motifs Found | motif file (matrix) |
| 15 \* | C T A G A C T G C T A G A C G T A C T G C G T A A C G T A T C G C T G A A C T G A C T G G T C A T G C A A C T G C T G A A G T C C T G A C A T G A C T G A C G T | 1e-9 | -2.251e+01 | 0.85% | 0.04% | 49.2bp (89.0bp) | ELF3(ETS)/PDAC-ELF3-ChIP-Seq(GSE64557)/Homer(0.577) More Information \| Similar Motifs Found | motif file (matrix) |
| 16 \* | A G T C C G T A A G T C A C T G C G T A C G T A A G T C C G T A C G T A C G T A A C T G A G T C C T G A A C T G A C T G A C T G A G T C A C T G A T C G C G T A | 1e-9 | -2.083e+01 | 0.66% | 0.01% | 46.6bp (0.0bp) | Sox4(HMG)/proB-Sox4-ChIP-Seq(GSE50066)/Homer(0.611) More Information \| Similar Motifs Found | motif file (matrix) |
| 17 \* | A C T G A C T G A C T G C T G A G T C A C A T G T A G C C T G A C T A G A C T G A C T G A T G C C G T A A C T G A C T G A C T G G T C A A T C G A C G T G T A C | 1e-9 | -2.083e+01 | 0.66% | 0.03% | 56.5bp (32.3bp) | EWSR1-FLI1/MA0149.1/Jaspar(0.630) More Information \| Similar Motifs Found | motif file (matrix) |
| 18 \* | G A C T G C T A T C A G T C A G C A G T T C G A C G A T C T A G A C G T T G C A C A G T G T A C G A C T G C T A G C T A G T C A G T C A A C T G G T C A C T G A | 1e-8 | -1.925e+01 | 0.94% | 0.08% | 48.9bp (38.2bp) | KHDRBS1(KH)/Homo\_sapiens-RNCMPT00169-PBM/HughesRNA(0.541) More Information \| Similar Motifs Found | motif file (matrix) |
| 19 \* | A C G T A C G T A C G T A C G T A G T C C G T A A C T G A C T G A G T C A C G T A C T G A G T C A G T C A C G T A C T G A G T C C G T A A G T C A G T C A G T C | 1e-7 | -1.714e+01 | 0.56% | 0.02% | 60.8bp (46.8bp) | RCS1(MacIsaac)/Yeast(0.580) More Information \| Similar Motifs Found | motif file (matrix) |
| 20 \* | G T C A T A G C C G A T A T C G T G C A G T A C G T A C G T A C G A C T T G C A T C G A A G C T C T G A T A C G T G A C G C T A T C A G C T G A T C A G T A G C | 1e-7 | -1.714e+01 | 0.56% | 0.03% | 55.6bp (0.0bp) | Hr46/dmmpmm(Bergman)/fly(0.555) More Information \| Similar Motifs Found | motif file (matrix) |
| 21 \* | G T A C T G A C A G T C G T A C A T G C C T G A G A T C G T C A A G T C A G T C G C A T A C T G A G T C G T A C A G C T A C G T G A T C G A C T T A G C T G A C | 1e-5 | -1.313e+01 | 0.56% | 0.04% | 31.2bp (37.8bp) | ZEB2(Zf)/SNU398-ZEB2-ChIP-Seq(GSE103048)/Homer(0.561) More Information \| Similar Motifs Found | motif file (matrix) |
