## Supplemental Data 1 for "Recurrent evolution of vertebrate transcription factors by transposase capture"

**Data S1 – Transposon consensus sequences**

TIR

ORF

**Splice site Exon|Intron**

>KRABINER_Mlmar1

CTAGGTGTACCGGTTAATAATGCGGATTTTTTTCAATAGATGGAGTTACACATATGTTGATATATATGCGATTTGATATGTATGCTATTTTGTTGTATTGA**CAG|CAA**GCTTCAAAACTTCATATGTCAAATTTGCTGAAGGTGTTAACATCATAGATATTTTTACGCTTAAAAATGTCGAATTTCGTGCCAAAAAAAGAGCATTTGCGGGAAGTTTTAATTCATTACTTTATTTTGAAGAAAAGTGCTGCTGAAAGTTATCGTATACTTCGGGAAGCTTATGGTGAACATGCTCCATCTCAAGATACTTGTGAACGCTGGTTTAAACGCTTTAAAAGTGATGATTTCGATGTGAAAGACAAAGAACGTCCAGGTCAACCGAAAAAGTTTGAAGACCAACAATTACAAGCATTATTGGATGAAGATGCGTGTCAAACTCAAAAACAACTTGCAGAAAGATTAAACGTTGCTCAGCAAACAATTTCCGATCATTTACAAGCAATGGGAAAGATTTTAAAGGAAGGAAAATGGGTGCCACATCAACTGAACGAAAGACAAATGGAAAACCGAAAAGTCATCAGTAAAATGTTGCTTCAACGGCACGAAAGAAAGTCTTTTTTGCATCGAATTGTGACTGGCGATGAAAAGTGGATTTATTTTGAGAATCCCAAATGCACAAAATCATGGGTTGATCCAGGTCAACCATCAACATCGACTGCAAGGCCAAATCGCTTCGGAAAGAAGACAATGCTCTGCGTTTGGTGGGATCAGGAAGGTGTGGTGTATTATGAGCTTCTAAAACCAGGTGAAACCGTTAATACTGATCGCTACCGACAACAAATAATCAATTTGAACCACGCTTTGATCGTGAAACGACCAGAATGTGCCAGAAGACACGGCAAAGTAATTTTGCTTCATGATGACGCACCATCACACACTTCAAAACCAGTTAAAGACACGTTAAAAGATCTTGCCTGGGAAGTATTAACCCACCCGCCGTATTCACCAGACCTTGCTCCTTCAGATTACCACTTGTTCCGATCGATGGCACACGCACTTTCTGAGCAGCACTTCAAAACGTACGAAGAAGTGGAAAATTGGGTCTCTGAATGGTTTGCCTCAAAACAAGAAAAGTTCTATTGGGACGGTATCCACAAATTACCTGAAAGATGGGGGAAATGTGTAGCTAGCGATGGACATTACTTTGAATAAAGCACTTTTGATGTTTCTCTTGAAATTATCGTGTTTTCTTTGATTACAAAATCCGCATTATTAACCGGTACACCTAG

>KTIGD1

CAGTCATGCGCCGCATAACGACGTTTCGGTCAACGACGGACCGCATATACGACGGTGGTCCCATAAGATTATAATGGAGCTGAAAAATTCCTATCGCCTAGTGACGTCGTAGCCGTCGTAACGTCATAGCGCAACGCATTACTCACGTGTTTGTGGTGATGCTGGTGTAAACAAACCTACTGCGCTGCCAGTCGTATAAAAGCATAGCACATACAATTATGTACAGTACGTAATACTTGATAATGATAATAAATGACTATGTTACTGGTTTATGTATTTACTATACTATACTTTTTATTGTTATTTTAGAGTGTACTCTTTCTGCTTATAAAAAAAAATTTTGCTGTAAAACAGTATGCCGTGTTATGCCGGCAGCAGCCTCATACATCTTGTGTTTACCGCGTCTCTTGATTGCATCATTTTCTCTTGTGCTTGATTTAATCTCGTGTTGTTTTGTTCATCA**TG|GCC**CCTAAGCGTACAAAATCCACTGCTAATGTTGCCAGCAAGAGGCCACGTCGAGCGATTGACCTGGAAACGAAATTAAAAGTGATTAAGGACTACGAAGGTGGAAAATCAGTGATGGTTATTGCTCGCCAGTCAGGCACGTCCCATTCCACCATAGCTACGATCTTGAAGAACAAGAACAAAGTGGCAGAAGCTGTTAAAGGATCTGCTTCATTGAAGGCAACGAGACTAACGAAAATTCGAGAAGGGCCTATATCAGGCATGGAGAAACTTCTAATGGCCTGGGCTGAAGACCAGACACAGAAGCGTATCCCTCTCAGCACCGTGGCAATCACGGCCAAAGCAAAAAGTTTGCTTGCGATGTTGAAAGAAAAGGCTGGACCCGACTACGATGTTGAATTTACTGCTAGCTCTGGGTGGTTTAAACGATTCAAGAATCGTTATTCATTACATAATGCGAAAGTGAGTGGTGAGTCTGCGAGTGCTGATGTGAAGGCAGCTGAAGAATTTTTGGAAACTCTAGATAAGCTGATTGTGGAGGAAAATTACTTGCCAGAGCAAATATTCAATATGGATGAAACCTCCCTATTCTGGAAACGGATGCCTGAAAGGACTTTCATCCATAAGGAGGCCAAGTCAATGCCAGGTTTCAAGGCTTTTAAGGACAGGATAACAGTCTTGCTTGGGGGCAATGTTGCAGGCTACAAATTGAAACCCTTTGTGATCCGGCACAGTGAGAACCCCAGGGCCTTCAAGCATATCAATAAGCACACACTGCCAGTGTACTACAGGAGCAATAAGAAGTCATGGATGACCCAGCTCCTCTTCCAAGATGCCCTCCTGAATTGCTATGCCAGCGAAATGGAGAAGTACTGTTTGGAGAATAACATACCTTTCAAGATTTTGCTTATTGTTGATAATGCTCCCGGACATCCTCCTTTTATTGGTGATCTTCATCCCAATATCAAAGTGGTGTTTCTCCCTCCAAACACCACCTCTTTGATCCAACCAATGGATCAAGGAGTTATAGCAGCTTTTAAGGCCTACTGCCTGAGGAGGACCTTTGCCCAGGCTATTGCTGCAACTGAGGAAGACACCGAGAAGACACTGATGCAATTCTGGAAGGATTACAACATCTATGACTGCATCAAGAACCTTGCTTGGGCTTGGGGTGATGTCACCAAGGAGTGTATGAATGGCATCTGGAAGAAGACACTCAAGAGGTTTGTCCATGACTTCAAAGGATTTGCCAAGGATGAGGAGGTTGCAAAAATCAACAAGGCTGTGGTTGAGATGGCAAACAACTTTAACCTGGGTGTGGATGAGGATGACATTGAGGAGCTCCTAGAGGTGGTTCCTGAGGAATTGACTAATGAGGAGTTGTTGGAACTGGAACAGGAACGCATAGCTGAAGAAGAGGCAAGAGAAAAGGAAACTGCAGGAGAAAAAAAAAAAGAACCCCCAAGAAAATTCACAGTGAAGGGTTTAGCAGAAGCTTTTGCAGACCTCAACAAGCTCCTTAAAAAGTTTGAAAACATGGACCCCAACACCGAAAGGTTTTCATTAATAGAGAGGAATGTTCATGGTGCATTATCTGCTTACAAGCAAATCTATGATGAAAAAAAGAAACAAACCAAGCAAACCACCATGGACATATTTCTGAAAAGAGTGACACCTCCTCAAGAAGAGCCTCAGGCAGGTCCTTCAGGAGGTATTCCAGAAGAAGGCATTGTTATCATAGGAGATGACGGCTCCATGCGCGTTATTGCCCCTGAAGACCTTCCAGTGGGACAAGATGTGGAGGTGGAAGACAGTGATATTGATGATCCTGACCCTGTGTAGGCCTAGGCTAATGTGTGTGTTTGTGTCTTAGTTTTTAACAAAAAAGTTTAAAAAGTAAAAAAAAAATAAATTTAAAAATAGAAAAAAGCTTATAGAATAAGGATATAAAGAAAGAAAATATTTTTGTACAGCTGTACAATGTGTTTGTGTTTTAAGCTAAGTGTTATTACAAAAGAGTCGAAAAGTTTAAAAAATTAAAAAGTTTATAAAGTAAAAAAGTTACAGTAAGCTAAGGTTAATTTATTATTGAAGAAAGAAAATTTTTTTAAATAAATTTAGTGTAGCCTAAGTGTACAGTGTTTATAAAGTCTACGGTAGTGTACAGTAATGTCCTAGGCCTTCACATTCACTCACCACTCACTCACTGACTCACCCAGAGCAACTTCCAGTCCTGCAAGCTCCATTCATGGTAAGTGCCCTATACAGGTGTACCATTTTTTATCTTTTATACCGTATTTTTACTGTACCTTTTCTATGTTTAGATACGTTTAGATACACAAATACTTACCATTGCGTTACAATTGCCTACAGTATTCAGTACAGTAACGTGCTGTACAGGTTTGTAGCCTAGGAGCAATAGGCTATACCATATAGCCTAGGTGTGTAGTAGGCTATACCATCTAGGTTTGTGTAAGTGCACTCTATGATGTTTGCACAATGACGAAATCGCCTAACGACGCATTTCTCAGAACGTATCCCCGTCGTTAAGCAACGCATGACTG

>KMARD1

CAGTCGTCCCTCGCTATATCGCGGTTCACTTATTGCRGCTTCACTGTATCGCGGGTTTTTTAAAAATATATATCTAATTCTGTATCGCGGAGTTTTCGCTATATCGCGGGATTTTGCGGTATATAGGTATTTATATAATATTTATGATTTTAATTATTTTTGCCTAAAAAATTTAAAACAATATAAAAATATAAAAAATGTTAATTCAAACATATTAAAAGAATATTAAATCGAAGCAAAATACAGTATGTATTTCATCAGCGGATGAATACCGCGCTGTACACAATGGAAACGCAGTACGCCACTACCGCTTGCGCAAGTTTGCCAACGTGAGATTGTACACGGCACACTATTGGCTGATGGAATGGAAGGCGACCAACCACAGCACTGTGTTCTGTATCCTGGGCGCTGATTGGCTCAGTGACCATAGCATCGGTCTGTTTGCTCTCCCGCACTGTGCCCGCACGTTCAGCATCTCTCCTTGTCCACTTCACATCAACGTCTGATCACTGCACGTAGTGTTGCTCTGCGGTGTTGTGTTTTTGTGAAATTTTCGTAAAAGTTTGAATTTATTTC**AAG|CCC**TACGATGCCACCCAAACGTTCTGCTCCTTCTAAGGCTTCTGACAGTGAACCCAAGCGCCAGAGGAAGATGCTTACCATCAAAGAAAAGGTGGAACTTCTCGACATGTTAAAGGAAGGCAAAAGCTATGCGGCTGTAGGCCGCCATTATGGGATCAACGAATCTACTGTTCGCTACATAAAGAAAGACGAGAAGAATATTAGGTCTACGGCAACAATGACTTTTAACAAGACTGCAAAGAGGGTGGTCGCTTCCCGCAATAAGGCCATTGTGAGGATGGAGTCTGCATTGGCCCTGTGGATAAATGACTGCAGGAAGAAGAATATCTCACTGGATACCAACACCATCCGGACGAAGGCCAAGAAACTTTATGATAGTTTTGCGGAAAGCACGGACGTTCACGACGGTGACGAGGATGAAGACGAGGATGACGACACAGAAGCAGGACCGTTGAGTGCTTCTCCCACCAGACCAACCCCATTCAGTGCCAGCAAGGGCTGGTTTGACAAATTTCAGAGACGCTTTGGACTCAAGAGCGTTTCTCTGCATGGAGAAGCTGCCTCTGTGGATAAAGCCGGAGCCGAGGAGTACGCGAACGACATGTTCAAATCCATCATCGAGGAAGGGGGCTATAAGCCTGAACAAGTTTTTAACATGGATGAGATGGGCTTATTTTGGAAATGGATGCCGTCTCGCACTTTTATTATGAAGGACGAAGCCAAAGCCCCTGGTTTTAAGGCCCAAAAAGATCGAGTAACTCTGATTATGTGTGGGAATGCTGCGGGCTTTATGATAAAGCCAGGGCTTATTTATAAGTCAAAAAACCCAAGGGCCCTAAAAAACAAAAATAAGAATGTGTTGCCTGTGTACTGGATGCGCAACCCCAAGGCCTGGATCACGAAACTCCTCACGGCTGATTGGTTCCACCAGTGCTTTATCCCGGAGGTTAAGCTTTATCTGGCAGAGAAAAACCTCGAATTCAAAGTGCTTCTGATCATGGACAATGCTGGAGGCCATGCTGTCGATCTGTCATATAATGGCGTCCAGATAGAGTTCTTGCCGCCGAACACCACATCGCTCATTCAGCCCATGGATCAAGGCGTTATTCGCGCTTTTAAGGCACTCTACATGCGAAATGCCCTGCAAAACCTGGTTGAAGCTATGGACTCGGATGAAAACTTCTCACTAAAGGCGTACTGGTGTGACTACAATATTGCATTGTGTCTCCTAAATATTCAGAAGGCCATAAAGGAGATGAAGAGTGAAACAGTAAATGCCAGCTGGAAAAATTTGTGGCCAGAAGTGGTCCACGATTACAAGGGATTCTCTCCTGATGAAGTCCATCACTCTGCGGTAGCTAAGGCCGTGAAGCTGGCAAAACTACTGGGAGGAGATGGCTTCAATGATATGACTCCTGACGACGTGAATGACCTGATCGACGTCCACTCACAACCGCTGACGGACGAAGACCTGACGGAGATGACGAAGTCAGCAAGTGAAGAAGAGGATGAGGAACAAGACGAACTACGGGTTGGAGAAGAGGAGGAGGTTGGCCTAACACTCGATCGTCTTGCAACCATGGTCAGAATGGCCAAAGAACTTCAGCGAGTGGCTGAAGAATGGGACCCCCAGATGCTTCGTTCATTGCACTTCGTAAATACAATCGAGAGTGGCATGTCAGTTTATAAAAACCTTCTCACTCAGAAGAAAAAGCAGCGCCAGCAACTCCCCATAACTATGTTCATCACTCGGAAGAAGAGATCAGTTACGGCCCAGGGCGAAGATACTGCACCACCTGATGAAGCGCCTGATGAAGTGGTGCCTTCAGAAGAGCTGTAATGCTCTTCTTGGCTGTGCAGTACATTCATCTCATCGTCATCACCTTCATCGCCATCAGTACTGCACAGCTACATAATTCATGATCTTCATCATTCAAGTTGTAGTATACTTCACTGGCGAGTACCCGTATAGAATTTTACATGTTGTTAAAATTACGTAGGTTTAAGAGTGTAGAAAGTGTTTAAGAGCATATGAAGTGTTTATAAGAGTGTGGGAAAGGTTAATAAGAGAGTGGGAAAGGTTTATAGGAGTGTGGGAAGGGTTTATAAAGCCTTAAAATATATATAAATAATAAAATAAATATAACGCCGCTACTTCGCGGATTTTCACCTATCACGGGGGTCTCTGGAACGTAACTCCCGTGATAGGTGAGGGATCACTG

>KTIGD3

CAGACAGTTCTCGCTTTACGTAAATTCGCGTTACGCAAATTCGACTTTACGTAAAGAATTCTGAAAGAAA

TCCGCATTCCAAAAAAAATTACGAAAATTCCTTACGCAAAGCAGACGCTGTACGATAAGACGCTGTATGATGCAGGCAGCGTAAACGGTGGTGGGAAGCTGAGAAGCATCTCTAAGTCAGTTCCCGCGCGTCTGCAGACAGTGAGTGATACTGTGCCTCTCCCTGTGTTCAA**TAG|CGA**TCCTTATATTTTCACCATGGCTGGCGGCAAACGTGCTCTTGAGGGGAAAAGTGAACCATCTCGTAAGCGTAAGCCAATAGACTTGGAAATGAAAATGAAAATAATTAGAAAATATGAAGGAGAACAAAAATTGTCATCGATAGCACGTGAACTTAGTCTTGCGGTGTCGACTGTGAACACCATAGTGAAGGATGCTGCCCGGATAAAGGAGCATGCAAAAGGCAGTGCTTGCATGAAGTCGACTTATCTGTCGAAACAAAGAGAAGGTGCTATTCTAGAGATGGAGAAACTATTAACTATGTGGATAGAAGACCAGACTCAGAAGAATATTCCTTTAAGCTTGATGACAATTCAAGCCAAGGCTCGAAGCCTCTTTGAAGATTTAAAGGTAAAGTACCCTGAAGGAACACAGGTATTTACAGCAAGCAGTGGTTGGTTTGCACGATTTAAAAATCGTGCAGGATTTCATAATGTGCAAGTGTCAGGAGAGGCTGCTAGTGCAGATATTGAAGCAGCAAAGAAGTTCCCAGAGTTCCTGCAAAAAATAATAGATGAAGGTCCCTACTTGCCGGAGCAAATCTTTAATGTAGACGAAACCGGCCTATTTTATCAAAGAATGCCATCTCGAACCTGCATCTCGGTGGAGGAAAAAAAAACTATGCCGGGGCATAAAGCTTCGAAAAATAGAGTTACTCTTCTGCTTGGAGGGAATGCATCTGGGACTTGCAAATTAAAACCTGTTCTTGTGCACCACTCAGAAAATCCCAGAGCATTGAAAGGAATCAGCAAGGCAACACTACCTGTGCATTATTTCGCTAACCCTAAGGCATGGATAACCCTTATCATCTTTGAACAATGGTTTATGCACTGCTTCATTCCCGAAGTGGAGAAGTTTTGTAATGATAATGGCATTCCTTTCAAGATTCTACTTATTTTGGATAATGCCCCAGGTCACCCTCCCCATCTGGATGATTTTAATGAAAATGTAAAAGTGGTCTACCTTCCCCCTAACACTACCTCATTATTACAACCTATGGATCAGGGAGTGATTGCGAATTTCAAGGCGTATTATTTACGCACTACATTTGCCCAGGCAATTGCAGCTTTGGATGCAGATAAGGACTTGACATTGAGAGATTTCTGGAAGTCCTATAATATTTACCATGCAATCCAGAATATTGCTAAGGCATGGGAAGATGTAACAGAAACATGTATGAAGAGAGTTTGGAAAAAAGTGTGCCCGCGGTTCGTTCCCGATTTTCGTGGATTTGACAAAGATGAAACGTTAAAAGAAATGAGTAACAAGATTGTAAAATTAGCACAAGAATTGGAGCTAGAAGTGGATGAGAATGATGTGGAGGAGTTGATTGAGTCTCATGGAGAAGAACTGTTGAATGAGGATCTGATTGAGCTGCTAGCTGCAAAGATTGCAGAAGAAGAAAGGGAAGCTTCGGAACCCGAGGTCGAACCAAAAAGGTTCACAACGAAACAGTTGGCAGAAGCATTTCATCATTTGAGTGAATTTTTTTCTTGTATTGAAAAGATGGATCCAAATACTTCAAGATTTTTAAAAGTTCAAAGACTAGTTGAGGATAATATTGCTTGCTATCATCAGATATATGAGGGAAAAAAGAGAGCCACTGTCCAAACATCTCTCGATAGCTTCATAAAAAAAGTTGCGCAGCCAACCACCAGTGATAAAGGCTAGGTGCAGTGGGTGGGGAAGGGGCAACATACTGCACACCTATGTTAACTTTACTGTACAGTACTGTGCATACTTTATCATTAACTTTACCTTTCATTTCATAATTCTGCTTATTATCATGGTACAGTATTTAATATGTCTGCATTTTAGTACATTTTATGACTTGCATAAAGGTATTATTTTGTGTATGCCTTTGTTGCATAGGCTAAGTGAAGTGGGTGGGTAGGGGCAAATTTGCATTACGTAAAAATCGCTTTATGTAGAGGTCTGCGGAACGGTCCACTTACGTAAAGCGAGAACTGTCTG

>KMARD4

cagtacaagccttgttatccggcactttaatatccagaaaactcggttaaccggcatctgccatagacggcaatacaattcccccttcagttaaccggaaaaattccgatagccatccggcatcattttccagcgcttgtggaagtcgcatttgcaaacttgcggcagtaaaggctgttcttgtcctgcctctgagtacartacwgtatcatcaycggatattatctgacccatcctg**cag|cta**tggctagcaatacaccagcaacaggcaaaggacagaaaagaaaacgtgttgtgctgacgataacccagaaattggacctaattaggcgccttgaaagaggagaaaacaggagccaattgatgcaagagtttggcgttggctcatcgacaatctacgatattaaggtgcagaaggcagawctgatgaaattcgtgtcaagtgctgagacgagcaaagcaattgagaaacgccgtactttgcacaaaccaaaattggagcaattagacaacgtgctctatgaatggttttccttgaagcgttctgagggagctccaatttccggcccaatgttgatcgaaaaggccaaggacttttatcagcagatggaaatgaccgagacctgcgttttttctgatggttggctggcatgctttaagatgcgacatggtatcagaaggcttgatgtctctggagagaaacagtctgctgatcatgaggccgcagaaaaatattgtgtgttttttaagaagctgattggggagcatggcataaccccagagaacatctacaatgctgatgaaatgggactcttttggcgttgcctgccgtcttcaacccttgctggtgcaggtgaggttggtgcacctggattcaagaggaacaaggaccgcatcactctgcttacgtgtgcaaatgctgctggcacacacaaaatcaagttgctagtggtttcaaaattcaagcggccaagagcattcaagggtattgtccatctgccagtggagtacaaggcacagcctaatgcatggatggacaaggacatatttttgtattggttccaccatatctttgttcctgctgtraaagaccacctgaaagaacttggagtgccagaagacacaaaggttattctggttttagataactgtcgagctcacycttctgaagcagagcttgtgtctggcaatattttcacggtcttcctgcctgcaaatgtcacctcacttattcagccaatggatcagggaattattcaaaacctaaaagtcatctacagaagggattttatgaggaagctgctcaactttgaaggaacaatccaagagtttcagtcacggtacagcatcaaggatgcaatcttcaatgcggcatgtgcctggtctgctgtcaagcctacaactttaaagagagcatggagaaaattgtgggcggaagtgatatttgtggaggggtcctctgatgaagaagaatttcaaggcttcaacgtaagaccaaaaaggagagcgctacaggagatacttgatgtccttcaaaatggcgaccccacaaacccaatcactaaactgaaggataacgaagttgaagagtgggtaaacgtcaatcaaggtctagatgttacaaaaaccctgacagacwctgaaattatagaaatggttgtccatccggagagaaatacctgtactgcagatgaaatagattcagaggaagaagaccatagtgaagaaaccaaagtctcttgggctactgctgcacagtgtctggaaactttggtgaagtttgccgagcagcagtcttcttattctgctcaggaagtcatgcagctgcacgttatccacaacaatttcctcaagaaaargcaactgacttgcaggcaggcagacatctgcaagatgcttcaaaaagcttctgcagcttgtgccactggtccagcaccatcaagatggacagcagaagttattgatrttgatgaccctgaggcartagagwagcagtaaattacggtactgtwtacagtaatagtgttaaaagrtgtagtgttaagtaaggttatttcatttcttaaacatttttgcatttggtgaaaaagctcaag

>KHATD2

Ttttt**cag|gaa**caaagaagagaccctttgttatttctttccaatgattgtggagagtggccactcaaaataaatgatgaggcacggaaaataattgttgagcgaggaccacagcaagtcagaggcataaagtttcctaaggacattcatggcagaaaattttccccatttcattattcaaggaagttgtgtaatggagagcgtgtcaacagatattggctacagtattctgtatctaaaaatgcagtgttttgtattacttgcaaaatttttggaaacgacacatccagtcttgcaggtagccaagggttttctgactggcggaacttgagcaggcttttgagcagtcacgagaaatcccgtcctcatatgaaaaaccgttcatcttggcgtgagctctcgcaacgtttgcatttaaacaaaactattgatgcagagcatgaaagacttattaatgctgaaattaaacattgggatcaaattctgaaacgcttgctatgtgttacacggtttttgggggttcaaggattgccatttagagggacaaaagatgttctatttgaacctaataatggcaactttttaaaattgatagaacatatagcacagtttgatgatccgatggctgaacatgtgagaagaatcacttccaaagaaacacatgtccactatttgagtaaaaatgtccagaacgagtttatttctttcttggcaggcaaggtccagaataatattttggaacaactacatgaggcaacatattattctattatattggactgcacaccagatattagcaacaccgaacaaatgacgttggtggttagatttgttacatgtaaagcaaatgaagatatcttgattaaggaacattttttaggctttgttcctgttgccagtccttcaggtgaaggtatgacagaaatattactccatgagttggaagcacgacgtattccattaaaaaacatgagaggccagggatatgataatggttctgcaatgaagggaaaacatgttggagtccagaggaggatccttgatttaaatcctagagccttttatgtaccatgtggaaaccactccctgaacttggtcataaatgatgcagctatgtcttgcaagattgcagcagactgtttcgccaccgtacaagacctctataaccttttttcaggttccccagtaagatggggtactttgttgaagcatgtttcaactctcacactcaaaccacttagcagtactaggtgggaaagtagaatcgaagcattgttgcctttgaggttccatattgaagaagtatatgatgccgtatatgaagcttctcatgatcagaaatttgatggactctgtaggagccgtgctggtgctcttcttaaaagactgcaaagcttcacatttctgtgcagcatagtgacatggcatgagatcctgcacaagataaatatggttagtaaacagttgcaaaaagtgtcaatcgatcttcaaaattctatggctcttattaagagtgtgaaaagttttctagaaaggatgagatctgaagaaggtctaaatagcatcattacagatgcaaaggaactggcagaaaaaatcgatgctactgccgactttgaaaacgaacaggaagctcgaccgagaaaagtaagcagacaaagtgaagatgaagctgcactttcaagtgagagtgaagatgaagctgcattttcgtatgaatgtaaagatgaagctgtacattctggcaaagagtcttttaaagtgaactttttttttgttgtgcttgacacagcaatatcctcattaaaggagagatttcagctaatggacaaccatagtggaagtttcaaatttctgtatgacatttcaagccttgggaaatgttggaatgaaaaagaattgaagtacgcctgtcaacgccttgaaactgttttgacagatggagaagaccacgatgtgaatgctggtgatctgtatacagagctacaattacttgcagatatgcttccccctggaagtctcccagctgatgccttgtcttttataaacaatcatggttcggaggatgttttcccaaatgtctacactgcattgagaattctgttaacacttccaatatccgtggccagcggtgaaaggagtttttcaaaattgaaacttataaaaaattacttaagatctactttgagtcaagagcgcttgagtggactatcaaccttggccattgaaaatagcttgttggatgacatggacacagattccctagtacatgagttttccaaattgaaagtcagaaaaatcaggttttaatttttacaagtgttcaagttaatttgtgttaaggaaaactggatgttaaaaaatccactcaagtgaaggggccagaggctggataggttgccacccccatgacaatggtccaggctggcatggcacttctgaaggacaagagttcgcttcctgaaaaa

>TIGD-G1

cagtagtacagtatctttatttc**aag|ccc**aggatgtctggaagcaagcgtaaaagcagcagtgatgtagctggtactgctaagaagtgccaagtgataacaatggaaacaaaagtgaaaataattgagagagtggagcaaggtgaaaagatggtagacatcgctcattcttataacatgaatcgttcaaccattggcactattctaaagaacaaggacaagatcatggaacatgtgaagtctgctgtgccgatgatgtcaacaataatatcaaagaagtgtgggaaagtgatggaggagatggagaaacttctcagtgtgtggatgcaggatcagcatcagcgtcgagtcccgctcagcttaatgctgattcaagagaaagctaaaagcctttatgaagacttgaagaagaaacacggcgaagaatcagagggcacatcttttaatgccagccatggctggtttcatcggttcaaggctagagccaaccttcacaatgtaaaagtaagtgacgaggcagtgagtgcagatacggtagctgcccgggaatttcctgaaatccttcaagaaattattgatgaaggtgcgtatttaccagagcgggtttttaatgtggatgagacaggactgtactggaagaggatgccagaccgaagttacatcagtaaggaggaaaagttgatgccaggctataaagcagcaaaggataggctaactctgttgtttggtggcagtgcttctggcaatatgaagctgaagcctctcttagtttatcattcagagaacccaagagcccttaaaaacatagccaagggctctcttcctgttgtgtggaagagtaaccccaaagcctgggttacacaggccattttccaggactcgtttttccaccactttatcccggaggtagagaaatattgcttggagaaggacatcccattcaacattcttttgctgctcgacagtgctctgggccaccccccattcatggacgactttcatcccaacgtcaaagtagtgcatctgccaccgaatactacattgcttatccaacctatggaccagggagttatagcgactttcaagaaatattatttacgtcacacttttcgtcaggcagtaaaggtgagtgatgaatcaggaacaaccttgcaacaattttggaaggactataacatctacaaggccataaaaaacattgactttgcttggtgtgaggttacagccgtcaccatgaatggggtttggaagaacctttgcctgcagtttgttcatgattttcgtggatttgagaaggtggatgaggagtccaaagaggtcttcagcaacttagtgaccctcagtgagaagctggagctagatctgcaagaggacgacttcactgaactccttgctgtgcaacacgaggagctyactaatgaagacctgatggaactggaggcccagagaaaggatgaagagagacaagaggaagaagtaaaacgagctgaagaactaaagagattcacgatgcaggaaatggcaagggggttttctttatttgaggaggcactgttagtttttgagacacaggacccgaatgtagaacggtacacgaaggttgcagcagccgttcagaatgcaatccagtgctaccgtgtcatctatgatgagaaaaaaagaaactactacccagacatcactggatcattttttttaaagggtagatagaattgaatccagcaaggaaccagaacctgtgccatcaacgtcaggcatgagtgaaattgcagcttgccctccgtctcctattgctgatgatccttcagctctaccatctcccacctcctctccctcctccagtcagtaactcttcttgcctgttcactcgatgccagyccctgtatgccagctgttgtactgtactactg

>TIGD-G4-Zscan29

cagtaataccttggttcacgaacgcttctgttcgcgaacaattcggttcgcgaacagaagcgttcggcaaaattttgcttccgttcgcgaactccgcatcgggtgacgaacacgaccggggccatcttccccgcacgaacacgaacgtggccatcttccccgctctgacacgtggtcgtggcttcctctagtcgagatcgcggtcgcagacagacctgtgcacacttgctgtgaagcttttttgtgtgctttttagcacatacagtactgtacagtattattcaatccagcatggctccaaagaaagtgcagagcgagcatggtggtaaaaagaaagtgcagagcggtaatgaaggtaacaagaatgtgcagagtggtaatgaaggtgacaagaatgtgcagagcaataatgaaggtaacaagaaagtgcagagcaagaatggtaagaag**aag|gtt**ttgaagaaaataaccattgaattgaagaaggaaattgtagaaaagcatgagcgtggtattcgtgtgactgatctggcctcggagtacaagatggcgaagtcgacaatctcgactattttgaagaataaagatgccatcaaaggagctgatgtggcaaaaggagtaacgatgttaaccaagcagaggacacaagtgctggaagaggtggaaaaacttttgttagtgtggttgaatgagaaacagctggcaggtgatagcgttagtgaagctatgatctgtgaaaaagccaggaaattgcacagtgatttactgcaagaaaacccctctacaagtgcaacaagtgatgaatttaaagccagtagagggtggtttgataaattccgcaagagaagtggcatccacagtgtgataagacatggtgaggcttccagttctgacaaggccgctgcagaagcctacaagaaagaattcgcggaattcatgaaggcagaaggatacatcgctcaacaagtgttcaactgtgatgaaacagggctcttctggaagaagatgccgaacagaacctatatcacgcaggaggaaaaggcactgccggggcacaagcccatgaaggacagattgacycttttgctgtgtgcgaacgcaagtgccgatctgaagattaagccactactggtgtaccattcccagacccctcgtgcattcaaggaacagaatgtgaacaaggccagactgcctgtcacgtggagggccaatgcgaaagcttgggtcacaaggcaattgttcatggaatggctgcacgaggtgttcgcacccaccgttaagaaatatctttctgacaaccaactgcctgaaaggtgccttcttctgatggacaatgccccggcacaccctccagccttggtggatgatatggacgctgagtatgacttcataaaggtcaagttcctcccccccaacacgacacctcttctgcagcccgtggaccagcaagtgatctgcaacttcaagaagctgtacacaaaggcgctcttcaccaggtgttttaatgtcactgaagagatgtccttgactctgaaagagttctggaagaaacatttcaatgtcctccactgcatcaacctcattgacaaagcctgggaagatgtcacgtaccggaccttaaactcagcctggaggaaactgtggccagaatgtgtcactgaacgtgactttgaagggtctgatgcacaggtagtggaggagattgtgtccatgggcaagagtatgggtctggacgtcgatggtgcagacatagaggagcttgttgaggaccataaggaggagctgacgacggaagaacttgctgaactccagagtgagcagcagaaggtgcttgttgaggagcattccactgaggaagaggaagacagggaggaggttagcagtgatgtgataaaatccattatggaaaaatggaatgagtgccaggatttctttgagaagcaccaccccaacataactgtaatgaacagagtgttgaatctgatgaacgataacgtggtctctcatttccggagggttatgcagcgcaggaaaaaacaagtgacattagacagatttttcatcaaaattgaccctgcagctaagagacaaagaagagaggaaacccctgaaggagatctccccgatgtccttatggagggggactctccctccaaacaataacctcctccccacccaccttcctgtcctccacgccagaagtcgcctcaagcaaggttagtgttctctctcttcatactgtactgtactgtactttgtttcatatttttgtggttaaaaaatacatatggttcagaacggattaaccttatttacattgaaccttatggggaaatttgattcggttcgcgaccaattcggttcgtgaccagaatcatggaacgaattaagttcgtgaaccgaggtaccactg

>MARD-G1-PUM2

CGAGGGGTCTTCAAAAAGTTCATGGAAAATGCGTATTATGAAAAAAATAATTACGGATTTTAAAAATTTTTTTGCACCAAAATAAACTCATACTAACTTGTTATAACATGTCTGAACAGGATCTAGTTTGAGGCACTAAGAAGGATAAGACATCAGTTTGAAAAGAGCCCCTATCAGAGCAACATGAATTCTGCTAAAATTGAAGCAAGAACAAACATCAAATTTATGGTGAAGCTTGGGTGGAAGAATGGTGAAATCGTTGATGCTTTACGAAAAGTTTATGGGGACAGCGCCCCAAAGAAATCAGCAGTTTACAAATGGCTAGCTCGTTTTAAGAAGGGACGAGACGATGTTGAAGATGAAGCCCGCAGCGGCAGACCGTCCACATCAATTTGTAAGGAAAAAATTAATCTTGTTCGTGCCCTAATTGAAGAGGACCGACGATTAACAGCAGAAACAATAGCCAACACCATAGACATCTCAATTGGTTCAGCTTACACAATTCTGACTGAAAAATTAAAGTGGAGCAAGCTTTCTGCTCGATGGGTGCCAAAACTGTTGCGCCCAGATCAGCTGCAGACAAGAGCAGAGCTTTCAATGGAAATTTTAAACAAGTGGGATCAAGATCCTGAAGCATTTCTTCGAAGAATTGTAACGGGAGATGAAACGTGGCTTTACCAGTACGATCCTGAAGACAAAGCACAGCCAAAGCAGCGGCTACCAAGAGGTGGAAGTGGTCCAGTCAAAGCAAAAGCGGGCCGGTCAAGAGCA**AA|GGTCATG**GCGACTATTTTTTGGGATGCTCAAGGCATTTTGCTTGTTGACTTTCTGGAGGGCCAAAGAACGATAACATCTGCTTATTATGAGAGTGTTTTGAGAAAGTTAGCCAAAGCTTTAGCAGAAAAACGCCCGGGAAAGCTTCACCAGAGAGTCCTTCTCCACCACGACAGTGCTCCTGCTCATTCCTCTCATCAAACAAGGGCAATTTTGCGAGAAGTTTCCGTGGAAATCATTAGGCATCCACCTTACGGTCCTGATTTGGCTCCTTCTGACTTCTTTTTGTTTCCTAATCTTAAAAAATCTTTAAAGGGCACCCGTTTTTCTTCAGTTAATAATGTAAAAAAGACTGCATTGACATGGTTAAATTCCCAGGACCCTCAGTTCTTTAGGGATGGACTAAATGGCTGGTATCATCGCTTACAAAAGTGTCTTGAACTTGATGGAGCTTATGTTGAGAAATAAAATTTTTATTTTTTATTTTGATCTTTTAATTCCATTTTCCATGAACTTTTTGAAGACCCCTCG
