## Supplemental Data 2 for "Recurrent evolution of vertebrate transcription factors by transposase capture"

>*Myotis_lucifugus_Mlmar1*-KRAB

CTAGGTGTACCAGTTAATAATGCGGATTTTTTTCAATAGTTGGAGTTACACATATGTTGATATATATGTGATTTGATATGTATGCTATTTTGTTGTATTGACAGCAAGCTTCAAAACTTCATATGTCAAATTTGCTGAAGGTGTTAACATCATAGATATTTTTACACTTAAAAATGTCGAATTCGTGCCAAAAAAAGAGCATTTGCGGGAAGTTTTAATTCATTACTTTATTTTGAAGAAAAGTGCTGCTGAAAGTTATCGTATACTTCGGGAAGCTTATGGTGAACATGCTCCATCTCAAGATACTTGTGAACGCTGGTTTAAACGGTTTAAAAGTGATGATTTCAATGTGAAAGACAAAGAACGTCCAGGTCAACCAAAAAAGTTTCAAGACCAACAATTACAAGCATTATTGGATGAAGATGCAGGTCAGACTCAAAAACAACTTGCAGAAAGATTAAATGTTGCTCAGCAAACAATTTCCGATCGTTTACAAGCAATGGGAAAGATTTTAAAGGAAGGAAAATGGGTGCCACATCAACTGAACAAAAGACAAATGGAAAACCGAAAAGTCATCAGTAAAATGTTGCTTCAACGGCACGAAAGAAAGTCTTTTTTGCATCGAATTGTGACTGGCGATGAAAAGTGGATTTATTTTGAGAATCCCAAATGCACAAAATCATGGGTTGATCCAGGTCAACCATCAACATCGACTGCAAGGCCAAATCGCTTCGGAAAGAAGACAATGCTCTGCGTTTGGTGGGATCAGGAAGGTGTGGTGTATTATGAGCTTCTAAAACCAGGTGAAACCGTTGATACTGATCGCTATCGACAACAAATAATCAATTTGAACCACGCTTTGATCGTGAAACGACCAGAATGGGCCAGAAGACACGGCAAAGTAATTTTGCTTCATGATGATGCACCATCACACACTTCAAAGCCAGTTAAAGACATGTTAAAAGATCTTGCCTGGGAAGTATTAACCCACCCGCCATATTCACCAGACCTTGCTCCTTCAGATTACCACTTGTTCCGATCGATGGCACGCGCACTTTCTCAGCAGCACTTCAAAACGTACGAAGAAGTGGAAAATTGGGTCTCTGAATGGTTTGCCTCAAAACAAGAAAAGTTCTATTGGGACGGTATCCACAAATTACCTGAAAGATGGGGGAAATGTGTAGCTAGTGATGGACATTACTTTGAATAGAGCACTTTTGATGTTTCTCTTGAAATTATCATGTTTTCTTTGATTACAAAATCCGCATTATTAACCGGTACCCCTGG

>*Myotis_velifer_Mlmar1*-KRAB

CTAGGTGTACCAGTTAATAATGCGGATTTTTTTCAATAGTTGGAGTTACACATATGTTGATATATATGTGATTTGATATGTATGCTATTTTGTTGTATTGACAGCAAGCTTCAAAACTTCATATGTCAAATTTGCTGAAGGTGTTAACATCATAGATATTTTTACACTTAAAAATGTCGAATTCGTGCCAAAAAAAGAGCATTTGCGGGAAGTTTTAATTCATTACTTTATTTTGAAGAAAAGTGCTGCTGAAAGTTATCGTATACTTCGGGAAGCTTATGGTGAACATGCTCCATCTCAAGATACTTGTGAACGCTGGTTTAAACGGTTTAAAAGTGATGATTTCAATGTGAAAGACAAAGAACGTCCAGGTCAACCAAAAAAGTTTCAAGACCAACAATTACAAGCATTATTGGATGAAGATGCAGGTCAGACTCAAAAACAACTTGCAGAAAGATTAAATGTTGCTCAGCAAACAATTTCCGATCGTTTACAAGCAATGGGAAAGATTTTAAAGGAAGGAAAATGGGTGCCACATCAACTGAACAAAAGACAAATGGAAAACCGAAAAGTCATCAGTAAAATGTTGCTTCAACGGCACGAAAGAAAGTCTTTTTTGCATCGAATTGTGACTGGCGATGAAAAGTGGATTTATTTTGAGAATCCCAAATGCACAAAATCATGGGTTGATCCAGGTCAACCATCAACATCGACTGCAAGGCCAAATCGCTTCGGAAAGAAGACAATGCTCTGCGTTTGGTGGGATCAGGAAGGTGTGGTGTATTATGAGCTTCTAAAACCAGGTGAAACCGTTGATACTGATCGCTATCGACAACAAATAATCAATTTGAACCACGCTTTGATCGTGAAACGACCAGAATGGGCCAGAAGACACGGCAAAGTAATTTTGCTTCATGATGATGCACCATCACACACTTCAAAGCCAGTTAAAGACATGTTAAAAGATCTTGCCTGGGAAGTATTAACCCACCCGCCATATTCACCAGACCTTGCTCCTTCAGATTACCACTTGTTCCGATCGATGGCACGCGCACTTTCTCAGCAGCACTTCAAAACGTACGAAGAAGTGGAAAATTGGGTCTCTGAATGGTTTGCCTCAAAACAAGAAAAGTTCTATTGGGACGGTATCCACAAATTACCTGAAAGATGGGGGAAATGTGTAGCTAGTGATGGACATTACTTTGAATAGAGCACTTTTGATGTTTCTCTTGAAATTATCATGTTTTCTTTGATTACAAAATCCGCATTATTAACCGGTACACCTGG

>*Myotis_occultus_Mlmar1*-KRAB

CTAGGTGTACCAGTTAATAATGCGGATTTTTTTCAATAGTTGGAGTTACACATATGTTGATATATATGTGATTTGATATGTATGCTATTTTGTTGTATTGACAGCAAGCTTCAAAACTTCATATGTCAAATTTGCTGAAGGTGTTAACATCATAGATATTTTTACACTTAAAAATGTCGAATTCGTGCCAAAAAAAGAGCATTTGCGGGAAGTTTTAATTCATTACTTTATTTTGAAGAAAAGTGCTGCTGAAAGTTATCGTATACTTCGGGAAGCTTATGGTGAACATGCTCCATCTCAAGATACTTGTGAACGCTGGTTTAAACGGTTTAAAAGTGATGATTTCAATGTGAAAGACAAAGAACGTCCAGGTCAACCAAAAAAGTTTCAAGACCAACAATTACAAGCATTATTGGATGAAGATGCAGGTCAGACTCAAAAACAACTTGCAGAAAGATTAAATGTTGCTCAGCAAACAATTTCCGATCGTTTACAAGCAATGGGAAAGATTTTAAAGGAAGGAAAATGGGTGCCACATCAACTGAACAAAAGACAAATGGAAAACCGAAAAGTCATCAGTAAAATGTTGCTTCAACGGCACGAAAGAAAGTCTTTTTTGCATCGAATTGTGACTGGCGATGAAAAGTGGATTTATTTTGAGAATCCCAAATGCACAAAATCATGGGTTGATCCAGGTCAACCATCAACATCGACTGCAAGGCCAAATCGCTTCGGAAAGAAGACAATGCTCTGCGTTTGGTGGGATCAGGAAGGTGTGGTGTATTATGAGCTTCTAAAACCAGGTGAAACCGTTGATACTGATCGCTATCGACAACAAATAATCAATTTGAACCACGCTTTGATCGTGAAACGACCAGAATGGGCCAGAAGACACGGCAAAGTAATTTTGCTTCATGATGATGCACCATCACACACTTCAAAGCCAGTTAAAGACATGTTAAAAGATCTTGCCTGGGAAGTATTAACCCACCCGCCATATTCACCAGACCTTGCTCCTTCAGATTACCACTTGTTCCGATCGATGGCACGCGCACTTTCTCAGCAGCACTTCAAAACGTACGAAGAAGTGGAAAATTGGGTCTCTGAATGGTTTGCCTCAAAACAAGAAAAGTTCTATTGGGACGGTATCCACAAATTACCTGAAAGATGGGGGAAATGTGTAGCTAGTGATGGACATTACTTTGAATAGAGCACTTTTGATGTTTCTCTTGAAATTATCATGTTTTCTTTGATTACAAAATCCGCATTATTAACCGGTACACCTGG

>*Myotis_sp_Mlmar1*-KRAB

CTAGATGTACCAGTTAATAATGCGGATTTTTTTCAATAGATGGAGTTACACATATGTTGATATATATGTGATTTGATATGTATGCTATTTTGTTGTATTGACAGCAAGCTTCAAAACTTCATATGTCAAATTTGCTGAAGGTGTTAACATCATAGATATTTTTACACTTAAAAATGTCGAATTCGTGCCAAAAAAAGAGCATTTGCGGGAAGTTTTAATTCATTACTTTATTTTGAAGAAAAGTGCTGCTGAAAGTTATCGTATACTTCGGGAAGCTTATGGTGAACATGCTCCATCTCAAGATACTTGTGAACGCTGGTTTAAACGGTTTAAAAGTGATGATTTCAATGTGAAAGACAAAGAACGTCCAGGTCAACCAAAAAAGTTTCAAGACCAACAATTACAAGCATTATTGGATGAAGATGCAGGTCAGACTCAAAAACAACTTGCAGAAAGATTAAACGTTGCTCAGCAAACAATTTCCGATCGTTTACAAGCAATGGGAAAGATTTTAAAGGAAGGAAAATGGGTGCCACATCAACTGAACAAAAGACAAATGGAAAACCGAAAAGTCATCAGTAAAATGTTGCTTCAACGGCACGAAAGAAAGTCCTTTTTGCATCGAATTGTGACTGGCGATGAAAAGTGGATTTATTTTGAGAATCCCAAATGCACAAAATCATGGGTTGATCCAGGTCAACCATCAACATCGACTGCAAGGCCAAATCGCTTTGGAAAGAAGACAATGCTCTGCGTTTGGTGGGATCAGAAAGGTGTGGTGTATTATGAGCTTCTAAAACCAGGTGAAACCGTTGATACTGATCGCTATCAACAACAAATAATCAATTTGAACCACGCTTTGATGGTGAAACGACCAGAATGGGCCAGAAGACACGGCAAAGTAATTTTGCTTCATGATGATGGACCATCACACACTTCAAAGCCAGTTAAAGACACGTTAAAAGATCTTGCCTGGGAAGTGTTAACCCACCCGCCATATTCACCAGACCTTGCTCCTTCAGATTACCACTTGTTCCGATCGATGGCACGCCCACTTTCTCGGCAGCACTTCAAAACGTACGAAGAAGTGGAAAATTGGGTCACTGAATGGTTTGCCTCAAAACAAGAAAAGTTCTATTGGGACGGTATCCACAAATTACCTGAAAGGTGGGGGAAATGTGTAGCTAGTGATGGACATTACTTTGAATAGAGCACTTTTGATGTTTCTTTTGAAATTATCATGTTTTCTTGTGATTACAAAATCCGCATTATTAACCGGTACACCTGG

>*Kerivoula_papillosa_Mlmar1*-KRAB

GACAGCAAGCTTCAAAACTTCATATGTCAAATTTGCTGAAGGTGTTATCATAGATATTTTTACGCTTAAAAATGTCGAATTCGTGCCCAAAAAAGAGCATTTGCGAGAAGTTTTAATTCATTACTTTCTTTTGAAGAAAAGTGCTGCTGAAAGTTATTGTATACTTCGGAAAGCTTATGGTGAACATGCTCCATCTCAAAATACTTGTGAACGCTGGTTTAAACGCTTTAAAAGTGATGATTTCAATGTGAAAGACAAAGAACGCCCAGGTCAACCCAAAAAGTTTCAAGATCAACAATTACAAGCATTATTGGATGAAGATGCAGGTCAGACTCAAAAACAACTTGCAGAAAGATTAAACGTTGCTCAGCAAACAATTTCTGATCGTTTACAAGCAATGGGAAAGATTTTAAAGGAAGGAAAATGGGCGCCACATCAACTGAACAAAAGACAAATGGAAAACCGAAAAGTCGTCAGTAAAATGTTGCTTCAGCGGCACGAAAGAAAGTCTTTTTTGCATCGAATTGTGACTGGCAATGAAAAGTGGATTTATTTTGAGAATCACAAGTGCACAAAAGCATGGGTTGATCCAGGTCAACCATCAACATCAACTGCAAGGCCAAATCGCTTCGGAAAGAAGACAATGCTCTGCGTTTGGTGGGATCAGGAAGGTGTGGTGTATTACGAGCTTCTAAAACCAGGTGAAACCATTAATACTGATCGCTACCGACAACAAATAGTCAATTTGAACCATGCTTTGATCATGAAACGACCAGAATGGGCCAGAAGACAAGGCAAAGTAATTTTGCTACATGATGGCGCACCGTCACACACTTCAAAGCCAGTTAAAGACACATTAAAAGATCTTGCCTGGGAAGTATTAACCCATCCGCCATGTTCACCAGACCTTGCTCCTTCAGATTACCACTTGTTCCGATCCATAGCACACGCACTTTCTCAGCAGCACTTCAAAACGTACGAAGAAGTGGAAAATTGGGTCTCTGAATGGTTTGCCTCAAAGCAAGAAAAGTTCTATTGGGACGGTATCCACAAATTACCTGAAAGATGGAGGAAATGTGTAGCTAGTGATGGACATTACTTTGAATAGAGCACTTTTTTTTTATATATATATATATTTTATTGATTTTTCACAGAGAGGAAGGCAGAGGGATAGAGAGTTAGAAACATCAATGAGAGAGAAGCATCGATCAGCTGCCTCTTGCACTCCCCCTACTGGGGATGTGCCCGCAACCAACGTACATGCCCTTGGCTGGGATCAAACCTGGGACCCTTGAGTCCGCAGGCAGATGCTCTACCACTGAGCCAAACCGGCTAGGGCTGAATAGAGCACTTTTGATGTTTCTCNTGAAATTATATATTTTCTTTAATTACAAAATCCACATTATTAAC

>*Eptesicus_fuscus_Mlmar1-*KRAB

CTAGGTGTACCAGTTAATAATGCGGATTTTTTTCAATTGATGGAGTTACACATATGTTGATATATATGTGATTTGATATGTATGCTGTTTTGTTGTATTGACAGCAAGCTTCAAAACGTCATATGTCAAATTTGCTGAAGGTGTTAACATCATAGATATTTTTACACTTAAAAATGTCGAATTCGTGCCAAAAAAAGAGCATTTGCGGGAAGTTTTAATTCATTACTTTATTTTGAAGAAAAGTGCTGCTGAAAGTTATCGTATACTTCGGGAAGCTTATGGTGAACATGCTCCATCTCAAGATACTTGTGAACGCTGGTTTAAACGCTTTAAAAGTGATGATTTCAATGTGAAAGACAAAGAACGTTCAGGTCAACCGAAAAAGTTTGAAGACCAACAATTACAAGCATTATTGGATGAAGATGCGGGTCAAACTCAAAAACAACTTGCAGAAAGATTAAATGTTGCTCAGCAAACAATTTCCGATCGTTTACAAGCAATGGGAAAGATTTTAAAGGAAGGAAAATGGGTGCCACATCAACTGAACGAAAGACAAATGGAAAACCGAAAAGTCATCAGTAAAATGTTGCTTCAACGGCATGAAAGAAAGTCTTTTTTGCATCGAATTGTGACTGGTGATGAAAAGTGGATTTATTTTGAGAATCCCAAATGCACAAAATCATGGGTTGATCCAGGTCAACCATCAACATCGACTGCAAGGCAAAATCGCTTCAGAAAGAAGACAATGCTCTGCGTTTGGTGGGATCAGGAAGGTGTGGTGTATTATGAGCTTCTAAAACCAGGTGAAACTGTTAATACTGATCGCTACCGACAACAGATAATCAGTTTGAACCATGCTTTGATTATGAAACGACCAGAACGGGCCAGAAGACAAGGCAAAGTAATTTTGCTTCATGATGATACACCATCACACACTTCAAAACCAGTTAAAGACACATTAAAAGATCTTGCCTGGGAAGTCTTAACCCACCCACCGTATTCACCAGACCTTGCTCCTTCAGATTACCACTTGTTCCGATCGATGGCACGCACACTTTCTCGGCAGCACTTCAAAACGTATGAAGAAGTGGAAAATTGGGTCTCTGAATGGTTTGCCTCAAAACAAGGAAAGTTCTATTGGGACGGTATCCACAAATTACCTGAAAGATGGGGGAAATGTGTAGCTAGCAATGGACATTACTTTGAATAAAGCATTTTTGATGTTTCTCTTGAAATTACCGTGTTTTCTTTGATTACAAAATCCGCATTATTAACCGGTACACCTGG

>*Eptesicus_furinalis_Mlmar1*-KRAB

CTAGGTGTACCAGTTAATAATGCGGATTTTTTTCAATTGATGGAGTTACACATATGTTGATATATATGTGATTTGATATGTATGCTGTTTTGTTGTATTGACAGCAAGCTTCAAAACTTCATATGTCAAATTTGCTGAAGGTGTTAACATCATAGATATTTTTACACTTAAAAATGTCAAATTCGTGCCAAAAAAAGAGCATTTGCGGGAAGTTTTAATTCATTACTTTATTTTGAAGAAAAGTGCTGCTGAAAGTTATCGTATACTTCGGGAAGCTTATGGTGAACATGCTCCTTCTCAAGATACTTGTGAACGCTGGTTTAAACGCTTTAAAAGTGATGATTTCAATGTGAAAGACAAAGAACGTTCAGGTCAACCGAAAAAGTTTGAAGACCAACAATTACAAGCATTATTGGATGAAGATGCGGGTCAAACTCAAAAACAACTTGCAGAAAGATTAAATGTTGCTCAGCAAACAATTTCCGATCGTTTACAAGCAATGGGAAAGATTTTAAAGGAAGGAAAATGGGTGCCACATCAACTGAACGAAAGACAAATGGAAAACCGAAAAGTCATCAGTAAAATGTTGCTTCAACGGCACGAAAGAAAGTCTTTTTTGCATCGAATTGTGACTGGCGATGGAAAGTGGATTTATTTTGAGAATCCCAAATGCACAAAATCATGGGTTGATCCAGGTCAACCATCAACATCGACTGCAAGGCCAAATCGCTTCGGAAAAAGACAATGCTCTGCGTTTGGTGGGATCAGGAAGGTGTGGTGTATTATGAGCTTCTAAAACCAGGTGAAACTGTTAATACTGATCGCTACAGACAACAGATAATCAGTTTGAACCATGCTTTGATTGTGAAACGACCAGAATGGGCCAGAAGACAAGGCAAAGTAATTTTGCTTCATGATGATACACCATCACACACTTCAAAACCAGTTAAAGACACATTAAAAGATCTTGCCTGGGAAGTATTAACCCACCCACCGTATTCACCAGACCTTGCTCCTTCAGATTACCACTTGTTCCGATCGATGGCACGCACACTTTCTCGGCAGCACTTCAAAACGTATGAAGAAGTGGAAAATTGGGTCTCTGAATGGTTTGCCTCAAAACAAGGAAAGTTCTATTGGGACGGTATCCACAAATTACCTGAAAGATGGGGGAAATGTGTAGCTAGTGATGGACATTACTTTGAATAAAGCATTTTTGATGTTTCTCTTGAAATTACCGTGTTTTCTTTGATTACAAAATCCGCATTATTAACCGGTACACCTGG

>*Myotis_lucifugus_KRABINER*_CDS

ATGACCAAGTTCCAGGAGCCGGTGACGTTCAAGGATGTGGCTGTGATCTTCACGGAGGAGGAGCTGGGGCTGCTGAACCCGTCCCAGAGGAAACTGTACCGAGATGTGACTCTGGAGAACTTCCGGAACCTGGTCTCAGTGGGGACTCAGCTCTTCAAACCAGACCTTATCTTCCAGCTGGAGAGAGAAGAAAAGCGTTTGATGGTGGAGACAGAAACCCAGAAAGATGGGTGTTCAGCAAGCTTCAAAACTTCATATGTCAAATTTGCTGAAGGTGTTAACATCATAGATATTTTTACACTTAAAAATGTCGAATTCGTGCCAAAAAAAGAGCATTTGCGGGAAGTTTTAATTCATTACTTTATTTTGAAGAAAAGTGCTGCTGAAAGTTATCGTATACTTCGGGAAGCTTATGGTGAACATGCTCCATCTCAAGATACTTGTGAACGCTGGTTTAAACGGTTTAAAAGTGATGATTTCAATGTGAAAGACAAAGAACGTCCAGGTCAACCAAAAAAGTTTCAAGACCAACAATTACAAGCATTATTGGATGAAGATGCAGGTCAGACTCAAAAACAACTTGCAGAAAGATTAAATGTTGCTCAGCAAACAATTTCCGATCGTTTACAAGCAATGGGAAAGATTTTAAAGGAAGGAAAATGGGTGCCACATCAACTGAACAAAAGACAAATGGAAAACCGAAAAGTCATCAGTAAAATGTTGCTTCAACGGCACGAAAGAAAGTCTTTTTTGCATCGAATTGTGACTGGCGATGAAAAGTGGATTTATTTTGAGAATCCCAAATGCACAAAATCATGGGTTGATCCAGGTCAACCATCAACATCGACTGCAAGGCCAAATCGCTTCGGAAAGAAGACAATGCTCTGCGTTTGGTGGGATCAGGAAGGTGTGGTGTATTATGAGCTTCTAAAACCAGGTGAAACCGTTGATACTGATCGCTATCGACAACAAATAATCAATTTGAACCACGCTTTGATCGTGAAACGACCAGAATGGGCCAGAAGACACGGCAAAGTAATTTTGCTTCATGATGATGCACCATCACACACTTCAAAGCCAGTTAAAGACATGTTAAAAGATCTTGCCTGGGAAGTATTAACCCACCCGCCATATTCACCAGACCTTGCTCCTTCAGATTACCACTTGTTCCGATCGATGGCACGCGCACTTTCTCAGCAGCACTTCAAAACGTACGAAGAAGTGGAAAATTGGGTCTCTGAATGGTTTGCCTCAAAACAAGAAAAGTTCTATTGGGACGGTATCCACAAATTACCTGAAAGATGGGGGAAATGTGTAGCTAGTGATGGACATTACTTTGAATAG

>*Myotis_velifer_KRABINER*_CDS

ATGACCAAGTTCCAGGAGCCGGTGACGTTCAAGGATGTGGCTGTGGTCTTCACGGAGGAGGAGCTGGGGCTGCTGAACCCGTCCCAGAGGAAACTGTACCGAGATGTGACTCTGGAGAACTTCCGGAACCTGGTCTCAGTGGGGACTCAGCTCTTCAAACCAGACCTTATCTTCCAGCTGGAGAGAGAAGAAAAGCGTTTGATGGTGGAGACAGAAACCCAGAAAGATGGGTGTTCAGCAAGCTTCAAAACTTCATATGTCAAATTTGCTGAAGGTGTTAACATCATAGATATTTTTACACTTAAAAATGTCGAATTCGTGCCAAAAAAAGAGCATTTGCGGGAAGTTTTAATTCATTACTTTATTTTGAAGAAAAGTGCTGCTGAAAGTTATCGTATACTTCGGGAAGCTTATGGTGAACATGCTCCATCTCAAGATACTTGTGAACGCTGGTTTAAACGGTTTAAAAGTGATGATTTCAATGTGAAAGACAAAGAACGTCCAGGTCAACCAAAAAAGTTTCAAGACCAACAATTACAAGCATTATTGGATGAAGATGCAGGTCAGACTCAAAAACAACTTGCAGAAAGATTAAATGTTGCTCAGCAAACAATTTCCGATCGTTTACAAGCAATGGGAAAGATTTTAAAGGAAGGAAAATGGGTGCCACATCAACTGAACAAAAGACAAATGGAAAACCGAAAAGTCATCAGTAAAATGTTGCTTCAACGGCACGAAAGAAAGTCTTTTTTGCATCGAATTGTGACTGGCGATGAAAAGTGGATTTATTTTGAGAATCCCAAATGCACAAAATCATGGGTTGATCCAGGTCAACCATCAACATCGACTGCAAGGCCAAATCGCTTCGGAAAGAAGACAATGCTCTGCGTTTGGTGGGATCAGGAAGGTGTGGTGTATTATGAGCTTCTAAAACCAGGTGAAACCGTTGATACTGATCGCTATCGACAACAAATAATCAATTTGAACCACGCTTTGATCGTGAAACGACCAGAATGGGCCAGAAGACACGGCAAAGTAATTTTGCTTCATGATGATGCACCATCACACACTTCAAAGCCAGTTAAAGACATGTTAAAAGATCTTGCCTGGGAAGTATTAACCCACCCGCCATATTCACCAGACCTTGCTCCTTCAGATTACCACTTGTTCCGATCGATGGCACGCGCACTTTCTCAGCAGCACTTCAAAACGTACGAAGAAGTGGAAAATTGGGTCTCTGAATGGTTTGCCTCAAAACAAGAAAAGTTCTATTGGGACGGTATCCACAAATTACCTGAAAGATGGGGGAAATGTGTAGCTAGTGATGGACATTACTTTGAATAG

>*Eptesicus_fuscus_KRABINER*_CDS

ATGACCAAGTTCCAGGAGGCGGTGACGTTCAAGGACGTGGCTGTGATCTTCACGGAGGAGGAGCTGGGGCTGCTGGACCCGTCGCAGAGGAAACTGTACCGAGACGTGACTCTGGAGAACTTCCGGAACCTGGTCTCAGTGGGGACTCAGCTCTTCAAACCAGACCTTATATTCCAGCTGGAGAGAGAAGAAAAGCGTTTGATGGTGGAGACCGAAACCCAGAAAGATGGGTGTTCAGCAAGCTTCAAAACGTCATATGTCAAATTTGCTGAAGGTGTTAACATCATAGATATTTTTACACTTAAAAATGTCGAATTCGTGCCAAAAAAAGAGCATTTGCGGGAAGTTTTAATTCATTACTTTATTTTGAAGAAAAGTGCTGCTGAAAGTTATCGTATACTTCGGGAAGCTTATGGTGAACATGCTCCATCTCAAGATACTTGTGAACGCTGGTTTAAACGCTTTAAAAGTGATGATTTCAATGTGAAAGACAAAGAACGTTCAGGTCAACCGAAAAAGTTTGAAGACCAACAATTACAAGCATTATTGGATGAAGATGCGGGTCAAACTCAAAAACAACTTGCAGAAAGATTAAATGTTGCTCAGCAAACAATTTCCGATCGTTTACAAGCAATGGGAAAGATTTTAAAGGAAGGAAAATGGGTGCCACATCAACTGAACGAAAGACAAATGGAAAACCGAAAAGTCATCAGTAAAATGTTGCTTCAACGGCATGAAAGAAAGTCTTTTTTGCATCGAATTGTGACTGGTGATGAAAAGTGGATTTATTTTGAGAATCCCAAATGCACAAAATCATGGGTTGATCCAGGTCAACCATCAACATCGACTGCAAGGCAAAATCGCTTCAGAAAGAAGACAATGCTCTGCGTTTGGTGGGATCAGGAAGGTGTGGTGTATTATGAGCTTCTAAAACCAGGTGAAACTGTTAATACTGATCGCTACCGACAACAGATAATCAGTTTGAACCATGCTTTGATTATGAAACGACCAGAACGGGCCAGAAGACAAGGCAAAGTAATTTTGCTTCATGATGATACACCATCACACACTTCAAAACCAGTTAAAGACACATTAAAAGATCTTGCCTGGGAAGTCTTAACCCACCCACCGTATTCACCAGACCTTGCTCCTTCAGATTACCACTTGTTCCGATCGATGGCACGCACACTTTCTCGGCAGCACTTCAAAACGTATGAAGAAGTGGAAAATTGGGTCTCTGAATGGTTTGCCTCAAAACAAGGAAAGTTCTATTGGGACGGTATCCACAAATTACCTGAAAGATGGGGGAAATGTGTAGCTAGCAATGGACATTACTTTGAATAAAGCATTTTTGATGTTTCTCTTGAAATTACCGTGTTTTCTTTGATTACAAAATCCGCATTATTAACCGGTACACCTGG
