## Supplemental Data 3 for "Recurrent evolution of vertebrate transcription factors by transposase capture"

>*Myotis_occultus_KTIGD5*_CDS

ATGGCTCCTAAGCGAAAGGCAGATTCTTCTGATGGCAGTTCTACGAAAAGAAGGAAAACCACAATGGAAGTGAAAGTAGATGTAATAAGACGCTCAGAAAAGGGCGAAACACCCAGACACATCGGCCGGTTGCTTGGCCTCAGTCCTACGACTGTCTCCACGATCATCCGCGATAAAAATCGTGCACTTGACCATGTGAAAGGATCTGTTCCTATGAAATCAACTGTGATTACAGAGCAACGCAGCGATCTCATGATTGAGATGGAAAGATTATTAGTGCTTTGGCTGGATGACCAGCATCAACGGCGTATCCCAGTTAGCCTGATGTTGATTCAGGAGCAGGCTAAAAGATTGTTTGAGGCCTTGAAAAAAGAAAAGGGGGAAGGGAGTGAAAGGGAAGAGTTTGTGGCTAGTAAAGGCTGGTTTATGTGTTTTAAGGCTCGTGCTAATTTGCATAATCTTAAGGTGCCATGGGAAGCTGCTAGTGGTGATGTGAAAGCAGCAGGTGATTTTCCTAGTGCGTTGGCTGAGATAATTAGGGAGGGGGGTTACTGTGATAAACAAGTGTTTAATGTGGATGAGACAGGCTTGTTCTGGAAGCCCATGCCTACACGTATGCACATTGCCAATGAGGAGAAGACAGCATCAGGCCATCAAGCCAGCAAGGAGAGACTGACTTTGCTACTTGGGGCTAATGCTGCTGGTGACTTCAAGCTGAAGCCCTTGGTGTGTCTGGCTGAGAATCCAAGGGCAACCAAGGGCATATGGAAGGGTCAACTACCTGTCATCTGGAAATCTAACAAGAAGGCCTGGGTGACACTTGGCGTATTTGACGACTGGTTCACCAACCATTTTGTGCCAGAAGTGAAGGACTACTGTGCTTCCAAGGGGCTTCCCTTTAAGGTGTTGCTAGTGCTGGGCAGTGCCCCTGGTCACCCTGCCGATCTGAATGACTTTCATCCAAATGTCAAGGTGGTTTACCTTCCACCTAACATCACCTCCCTTTTACAGCCTATGGACCAAGGTGTCATTGCTTTTTTCAAGGCCTACTACCTCCGAAGGACATTTGCTATGGCTTTTCGGGCAACTGAGGATAAAGAGTTGACTCTCAGTGACTTTTGGAAATCTTACAATGTTCTTGCTGCTGTAAAGAACATTTCTGATTCTTGGGTTGAAGTTAAGCAGACGAACCTGAATGGTGTTTGGAAAAAATTGTGTCCCCAGTTTGTGAATGATTTCCATGGCTTTGAGGATTCAGTTGAGGTTGTCATCCAGAATGTCGTTGAGCTGAGTAAGCAGCTTGATTTGGAGGTGGAGGCTGAGGATGTCACAGAGCTGTTGGCATCTCATGGAGAGGAGCTATCTACCCAGGACCTCATTCAAGTGAAACAGCAGTTAATTGAAGAGGAAGACACACCAACCCCTGAGCCCAAGAGATTTACAAGCAAGGAACTGGCAGGGGCCTTTGCCATGATAGAGGATGCCTTGGCAAGGTTTGAGGCTCAGGACCCCAACATTGACAGGTACACTAAGGTTGCCAGAGGGGTCATGGATAGCCTGCAGTGTTACAAGGAGATTTGGGAGGACAAGAAGAAGGCATCCTTCCAGACTAGCCTTGAACATTATTTTAAGAAGGTAGAGAGGCCTGCAACAGATCCTGTACCCTCTACCTCATACGCCTCTCCAGACTCTCCTGACCCAGTATCACATGCACCTTCTGCAGGTCCTGCCTCTCCAGACTCACCTGACCCAGTATCTCCAGCACCTTCTGCAGGATCTGCCTCTCCAGACTCACCTGACCCAATATCTCCAGCACCTTCTGCAGGTGCTGCCTCTCAAAACTCCTCCCCCCAGCAGCCTAGTGTAAGCCTGTCTTTCCCTCCTTTACAAAGTCCAGTGTGCAAGCCAACAACAATAGTAAAGGATATGATCTTTGAGGACGTGGCCATTGCCTTCTCTCAGGAGGAGTGGGAGCTCCTTGATGAAGCTCAGCGACGTCTGTACTGTGATGTGATGCTGGAAGTGTTTGCACTTCTATCAATTGTAGGGGGGGTGACTATATCCAGATCCATGGCCTCCACTTCTGCACCTTTCCTGTAG

>*Myotis_lucifugus_KTIGD5_*CDS

ATGGCTCCTAAGCGAAAGGCAGATTCTTCTGATGGCAGTTCTACGAAAAGAAGGAAAACCACAATGGAAGTGAAAGTAGATGTAATAAGACGCTCAGAAAAGGGCGAAACACCCAGACACATCGGCCGGTTGCTTGGCCTCAGTCCTACGACTGTCTCCACGATCATCCGCGATAAAAATCGTGCACTTGACCATGTGAAAGGATCTGTTCCTATGAAATCAACTGTGATTACAGAGCAACGCAGgGATCTCATGATTGAGATGGAAAGATTATTAGTGCTTTGGCTGGATGACCAGCATCAACGGCGTATCCCAGTTAGCCTGATGTTGATTCAGGAGCAGGCTAAAAGATTGTTTGAGGCCTTGAAAAAAGAAAAGGGGGAAGGGAGTGAAAGGGAAGAGTTTGTGGCTAGTAAAGGCTGGTTTATGTGTTTTAAGGCTCGTGCTAATTTGCATAATCTTAAGGTGCCATGGGAAGCTGCTAGTGGTGATGTGAAAGCAGCAGGTGATTTTCCTAGTGCGaTGGCTGAGATAATTAGGGAGGGGGGTTACTGTGATAAACAAGTGTTTAATGTGGATGAGACAGGCTTGTTCTGGAAGCCCATGCCTACACGTATGCACATTGCCAATGAGGAGAAGACAGCATCAGGCCATCAAGCCAGCAAGGAGAGACTGACTTTGCTACTTGGGGCTAATGCTGCTGGTGACTTCAAGCTGAAGCCCTTGGTGTGTCTGGCTGAGAATCCAAGGGCAACCAAGGGCATATGGAAGGGTCAACTACCTGTCATCTGGAAATCTAACAAGAAGGCCTGGGTGACACTTGGCGTATTTGAGGACTGGTTCACCAACCATTTTGTGCCAGAAGTGAAGGACTACTGTGCTTCCAAGGGGCTTCCCTTTAAGGTGTTGCTAGTGCTGGGCAGTGCCCCTGGTCACCCTGCCGATCTGAATGACTTTCATCCAAATGTCAAGGTGGTTTACCTTCCACCTAACATCACCTCCCTTTTACAGCCTATGGACCAAGGTGTCATTGCTTTTTTCAAGGCCTACTACCTCCGAAGGACATTTGCTATGGCTTTTCGGGCAACTGAGGATAAAGAGTTGACTCTCAGTGACTTTTGGAAATCTTACAATGTTCTTGCTGCTGTAAAGAACATTTCTGATTCTTGGGTTGAAGTTAAGCAGACGAACCTGAATGGTGTTTGGAAAAAATTGTGTCCCCAGTTTGTGAATGATTTCCATGGCTTTGAGGATTCAGTTGAGGTTGTCATCCAGAATGTCGTTGAGCTGAGTAAGCAGCTTGATTTGGAGGTGGAGGCTGAGGATGTCACAGAGCTGTTGGCATCTCATGGAGAGGAGCTATCTACCCAGGACCTCATTCAAGTGAAACAGCAGTTAATTGAAGAGGAAGACACACCAACCCCaGAGCCCAAGAGATTTACAAGCAAGGAACTGGCAGGGGCCTTTGCCATGATAGAGGATGCCTTGGCAAGGTTTGAGGCTCAGGACCCCAACATTGACAGGTACACTAAGGTTGCCAGAGGGGTCATGGATAGCCTGCAGTGTTACAAGGAGATTTGGGAGGACAAGAAGAAGGCATCCTTCCAGACTAGCCTTGAACATTATTTTAAGAAGGTAGAGAGGCCTGCAACAGATCCTGTACCCTCTACCTCATACGCCTCTCCAGACTCTCCTGACCCAGTATCACATGCACCTTCTGCAGGTCCTGCCTCTCCAGACTCACCTGACCCAGTATCTCCAGCACCTTCTGCAGGATCTGCCTCTCCAGACTCACCTGACCCAATATCTCCAGCACCTTCTGCAGGTGCTGCCTCTCAAAACTCCTCCCCTCAGCAGCCTAGTGTAAGCCTGTCTTTCCCTCCTTTACAAAGTCCAGTGTGCAAGCCAACAACAATAGTAAAGGATATGATCTTTGAGGACGTGGCCATTGCCTTCTCTCAGGAGGAGTGGGAGCTCCTTGATGAgGCTCAGCGACGTCTGTACTGTGATGTGATGCTGGAAGTGTTTGCACTTCTATCAATTGTAGGGGGGGTGACTATATCCAGATCCATGGCCTCCACTTCTGCACCTTTCCTGTAG

>*Eptesicus_fuscus_KTIGD5*

ATGGCTTCTAAGCATAAGGCAGATTCTTCTTCTGGCAGTGCTACAAAAAGAAGGAAAGCCCTCACAATGGAAGTGAAAGGAGACATAATAAGACGCTCAGAAAAGGGCAAAACACCTACAAACATCAGCCGGTTGCTTGGCCAAAGTCCTACAACTGTCTCGACAGTCATCGACAATCAAAAGCGTATACTTGAACGTGTGAAAGGATCTGCTCCTATGAAATCAACTGTGGTTACAGAGCAGTGGAGTGGTCTCATTATTGAGATGGAAAGATTATTAGTGCTTTGGCTGGATGAGCAGCATCGATGGCGTATCCCAGTTAGCCAGATGTTGATTCAGGAGAAGGCTAAAAGATTGTTTGAGGCCTTGAAAAAAGAAAAGAGGGAAGGGAGTGAAAGTGAAGAGTTTGTGGCTAGCAAAGGTTGGTTTACACGTTTTAAGGCTCGTGCTAATTTGCATAATCTTAAGGTGCCAGGGGAAGCTGCTAGTGGTGATGCCAAAGCAGCAGGTGATTTTCCTAGTGCATTGGCTGAGATAATTAGGGACGGGGGTTACTGTGATCAACAAGTGTTTCATGTGCATGAGACAGGCTTGTTCTGGAAGCATATGCCTATACGTAAGTACATTGCCAAGGAGGAGAAGATAGCATCAGGCCATAAAGACAGCAAGGAGAGATTGACTTTGCTACTTGGGGCAAATGCTGCTTGTGATTTCAAGCTGAAGCCCTTGGTGTGTCTGGCCGAGAATTCAAGGGCAATCAAGGGCATATGGAAGGGTCTACTGCCTGTCATCTGGAAATCTAACAAGAAGGCCTGGGTGACACTCAGCGTATTTGAGGACTGGTTCACCAACAATTTTGTGCCAGAAGTCAAGGACTACTGTGCTTCAAAGGGGCTGCCCTTTAAGGTGTTGCTAGTGCTGGACAGTGCCCCTTGTCACCCTGCCGATTTGAATGACTTTCACCCAAATGTCAAGGTGGTTTTCCTTCCACCTAGCACCCCCTCCCTTTTACAGCCTATGGGCCAAGGTGTCATTGCCTCTTTCAAGGCCTACTACCTCCGAAGGACATTTGCTATGGCTTTTAGGGCAACAGAGAAGGATAAAGAGTTGACTCTCAATGACTTTTGGAAATCTTACAATGTTCTTGATGCTGTAAAAATCATTTCTGATTCTTGGGATGAAATTAAGCAGAGGAACTTGAATGATGTTTGGAAAAAATTGTGTCCCCAGTTTGTGAACGATTTCCAGGGCTTTGAGAATTCAGTTGAGGTTGTCATCAAGAATGTTGTTGAATTGAGTAAGCAGCTCGATTTGGAGGTGGAGGCTGAGGATGTCACAGAGCTGCTGGCATCTCATGGAGAGGAGCTATCTGCCGAGGACCTCACTCAACTGAAACAGCAGTTCATTGAAGAGGAAGACACACCAACCCCAGAGCCCAGGAGATTTACAAGCAAGGAATTAGCAGGGGCCTTTGCCATGATAGAGGAAGCCTTGGAAAGGTTTGAGGCTCAGGACTCCAACAGTGACAGGTACACTAAGGTTGCCAGAGGGGTCATGGATAGCCTGCAGTGTTACAAGGAGATTTGGGAGTACAAGAAGAAGGTATCCTTCCAGACTACCCTTGAACATTATTTTAAGAAGGTGGAGAGGCCTGCAACATATCCCGTACCCTCTACCTCATGTGCTTCTCCAGACTCACCTGACCCAGTATCTCCAGCACATTCTGCACGTTCTGCCTCTCCGGACTCACCTGACCCAGGATCTCCAGCACCTTCTGCAGGTTCTGCCTCTCCGGACTCACCTGACCTAGTAGCACATGCATCTTCTGCAGGTTCTGCCTCTCAAACCTCCTCCGCACAGCAGCCTAGTGTAAGCCTGTCTTTCCCCACTTTGCAATGTCTTGTGTGCAGGCCAACATCAGTACTAAATATGAACTTTGAGGACGTGGCCATTGCCTTCTCTCAGGAGGAATGGGAGATCCTTGATGAGACTCAGAGACGTCTTTACTGTGATGTGATGCATGAAGTGTTTGCACTTGTATCATTCATAGGTAAGACCTTCACATCTCTCCTAGTGTCCTGAGGTGA
